# A voltage-step method for detecting high-frequency transient current components in deep brain tissue: preliminary in vivo measurements in rats

**DOI:** 10.64898/2026.07.03.736373

**Authors:** Sultan Mahmud, Domenica Baez, Anyu Jiang, Yitong Zhao, Baibhab Chatterjee, Adam Khalifa, Chris Rourk

**Affiliations:** College of Engineering, Dept. of Electrical and Computer Engineering, University of Florida; Citizen Scientist, formerly with U.S. Nuclear Regulatory Commission, Office of Nuclear Regulatory Research; Virginia Tech

## Abstract

A test technique for measuring high-frequency transient current components in deep brain tissue is presented. The technique applies a voltage pulse with a high value in dV/dt, generating a corresponding current pulse with high dI/dt that can elicit measurable transient current responses from the electrode–tissue interface and adjacent brain tissue; responses are analyzed in the frequency domain by Fast Fourier Transform at a 200 kHz sampling frequency. The method was motivated by prior evidence that ferritin and neuromelanin in catecholaminergic tissue may support high-frequency conduction properties that have not previously been characterized in vivo. The protocol was applied in 277 measurements across five Sprague Dawley rats at cortical and basal ganglia locations in different locations in the brain. Preliminary spectral results show differences between catecholaminergic regions and cortical tissue that support further development and validation of the method.

## Introduction

Ferritin and neuromelanin are biophysical materials with unusual electrical properties.^i, ii^ They are present at elevated concentrations and quasi-ordered structures in the large dopamine neurons of the *substantia nigra pars compacta* (SNc), the ventral tegmental area (VTA), the *zona incerta* (ZI) and other dopaminergic neurons, and also in the large noradrenaline neurons of the *locus coeruleus* (LC), which produce dopamine as a precursor to noradrenaline.^iii, iv^ The level of ferritin and neuromelanin in those tissues is high enough to provide endogenous magnetic resonance imaging contrast.^v, vi^ In addition, it has recently been shown that the soma of these neurons are connected together by processes of tissue-resident oligodendrocyte precursor cells (OPCs) that contain high concentrations of ferritin.^vii^

Testing the electrical properties of the SNc tissue *in vivo* is technically challenging. While deep brain stimulation (DBS) studies have been performed, those are typically limited to specific low-frequency waveforms and are used to evaluate the systemic response to DBS.^viii^ Electrochemical studies have also been performed, but those are limited to the detection and evaluation of the electrochemical properties of dopamine *in vivo* as opposed to the electrical properties of the tissues.^ix^ Tests to determine the tissue response to high-frequency electrical stimulation in the brain are typically performed using sinusoidal waveforms and measuring root-mean square (RMS) parameters,^x^ which obscures high frequency transients above 10 kHz, where electron tunneling could be occurring. Also, conventional EIS does not utilize fast voltage steps to extract time-domain transient structures.

The CAFM tests of SNc tissue measured transient currents that were possibly at frequencies greater than 10 kHz, based on the CAFM probe scan rate and contact behavior. In order to further investigate whether the currents that were observed in those tests could also be observed *in vivo*, a novel high frequency testing protocol was developed. Methodologically, the high frequency test protocol exposes the tissue to a large current transient (dI/dt) and voltage transient (dV/dt) excitation pulse and then measures the frequency components of the currents that follow, to determine whether the excitation pulse resulted in high frequency currents. This high frequency protocol was used to measure currents in the cortical and basal ganglia tissues of Sprague Dawley rats, and the temporal current signals were analyzed in the frequency domain to determine whether evidence of transient currents was present in dopaminergic neuron tissue.

## Methods

Male Sprague Dawley rats (100–116 days old, 380–480 g) were used for this study. Animals were anesthetized with isoflurane, with induction performed at 3.5% and anesthesia maintained between 1.8–2.8% throughout the experiment. Body temperature was maintained using a heating pad during all surgical and experimental procedures. All procedures were conducted in accordance with approved ethical guidelines and institutional animal care protocols by Institutional Animal Care and Use Committee (IACUC) at the University of Florida.

Animals were secured in a stereotaxic frame (Kopf Instruments), and burr holes were drilled using bregma as the stereotaxic reference point. Target coordinates for the substantia nigra pars compacta (SNc) were initially selected using the Siibra Atlas and were subsequently refined based on histological validation results. Electrophysiological testing was performed on the right and left hemispheres when experimental time permitted. Stimulation was delivered using an SNE-100 Concentric Bipolar Electrode concentric bipolar electrode (Micro Probes). To enable postmortem localization of the electrode tract, the electrode tip was coated with the fluorescent tracer DiI prior to insertion.

Following completion of the experiments, animals were euthanized and the brains were extracted and fixed in 4% paraformaldehyde (PFA). Fixed brains were sectioned and imaged using fluorescence microscopy to confirm electrode placement and assess the anatomical location of the stimulation sites as shown in Figure 1.

**Figure 1.**
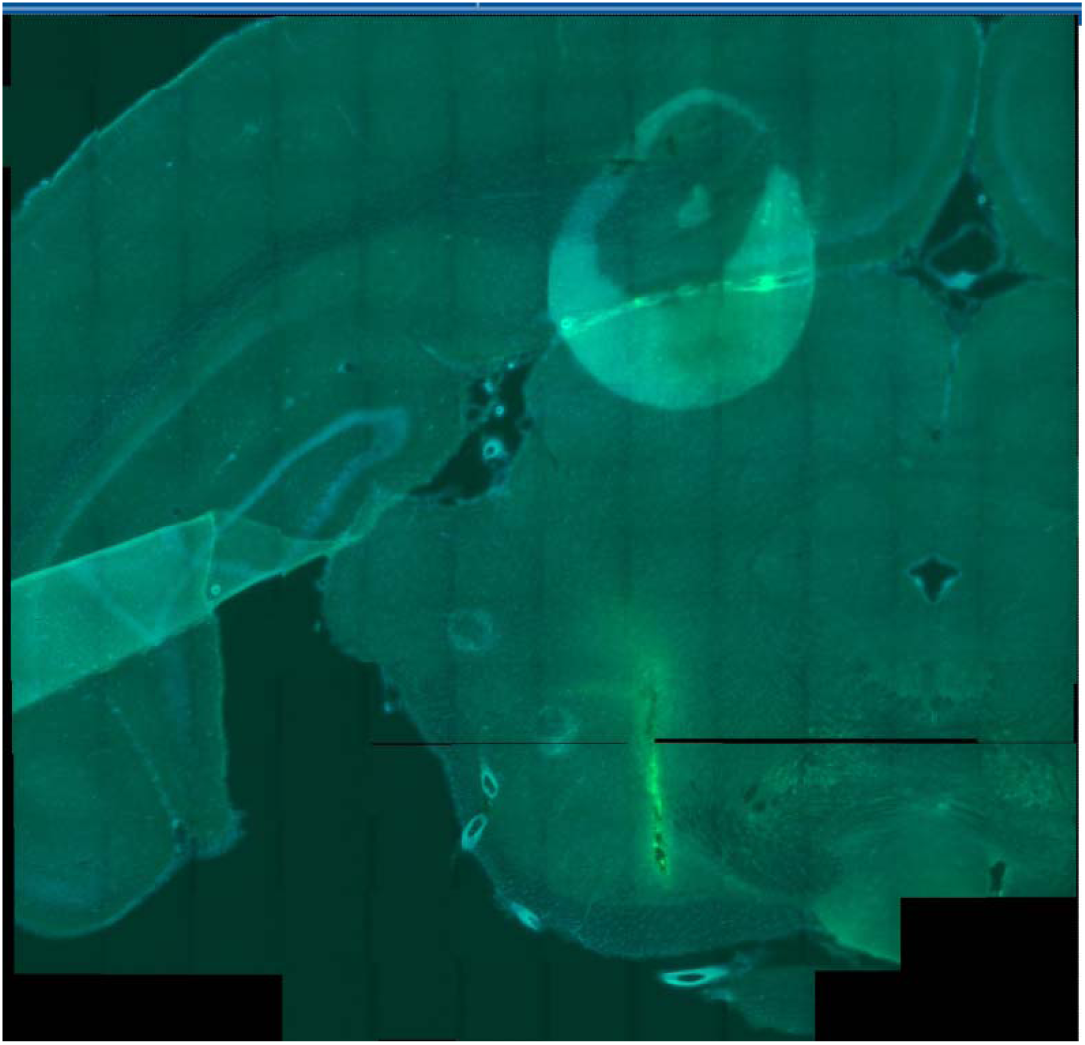
Electrode tract localization in the substantia nigra pars compacta (SNc). Representative coronal section from a rat brain showing the DiI-labeled electrode tract targeting the SNc.

A concentric electrode was used for the method development (Figure 2a). The optical image confirms the fabricated electrode dimensions, including an exposed electrode region approximately 160 µm in length, 75 µm in width, and a 50 µm sidewall/contact feature. These dimensions define the active electrode area that interacts with the electrolyte during electrochemical testing.

**Figure 2.**
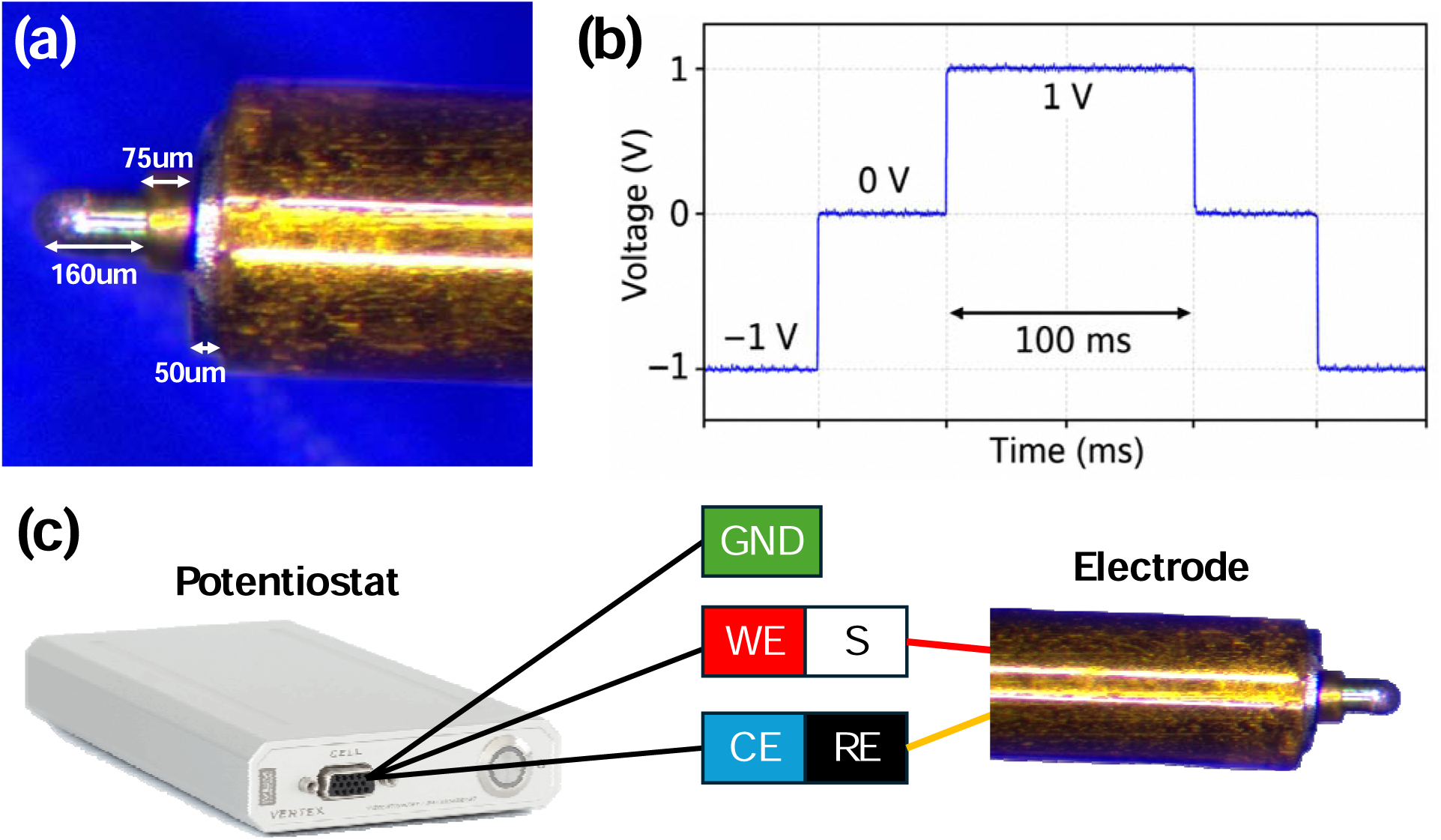
Experimental electrode and electrochemical measurement setup, (a) Optical image of the fabricated electrode showing the approximate electrode dimensions, (b) Applied step-voltage waveform used during the experiment, and (c) Schematic illustration of the potentiostat-based electrochemical impedance spectroscopy setup.

Electrical characterization included a novel voltage-step protocol as shown in Figure 2(b) in addition to standard electrochemical impedance spectroscopy (EIS), performed at 0.5 V across 50 frequencies from 0 to 10,000 Hz. The voltage-step sequence was −1 V to 0 V, 0 V to 1 V, 1 V to 0 V, and 0 V to −1 V. The current was measured at each step transition using a 5 µs sampling period, corresponding to a sampling frequency of 200 kHz. The measured current response was then analyzed in the frequency domain using a Fast Fourier Transform (FFT) in Python and JupyterLite to identify the frequency components (see Appendix I). In general, two voltage-step tests were performed first, followed by an EIS measurement and then two additional voltage-step tests, for a total of five tests at each depth. An Ivium potentiostat was used for all electrical measurements.

For EIS, a two-electrode configuration was used. The working electrode (WE) and sense (S) leads were connected to one electrode, while the reference electrode (RE) and counter electrode (CE) leads were connected to the second electrode. Therefore, the measured impedance represented the total electrode–tissue–electrode pathway, as illustrated in Figure 2(c). Measurements were also performed in saline and in an agarose brain phantom to verify the proper function of the experimental setup, and are shown in Appendix I side-by-side with a scan from the SNc. The Ivium potentiostat was configured with automatic filtering, 16-bit DC offset subtraction, and two DC decoupling filters. During in vivo measurements, the rats were grounded and shielded using a Faraday cage formed from a conductive blanket draped around the rat and the test equipment.

## Results

Table 1 presents the summary of the pilot study tests that were performed on 5 different rats, each on a different date. A total of 277 step voltage tests and 75 EIS tests were performed. Example step voltage test results are shown in Figures 3-7, and a sample EIS test result is shown in Figure 8. Additional results are provided in the supplemental materials, and the raw data is available in the data archive. The tests were performed across 5 animals, and most target conditions represented by a single animal and a single session. A data analysis is provided in Appendix 2 that shows a cosine similarity metric and root mean square error difference metric between pairs of FFT scans, but a statistical analysis that includes means, ranges, variability measures and other textbook-type statistics would be meaningless, and was not performed. A distribution of measurements would not provide any meaningful information and was not provided.

**Figure 3.**
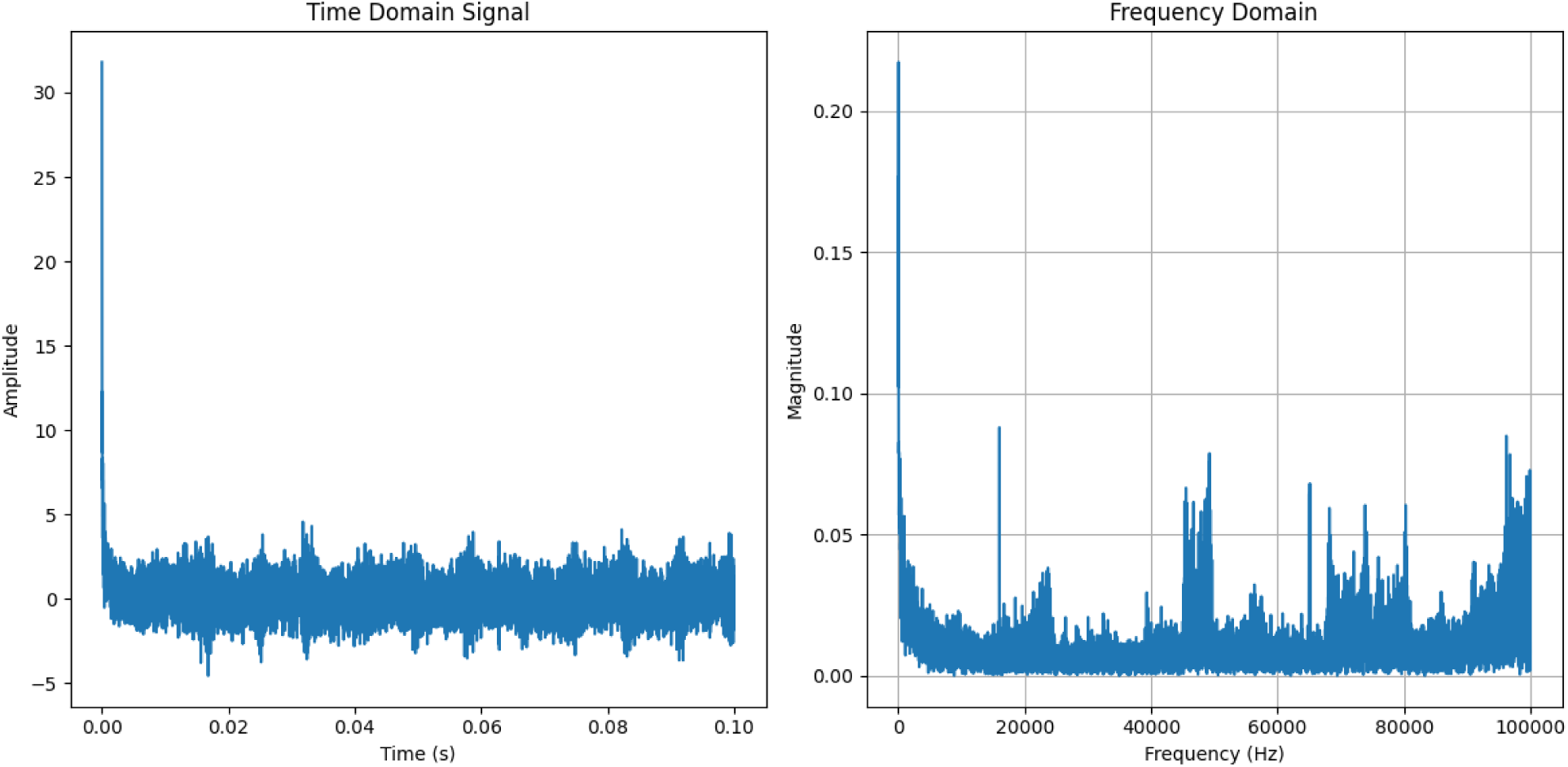
Signal data measured at ML = 1.304, AP = –3.311 Z = 8.6 mm, which corresponds to the zona incerta. Significant magnitudes of high frequency content are present from 0 to 100 kHz, with banding apparent at ∼15 to 25 kHz, 39 to 50 kHz and 65 to 80 kHz and 90 to 100 kHz.

**Figure 4.**
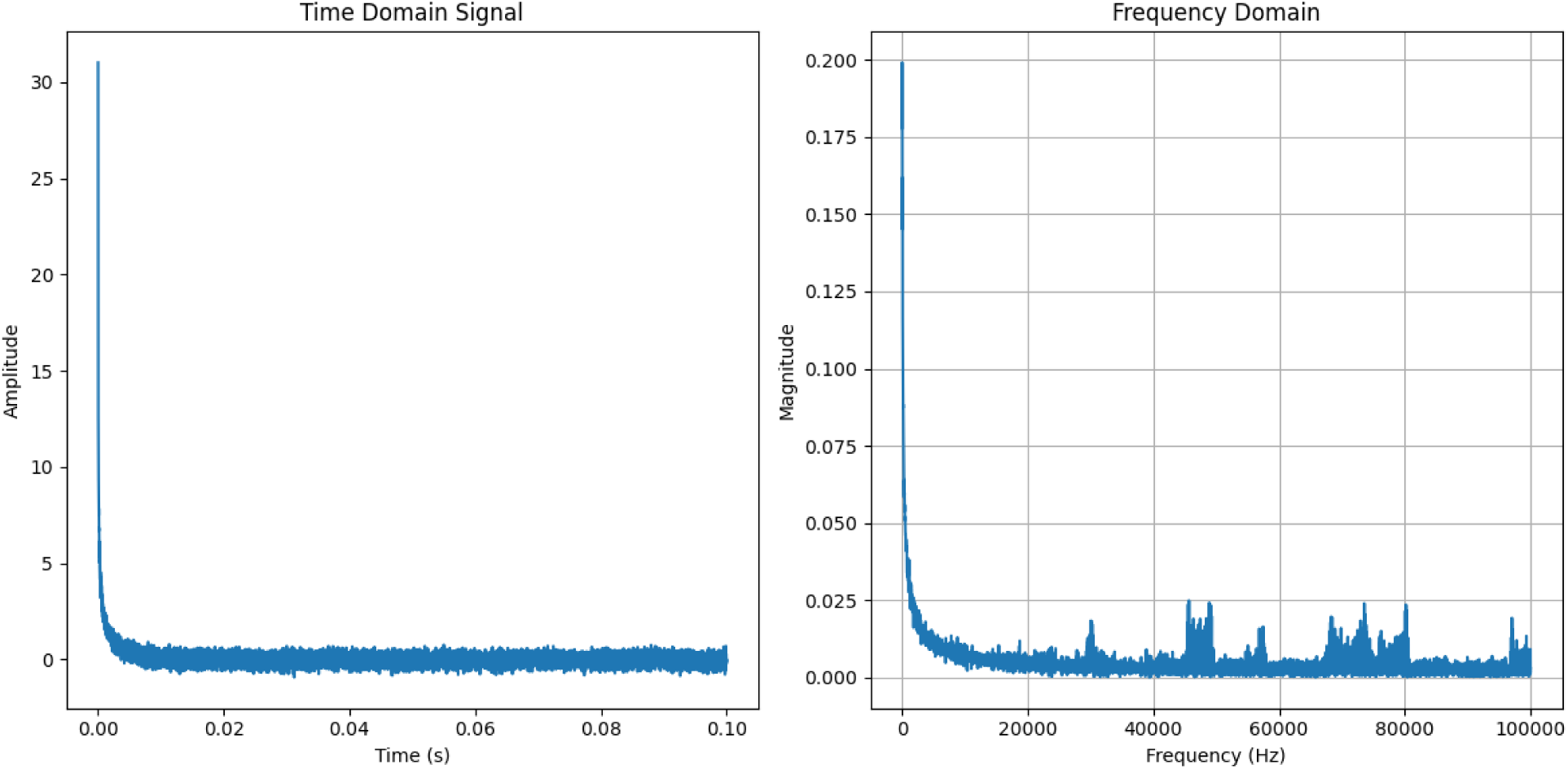
ML = Signal data measured at 1.537, AP = –4.532 Z = 8.8 mm, which corresponds to the lateral hypothalamus. Significant magnitudes of high frequency content are present from 0 to 100 kHz, with banding apparent at ∼45 to 50 kHz, 65 to 80 kHz and 95 to 100 kHz.

**Figure 5.**
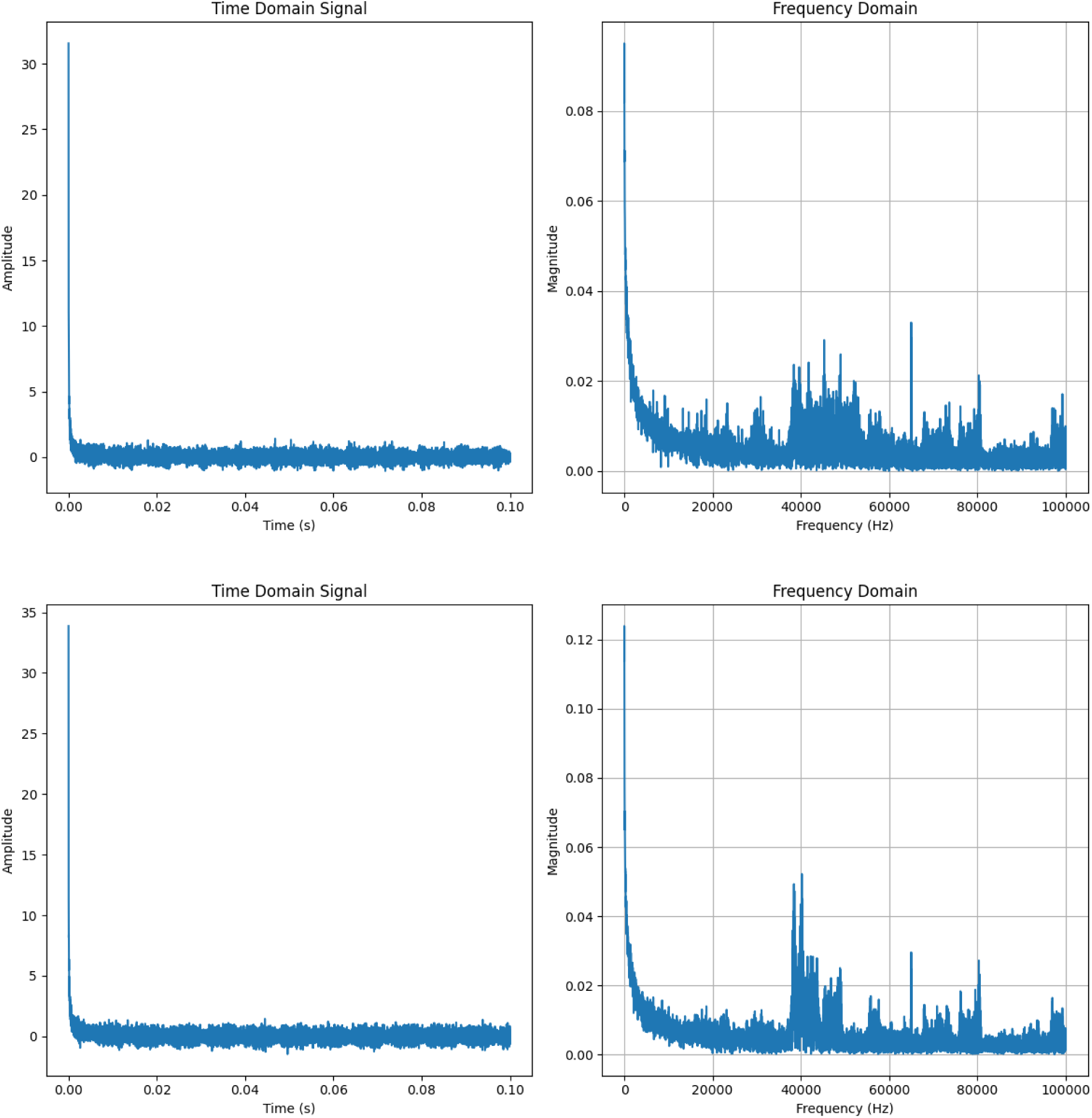
Signal data measured at ML = 2.162, AP = –5.235 and Z = 7.6 mm (top) and 7.7 mm (bottom), which corresponds to the substantia nigra pars compacta. Significant magnitudes of high frequency content are present from 0 to 100 kHz, with banding apparent at ∼38 to 55 kHz, 58 to 80 kHz and 95 to 100 kHz at 7.6 mm and from ∼38 to 43 kHz, 45 to 50 kHz, 55 to 58 kHz, 65 to 72 kHz, 75 to 80 kHz and 95 to 100 kHz at 7.7 mm. These two data sets are an example of the significant differences that were observed for some regions with minor changes in electrode depth

**Figure 6.**
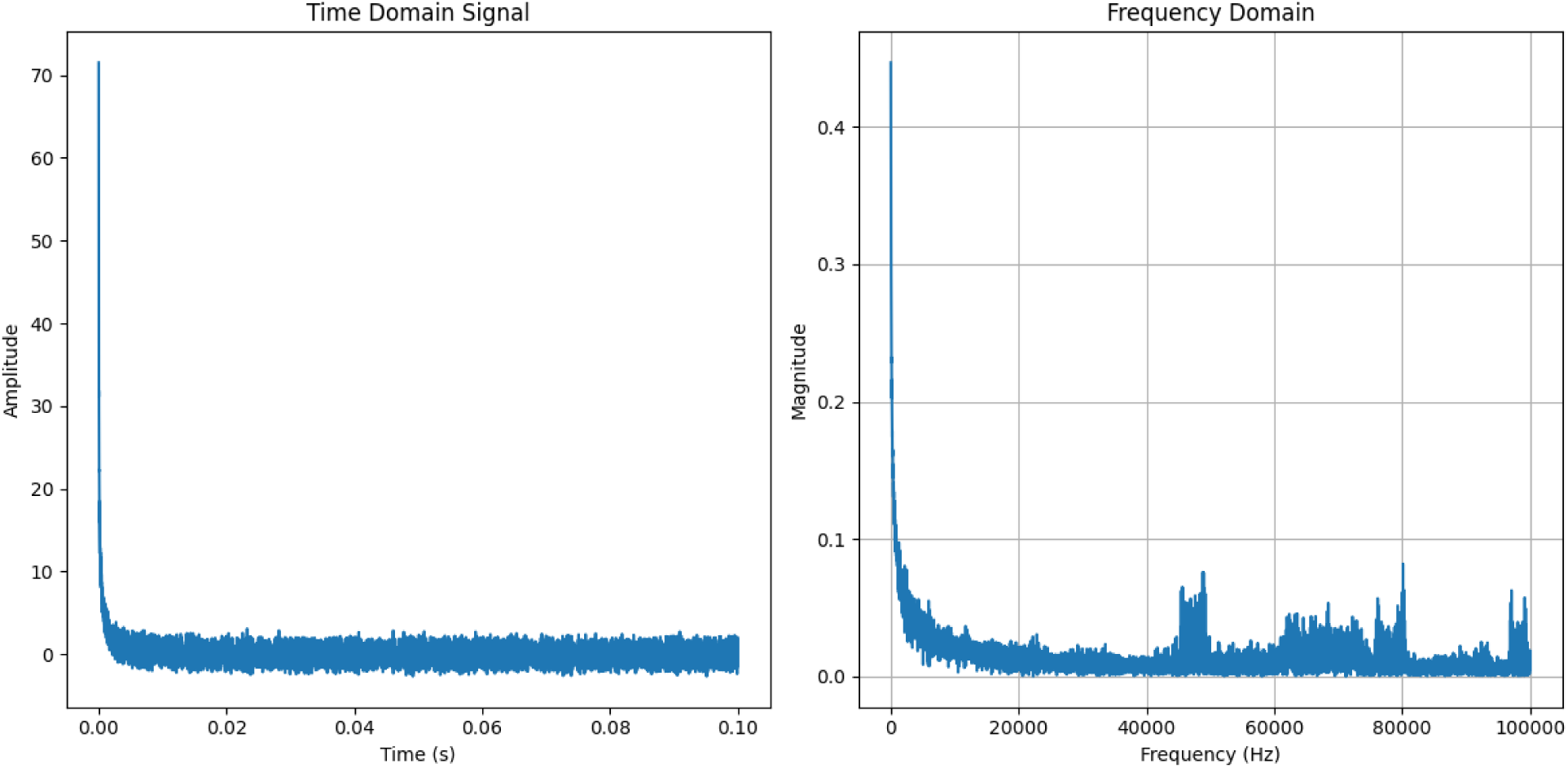
Signal data measured at ML = 2.621, AP = –5.513, Z = 7.0 mm, which corresponds to the substantia nigra pars compacta. Significant magnitudes of high frequency content are presen from 0 to 100 kHz, with banding apparent at ∼45 to 50 kHz, 60 to 80 kHz and 95 to 100 kHz.

**Figure 7.**
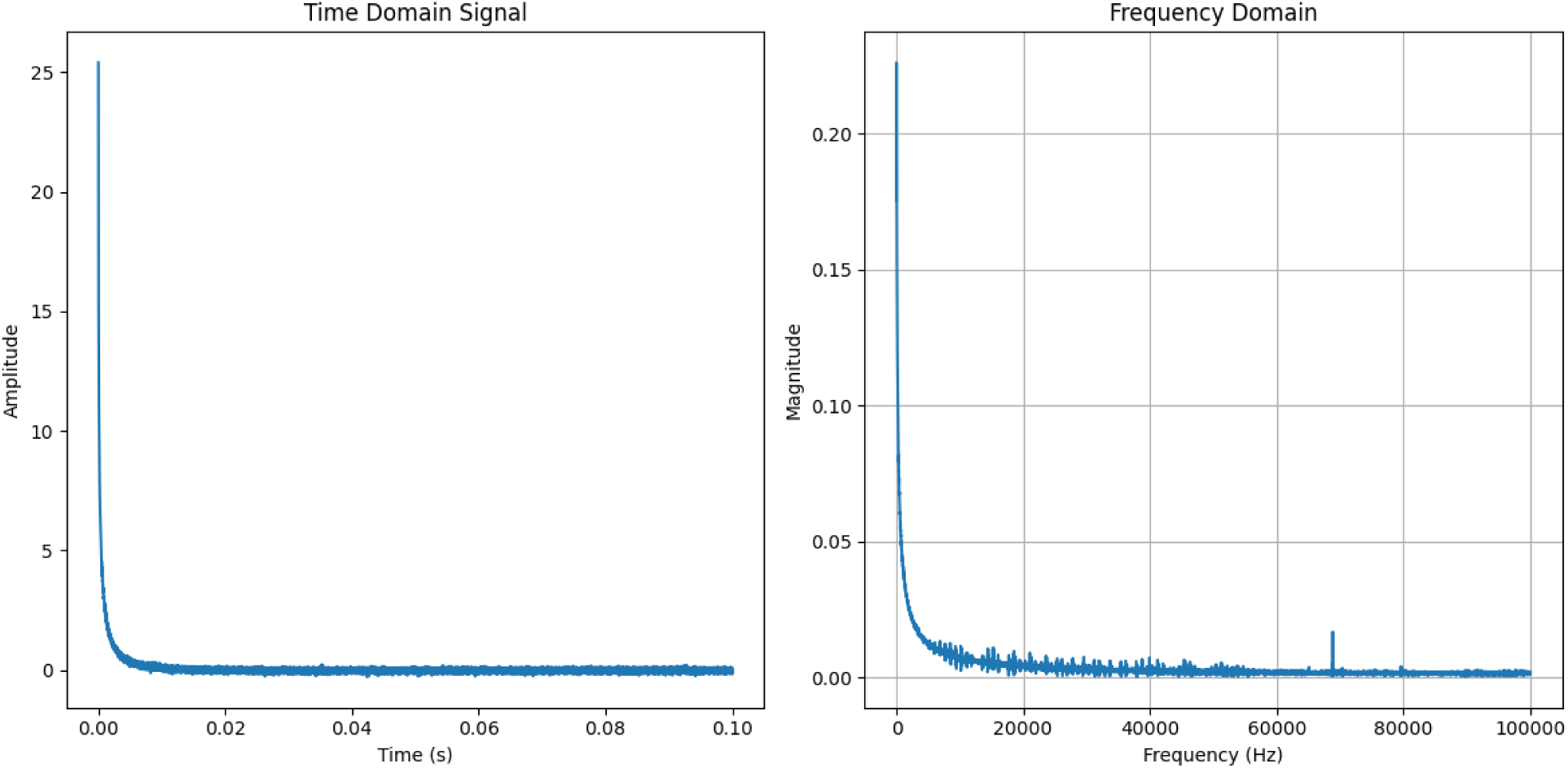
Signal data measured at ML = –2.15, AP = –6.194, Z = 5 mm, which corresponds to the cortex. The low-magnitude individual spikes in the data are present from 0 to ∼50 kHz, and include positive and negative spikes that are characteristic of Gaussian-type noise.

**Figure 8.**
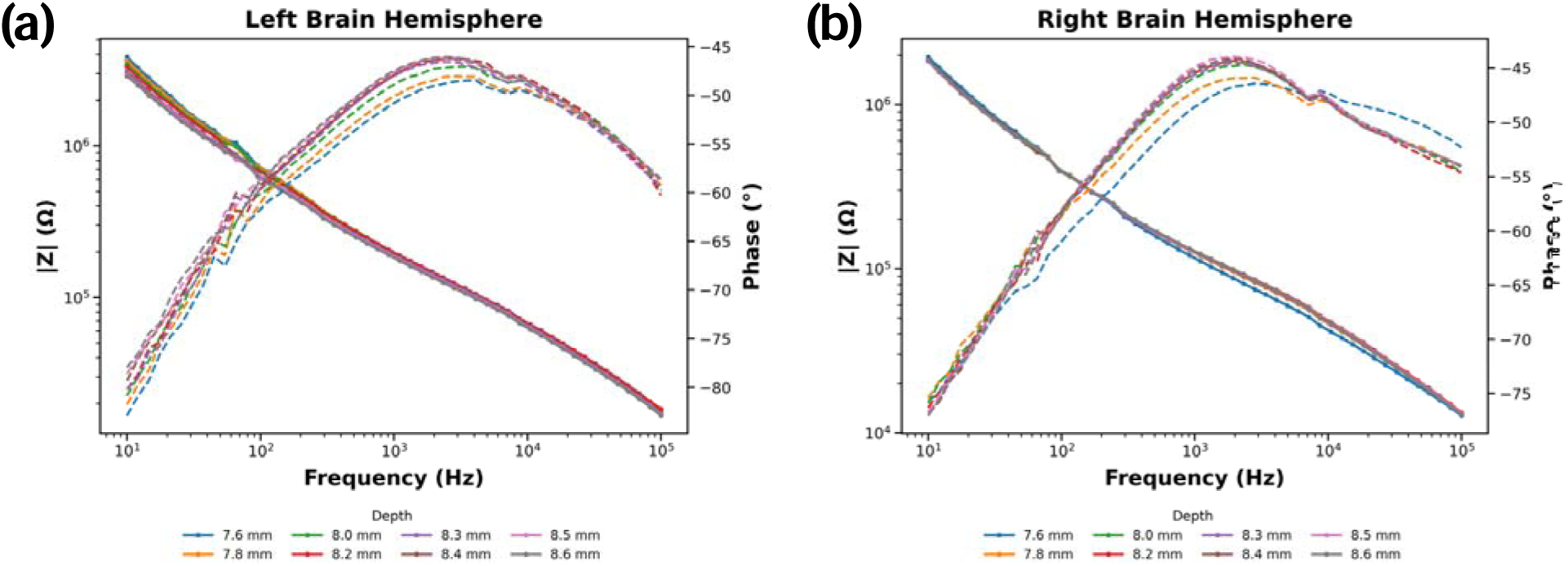
Depth-dependent in vivo EIS response of the left and right brain hemispheres. Bode plots showing and phase versus frequency at insertion depths from 7.6–8.6 mm.

**Figure 9.**
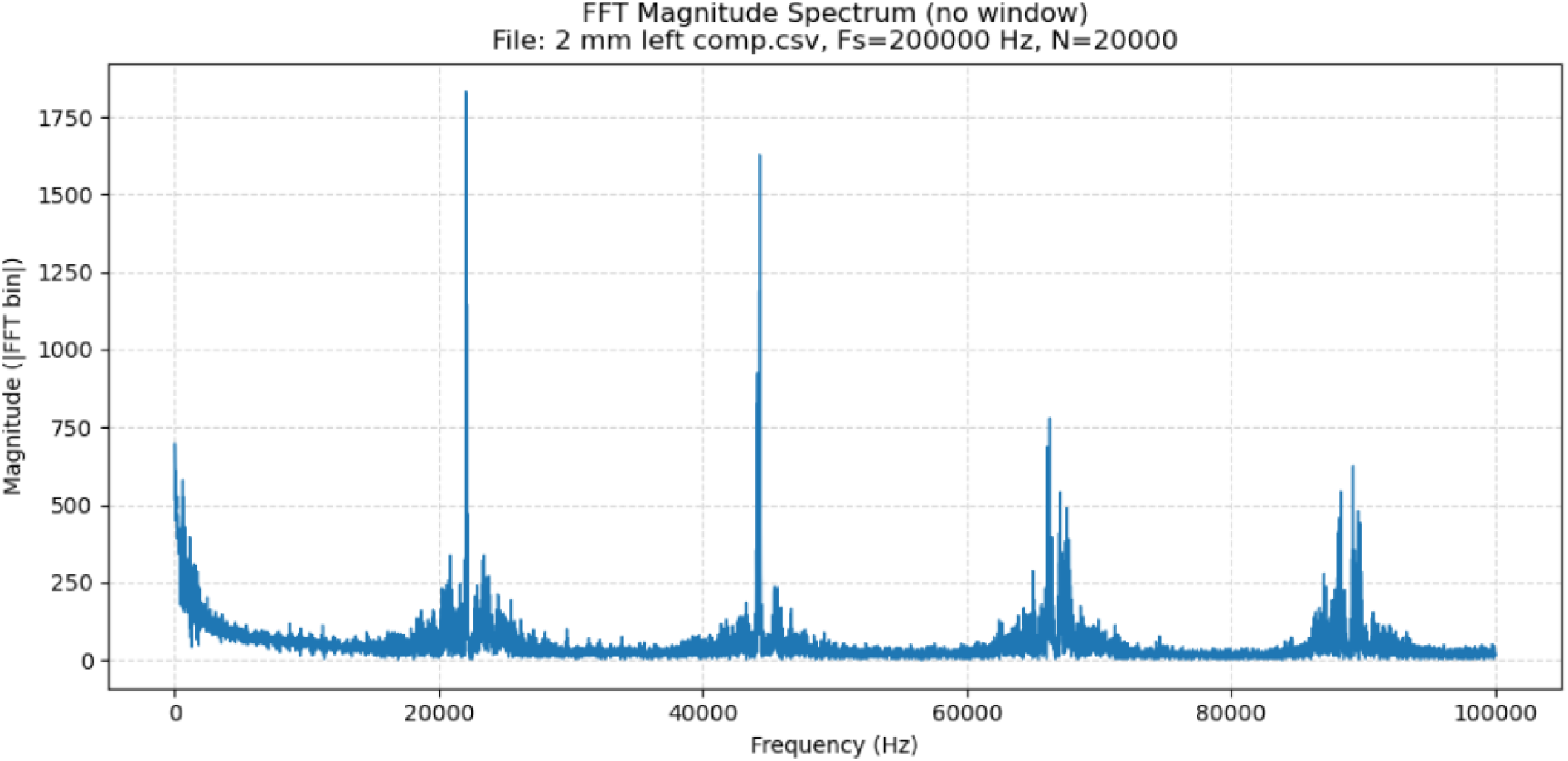
A typical noise signal. The high magnitude spikes are at harmonic multiples and include side bands that are indicative of frequency modulation.

**Table 1.** Summary of tests.

| Test,<br>figure | location | Coordinates (ML,<br>AP, DV) | di/dt<br>measurements | EIS<br>measurements |
| --- | --- | --- | --- | --- |
| 1, Fig. 3 | zona incerta | 1.304, -3.311, 7.6-8.6 (left only)<br>7.6, 7.8, 8.0, 8.2,<br>8.3, 8.4, 8.5, 8.6 | 32 | 8 |
| 2, Fig. 4 | Lateral<br>hypothalamus<br>(left)<br>Subthalamic<br>nucleus<br>(right) | 1.537, -4.532, 8.2-8.8 (L)<br>-2.268, -3.965, 8.2-8.8<br>(R)<br>8.2, 8.3, 8.4, 8.5,<br>8.6. 8.7. 8.8 (both) | 56 | 14 |
| 3, Fig. 5 | SNc | 2.16, +/-5.23, 7.6 –<br>8.1<br>7.6, 7.7, 7.8, 7.9. 8,<br>8.1 | 48 | 12 |
| 4, Fig. 6 | SNc (left)<br>VTA-SNc<br>(right) | 2.62, -5.51, 7.0–8.0<br>(L)<br>-1.84, -5.66, 7.0-8.0<br>(R)<br>7.1, 7.2, 7.3, 7.4,<br>7.5, 7.6, 7.7, 7.8,<br>7.9, 8.0 | 68 | 22 |
| 5, Fig. 7 | Cortex | 2.15, -6.194<br>2, 2.5, 3, 3.5, 4, 4.5,<br>5, 5.5, 6 (both)<br>6.5, 7 (right only) | 73 | 19 |

## Discussion

A step voltage excitation was chosen in order to generate a high dV/dt and dI/dt state in the tissue adjacent to the probe tip. The 200 kHz sampling frequency provided a maximum frequency detection of 100 kHz using the Fourier transform, consistent with the Nyquist limit. The specific values of dV/dt and dI/dt for each test can be obtained from the test data, but were typically greater than 200 kV/sec and 4 amperes/second.

In cortical tissue, it was expected that this type of excitation would not result in any high frequency components, because of the electrical impedance characteristics of cell membranes at high frequencies. In contrast, high frequency currents were expected in the dopaminergic neurons of the SNc, which have elevated levels of ferritin and neuromelanin that are arranged in structures that are capable of supporting sequential electron tunneling. The CAFM tests on fixed tissue that were previously conducted were not sufficiently accurate to provide an estimate of the frequency components of the currents that were measured, but there was evidence of the presence of current transients during the peak contact period. The CAFM oscillation period for the current measurements was 200 Hz, and the peak contact period is estimated to encompass 1% of the CAFM sampling oscillation period, which yields an effective transient frequency detection limit of ∼20 kHz.

As can be seen in Figure 3, high frequency currents were measured in an area that is believed from the histology data to have been the zona incerta, a dopaminergic neuron cluster near the SNc. Targeting the SNc is complicated by numerous factors, such as the variability of skull shapes and brain sizes in rats, indexing tolerances and so forth. However, the levels of ferritin and neuromelanin in the zona incerta are similar to those in the SNc, and high frequency current components were observed as shown. Current paths of the currents were generated in response to the step change from –1 V to 0 V appear to have been generated in that tissue. Referring to Table 1, 32 voltage step protocol tests were performed on the left hemisphere side only. As discussed further in the Appendices and the supplemental materials, the test results at each test location generated results at each depth that had high cosine similarity metrics and low root mean square error metrics, before and after the EIS tests, which is referred to herein as being “repeatable.” The measured currents changed slightly at each depth. No unusual noise or transients were observed at any tests. While only a single set of data is shown, the currents measured in the zona incerta were generally consistent in terms of frequency and magnitude, as can be seen in the data in Appendix 1 and the supplemental materials. The magnitudes of the measured frequency components varied from test to test but did not change frequency bands.

Based on the histology data, the results in Figure 4 are believed to be from the lateral hypothalamus and subthalamic nucleus, both of which have elevated levels of ferritin but which lack dopaminergic neurons. In general, it is difficult to map ferritin in neural tissue, and that has not been done largely because its unusual electrical properties are generally recognized. Further testing should include mapping of ferritin and identification of the types of cells that the ferritin can be found in. Referring to Table 1, 56 voltage step protocol tests were performed on the left and right hemispheres. As discussed further in the supplemental materials, the test results were repeatable at each depth and produced consistent frequency band data before and after the EIS tests. The measured currents changed slightly at each depth. No unusual noise or transients were observed at any tests.

The results from what is believed to be the SNc in Figures 5 and 6 are high frequency currents. The difference between Figures 5 and 6 is consistent with a biologically formed conducting pathway, but they do not uniquely identify a ferritin/neuromelanin-associated conduction pathway. Extensive additional imaging analysis (e.g. electron microscope, conductive atomic force microscopy) would be needed to better understand the signaling pathways and the reasons for those differences and should be included as part of future tests. Referring to Table 1, 116 voltage step protocol tests were performed on the left and right hemispheres. As discussed further in the supplemental materials, the test results were repeatable at each depth and produced consistent frequency band data before and after the EIS tests. The measured currents changed slightly at each depth. No unusual noise or transients were observed at any tests.

The cortical results shown in Figure 7 were consistent with the expected results. Cortical tissue has normal levels of ferritin and neuromelanin that are typical of cellular iron homeostasis and waste material processing. It is noted that there is a small amount of Gaussian-type noise in the results, which serves as a point of reference for the presence of noise in the other tests. All tests were performed in the same room, with the same equipment and research team. The shielding, grounding, cortical-control measurements, saline measurements and agarose brain phantom measurements reduce the likelihood that the observed signals are dominated by environmental noise, although additional controls are needed to confirm the biological origin of the signals.

Referring to Table 1, 73 voltage step protocol tests were performed on the left and right hemispheres. As discussed further in the supplemental materials, the test results were repeatable at each depth and produced consistent data before and after the EIS tests. The measured currents changed slightly at each depth. No unusual noise or transients were observed at any tests.

The EIS data in Figure 8 was generally consistent with earlier EIS data from deep brain neural tissue tests. The EIS test uses signal processing such as filtering of high frequencies and root mean square processing, which prevents detection of high frequency transients. It was also conducted over repeated cycles, which could potentially modify the high-speed transient behavior of the ferritin and neuromelanin structures. Because the currents in ferritin and neuromelanin were expected to be at frequencies much higher than those compatible with EIS testing, the measured results were consistent with expectation.

Appendix 1 provides an in-depth analysis of the readings from July 11 from the right hemisphere at depths of 7.6 mm and 8.1 mm. It is noted that there are a number of different frequency bands, and that these generally remain consistent for all readings. However, there are some notable changes in magnitude, particularly after EIS readings when the tissue was exposed to higher levels of voltage and a range of frequencies. Those magnitude changes were measured after the completion of the EIS tests, and would not be the result of capacitive charge storage from the EIS test voltages. It is also noted that the frequency bands were present in the first reading taken before any voltage transitions, with the electrode at 0 volts. In that configuration, it is likely that currents were being measured directly from neural activity that is unrelated to the test voltages. While this data shows technical recurrence within a limited dataset, it is not sufficient to determine whether there is biological replication between different animals. Peak matching, within-site variability, clean in-out differences, spectral correlation, coefficient of variation, and/or a predefined rule would be needed to determine whether a response has been reproduced. The Appendix 1 data supports qualitative recurrence, but additional testing will be required to determine whether there is strong repeatability between different animals because the measurements were obtained from different locations. These data show technical recurrence within a limited dataset rather than biological replication.

Additional tests should include depth-matched deep controls in addition to replication of the cortical data. Examples of such controls could include intentional off-target tracks immediately adjacent to the SNc target, deep white-matter controls such as cerebral peduncle, carefully localized substantia nigra pars reticulata as a near-neighbor comparison, and deep non-catecholaminergic gray-matter targets at comparable depths. These additional controls would enable scientific conclusions to be reached regarding whether tissue ferritin levels are associated with high frequency signals or if other target specific differences between locations may be associated those signals.

The cortex data is not a sufficient negative control to draw definitive conclusions about the mechanism that is responsible for the high frequency currents in the deep SNc-region electrical test data. It does not control for depth (other than having readings taken at numerous different depths), electrode-tissue interface characteristics (other than using the same electrode-tissue interface for all tests), insertion effects (other than using the same insertion protocol for all tests), local impedance (other than using the same electrode configuration for all tests), vascular environment (even though vascular environment is not believed to be a relevant variable for the electrical data), or to provide sufficient stereotaxic certainty to precisely identify the specific area where the readings were taken (although the electrical data was generally identical).

## Conclusion

While the data obtained from this initial study provides evidence of high frequency signals in basal ganglia tissue that has not previously been detected, a fully developed stereotaxic workflow for the targets, three-dimensional histologic reconstruction and a significantly larger number of animal tests would be needed to further investigate the hypothesis of a ferritin-based signaling mechanism. The presence of high levels of ferritin and neuromelanin in catecholaminergic tissue has perplexed neuroscientists since their discovery. The test technique presented in this paper was successfully used to measure currents above 40 kHz in those tissues. While further testing is needed to investigate these currents, these results indicate that additional research is both needed and warranted. The test technique presented in this paper can be used for further in vivo investigation into these currents, but multiple electrodes or arrays could provide better data for analyzing the transfer function of these tissues.

## APPENDIX 1 Saline/agarose/in vivo comparison

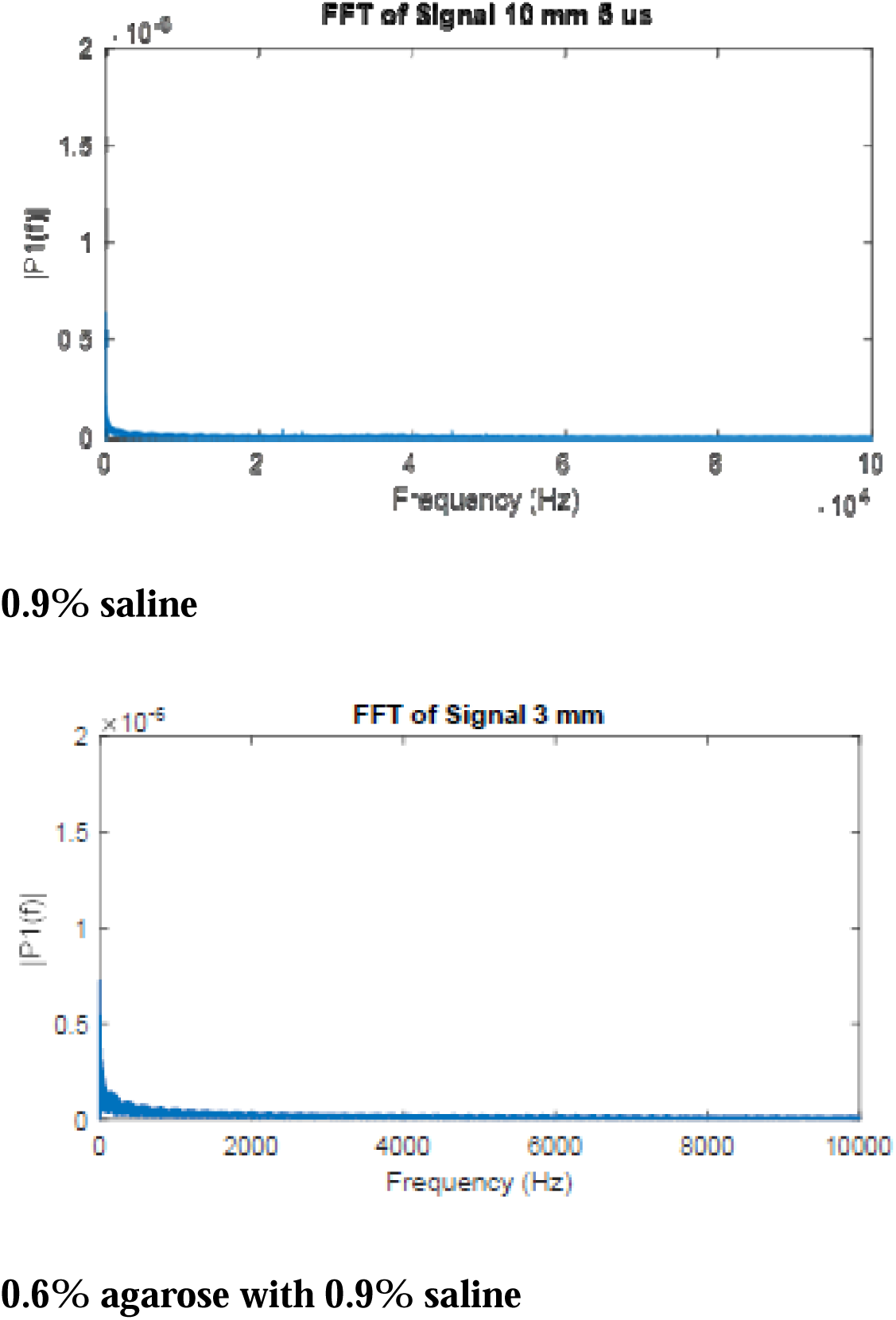
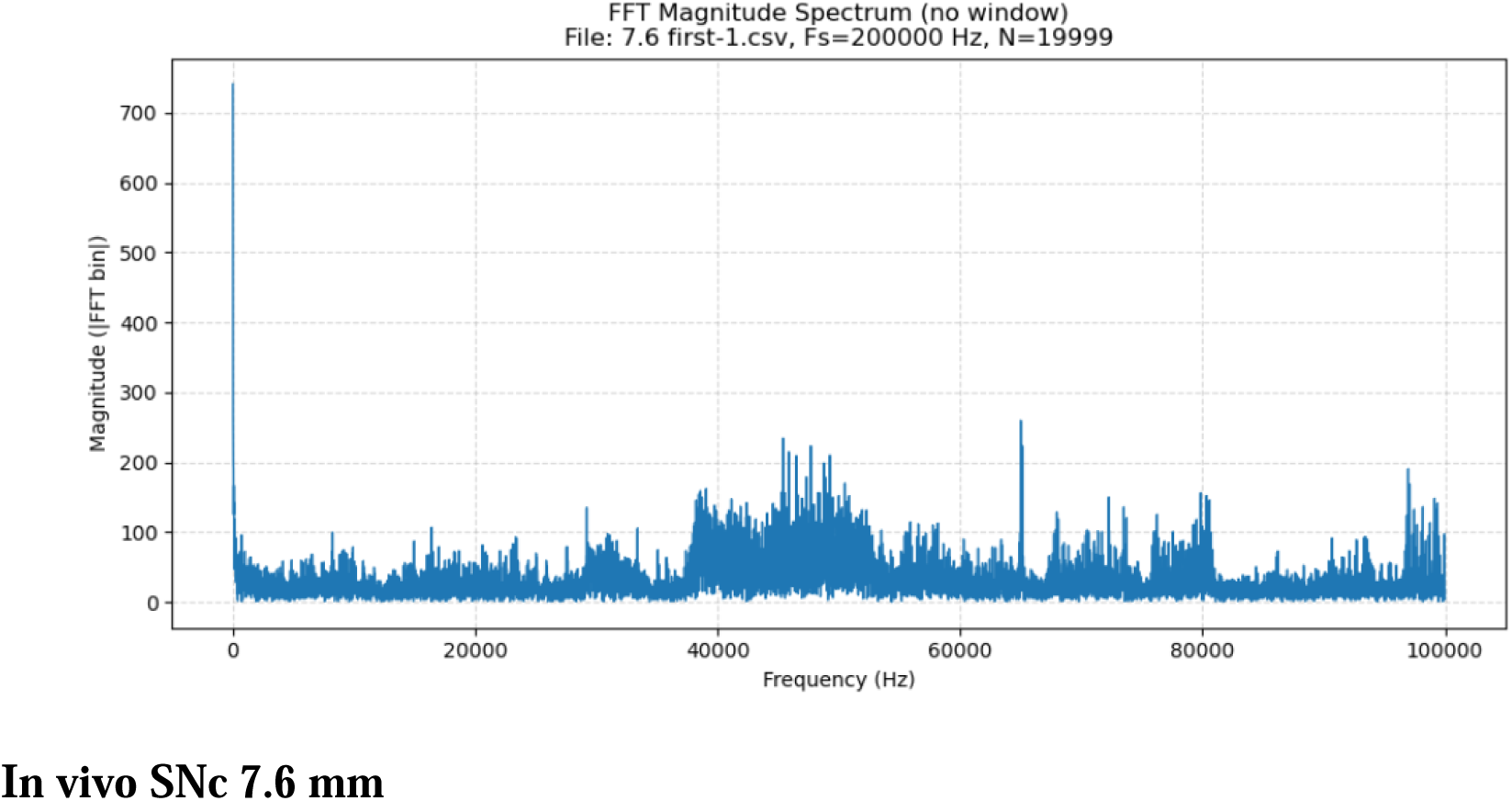

### FFT data for *in vivo* tests

#### 1. Right 7.6 mm, first pulse sequence, time 0-100 milliseconds

This measurement was taken with the electrode potential at 0.0 volts and prior to the transition to –1.0 volts. Based on the file sequence numbers and the time/date data in the file, this was the first measurement taken. The frequency transform shows bands between ∼37 kHz to 52 kHz, 56 kHz to 65 kHz with a spike at 65 kHz, 68 kHz to 74 kHz, 75 kHz to 81 kHz and 95 kHz to 100 kHz. There are no apparent 60 Hz or other low frequency noise components, indicating good shielding/grounding.

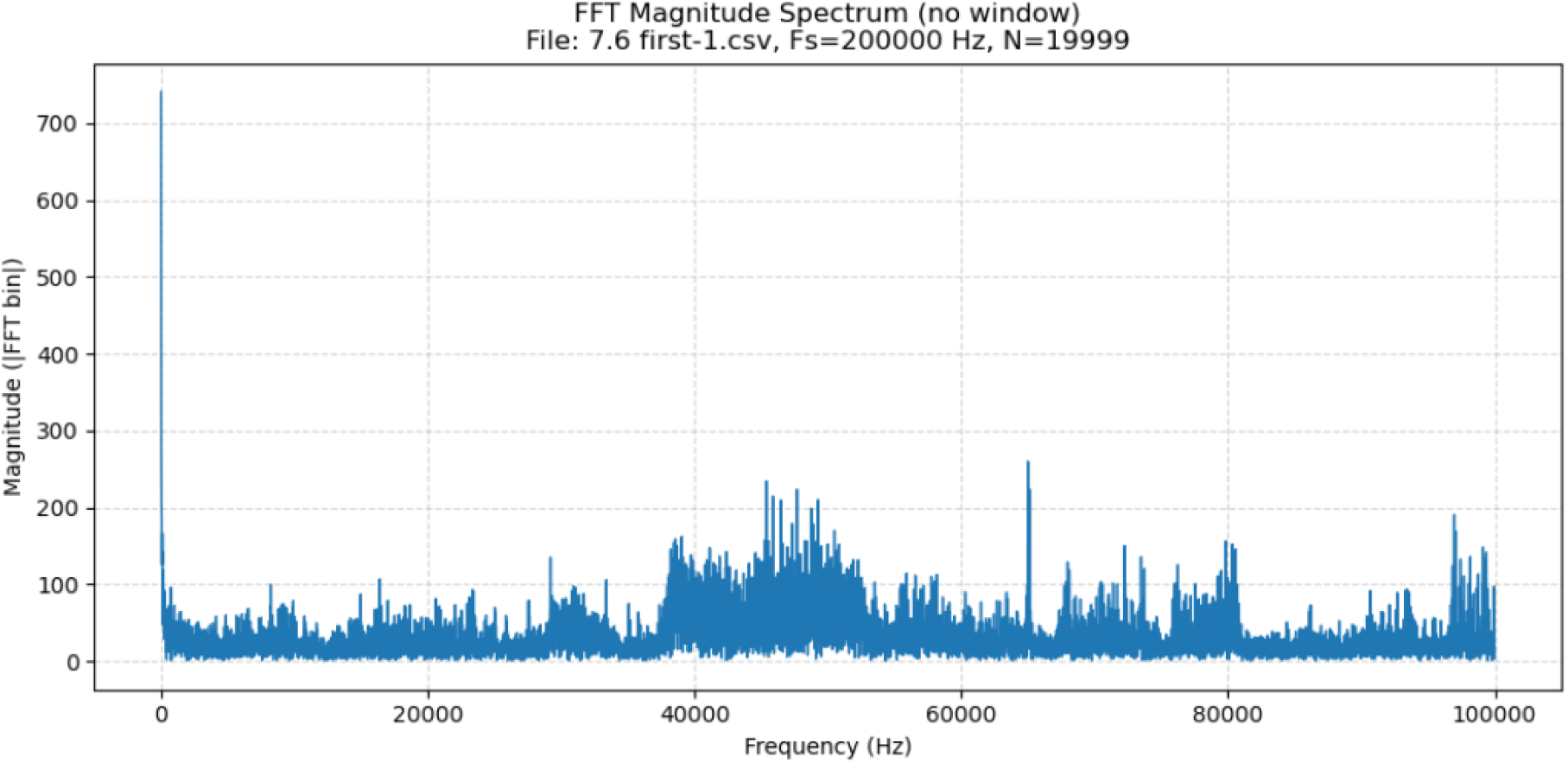
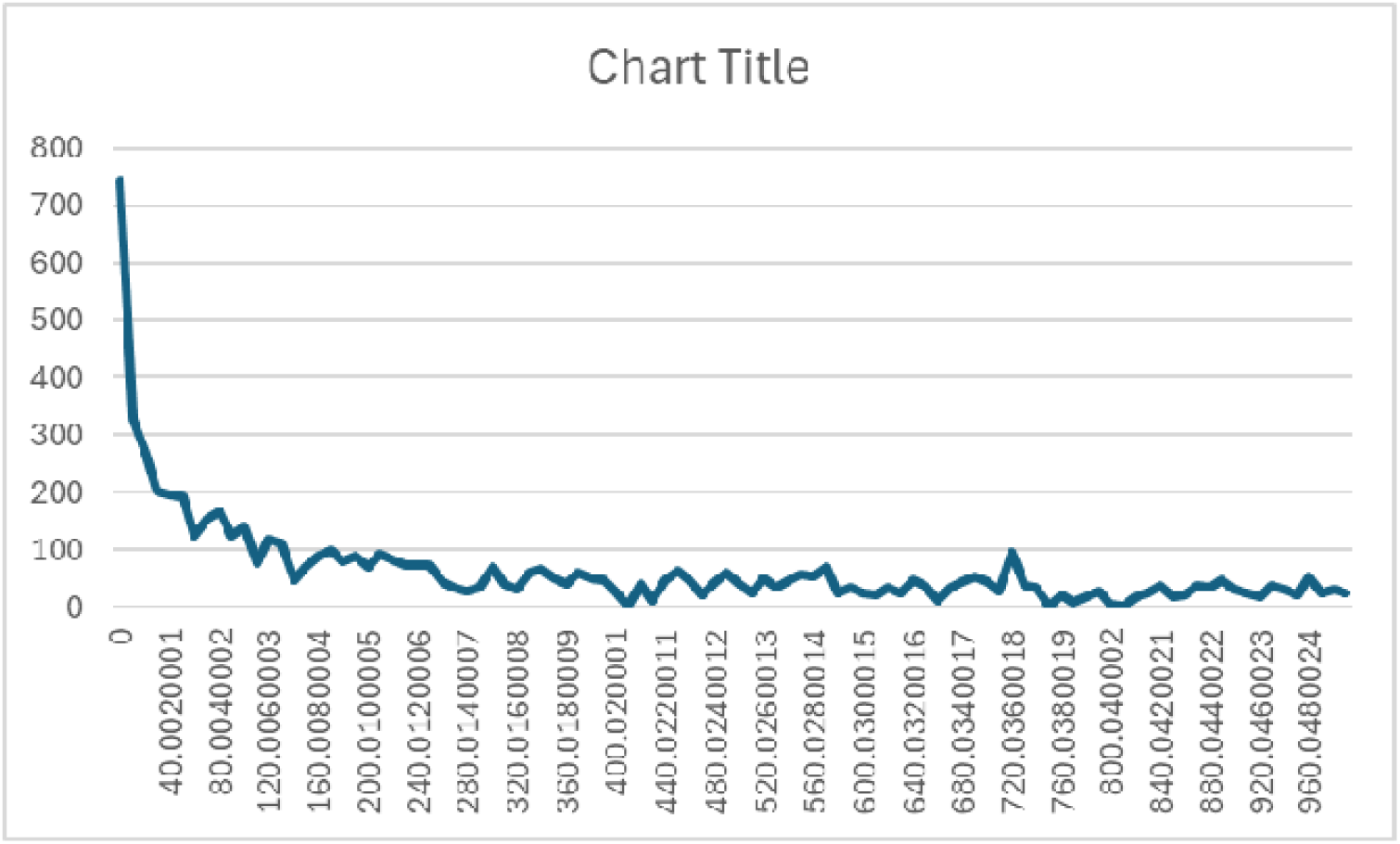

#### 2. Right 7.6 mm, first pulse sequence, time 100-200 milliseconds

This measurement was taken with the electrode potential at –1.0 volts after the transition from 0.0 volts. The frequency transform shows bands between ∼37 kHz to 52 kHz, 56 kHz to 65 kHz with a spike at 65 kHz, 68 kHz to 74 kHz, 75 kHz to 81 kHz and 95 kHz to 100 kHz, with changes to all bands from the 0-100 millisecond readings. There are no apparent 60 Hz or other low frequency noise components, indicating good shielding/grounding.

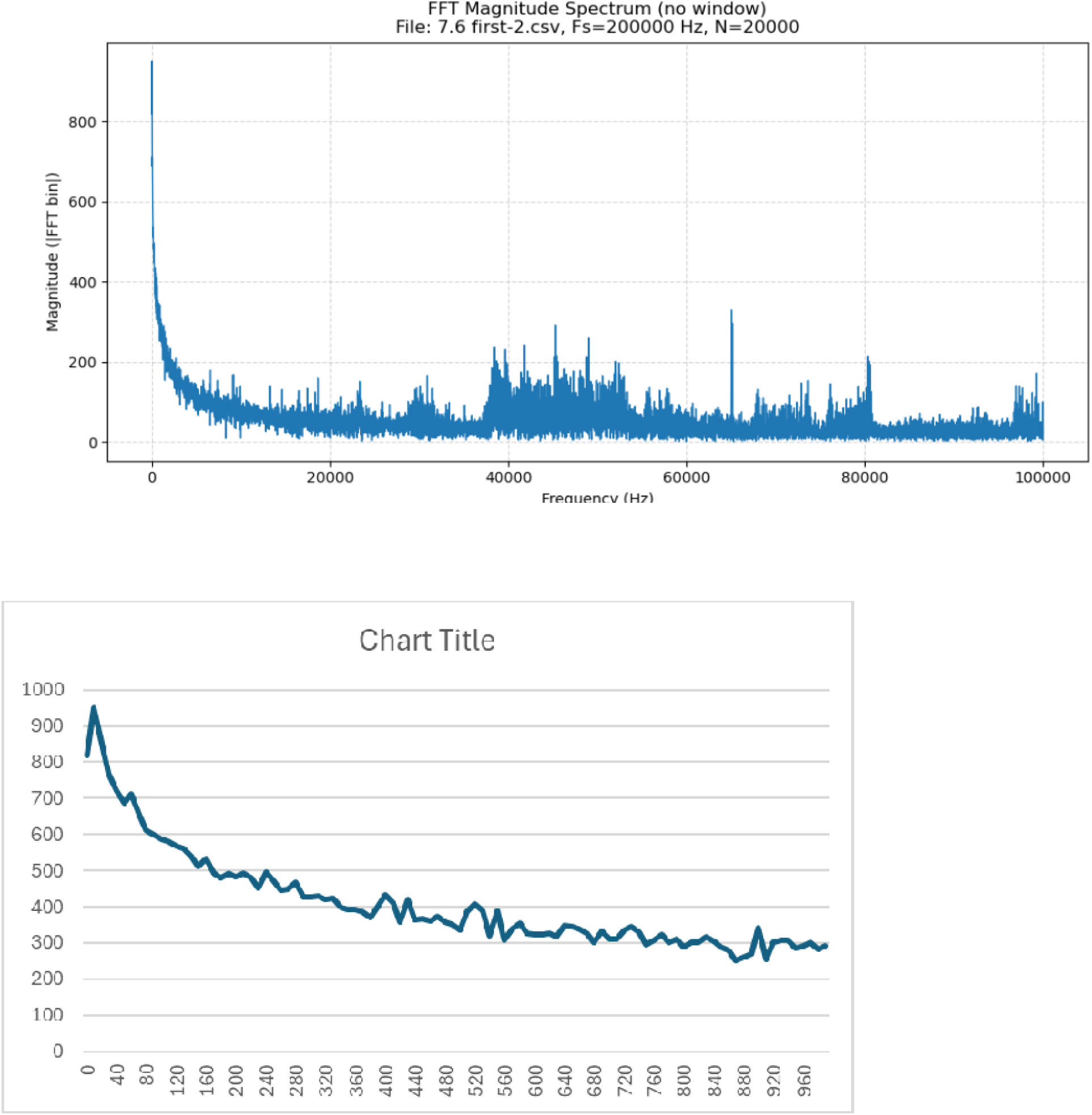

#### 3. Right 7.6 mm, first pulse sequence, time 200-300 milliseconds

This measurement was taken with the electrode potential at 0.0 volts after the transition from –1.0 volts. The frequency transform shows bands between ∼37 kHz to 52 kHz, 56 kHz to 65 kHz with a spike at 65 kHz, 68 kHz to 74 kHz, 75 kHz to 81 kHz and 95 kHz to 100 kHz, with changes to all bands from the 100-200 millisecond readings. While the magnitude of the bands looks lower, it is noted that the magnitude of the fundamental frequency component increased from 800 for the 100-200 millisecond transform to 1800, which offset the scale. As such, there does not appear to be any significant decrease, but there are apparent frequency profile changes from the 100-200 millisecond readings. There are no apparent 60 Hz or other low frequency noise components, indicating good shielding/grounding.

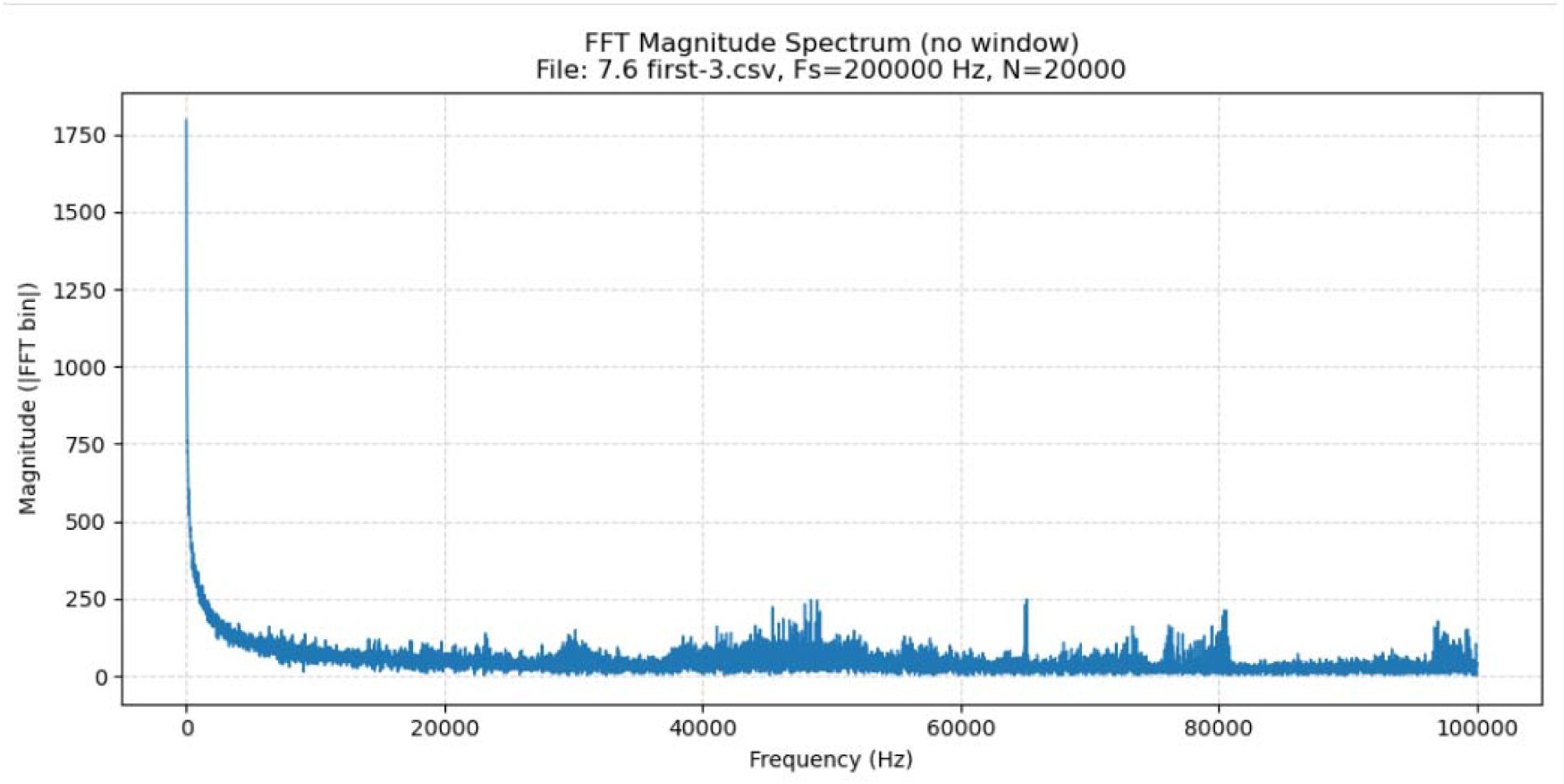
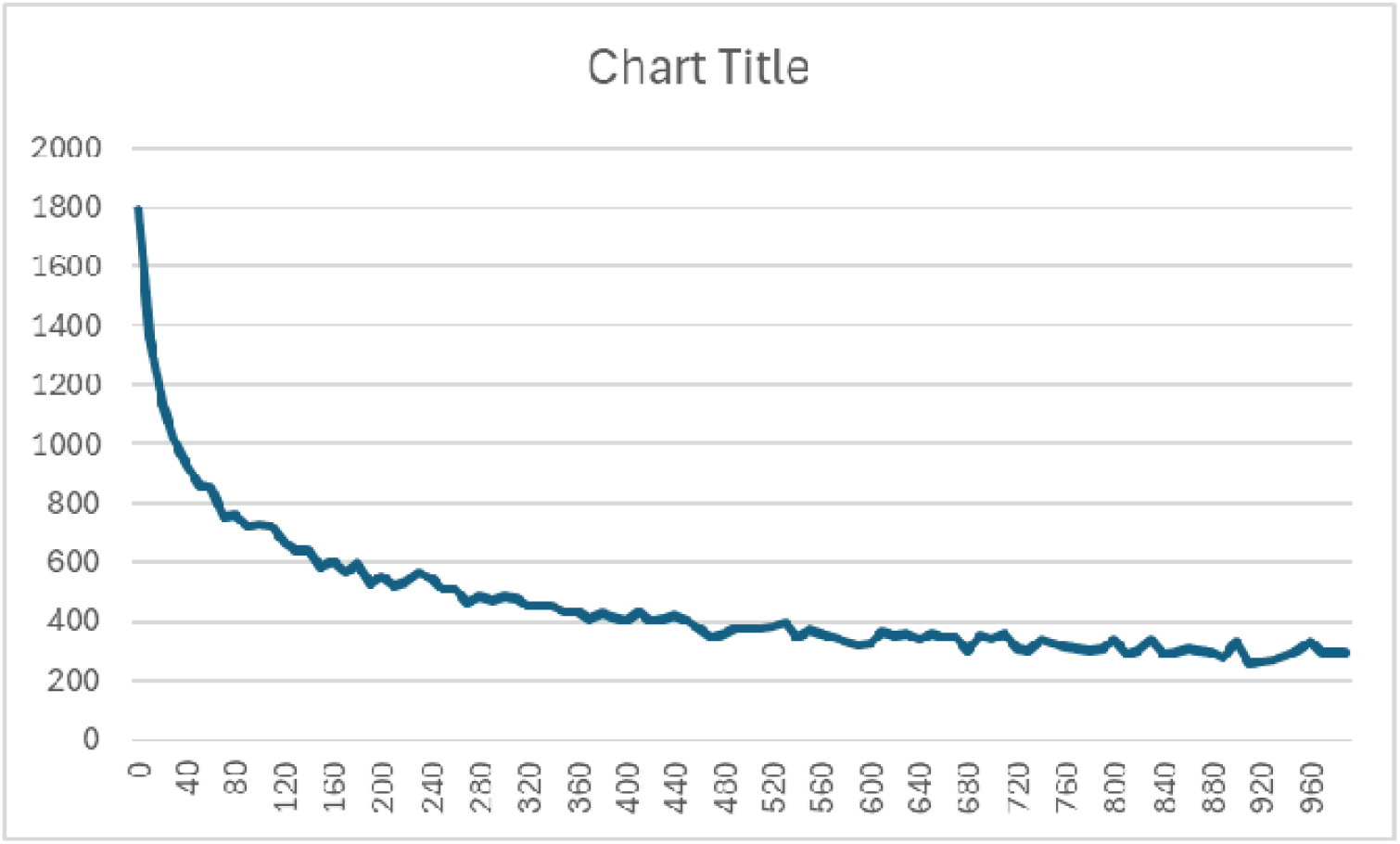

#### 4. Right 7.6 mm, first pulse sequence, time 300-400 milliseconds

This measurement was taken with the electrode potential at 1.0 volts after the transition from 0.0 volts. The frequency transform shows bands between ∼37 kHz to 52 kHz, 56 kHz to 65 kHz with a spike at 65 kHz, 68 kHz to 74 kHz, 75 kHz to 81 kHz and 95 kHz to 100 kHz, with changes to all bands from the 100-200 millisecond readings. While the magnitude of the bands looks lower, it is noted that the magnitude of the fundamental frequency component increased from 800 for the 100-200 millisecond transform to 2100, which offset the scale. As such, there does not appear to be any significant decrease, but there are apparent frequency profile changes from the 200-300 millisecond readings. There are no apparent 60 Hz or other low frequency noise components, indicating good shielding/grounding.

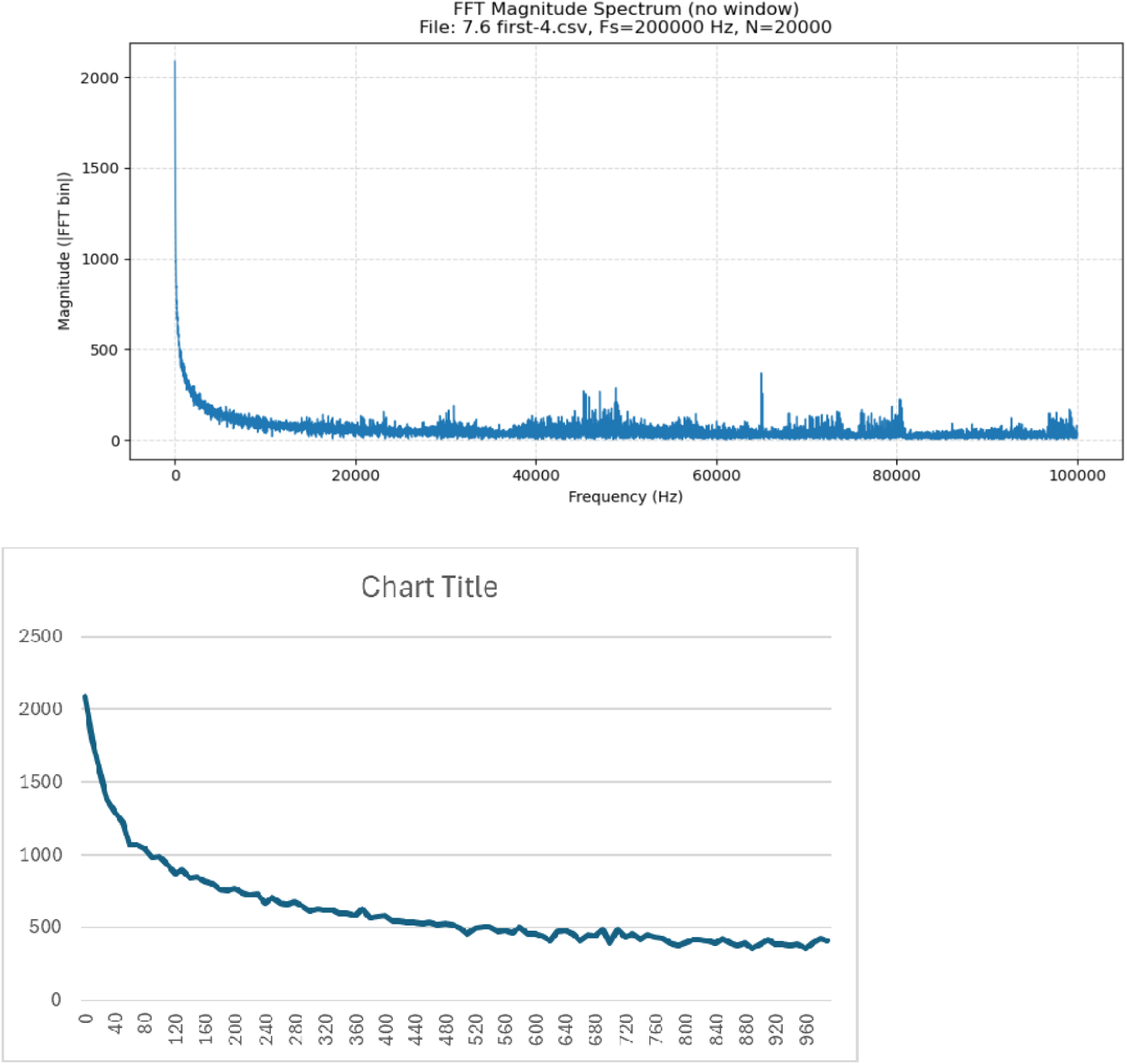

#### 5. Right 7.6 mm, first pulse sequence, time 400-500 milliseconds

This measurement was taken with the electrode potential at 0.0 volts after the transition from 1.0 volts. The frequency transform shows bands between ∼37 kHz to 52 kHz, 56 kHz to 65 kHz with a spike at 65 kHz, 68 kHz to 74 kHz, 75 kHz to 81 kHz and 95 kHz to 100 kHz, with changes to all bands from the 100-200 millisecond readings. While the magnitude of the bands looks lower, it is noted that the magnitude of the fundamental frequency component increased from 800 for the 100-200 millisecond transform to 1500, which offset the scale. As such, there does not appear to be any significant decrease, but there are apparent frequency profile changes from the 300-400 millisecond readings. There are no apparent 60 Hz or other low frequency noise components, indicating good shielding/grounding.

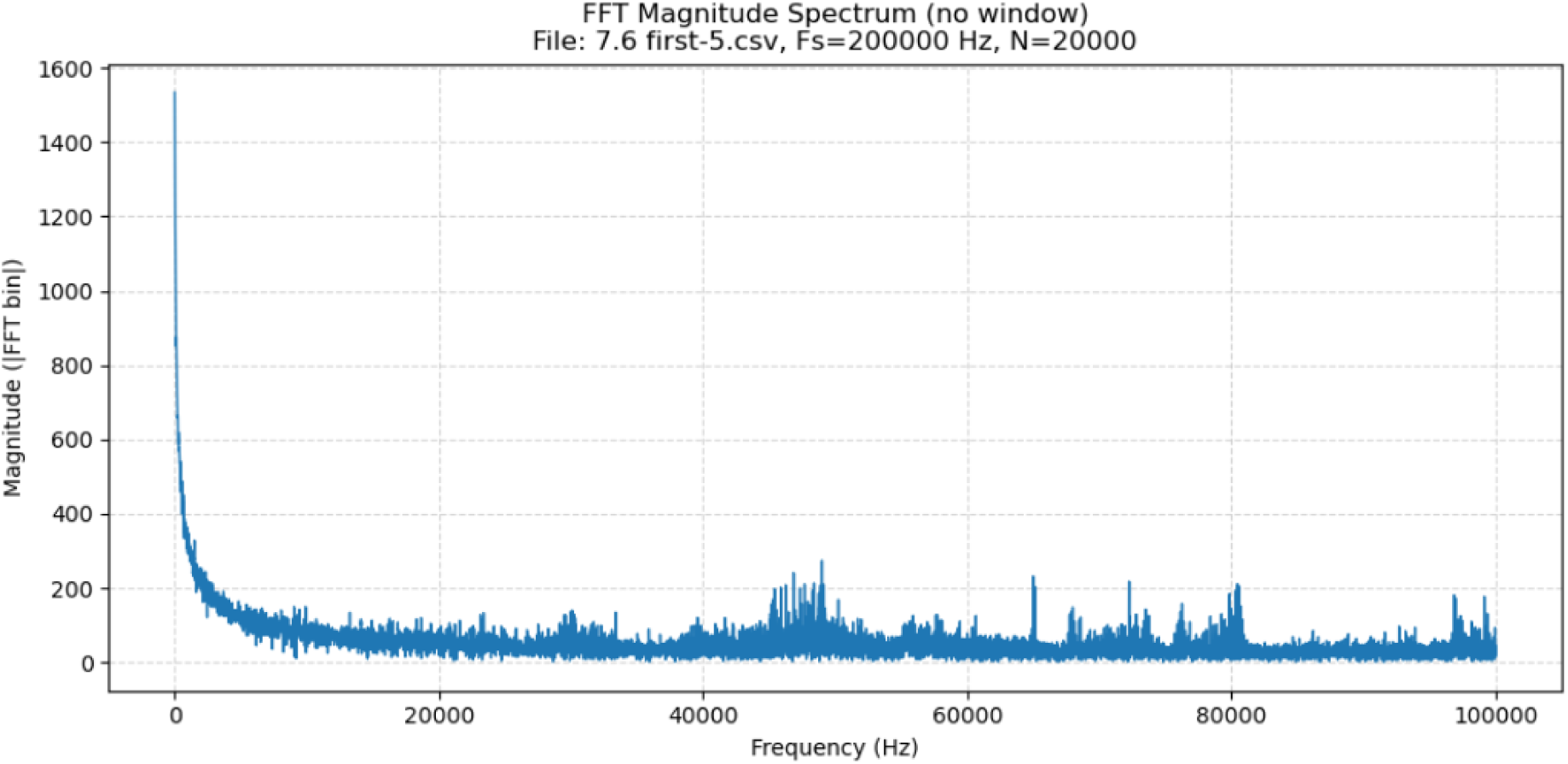
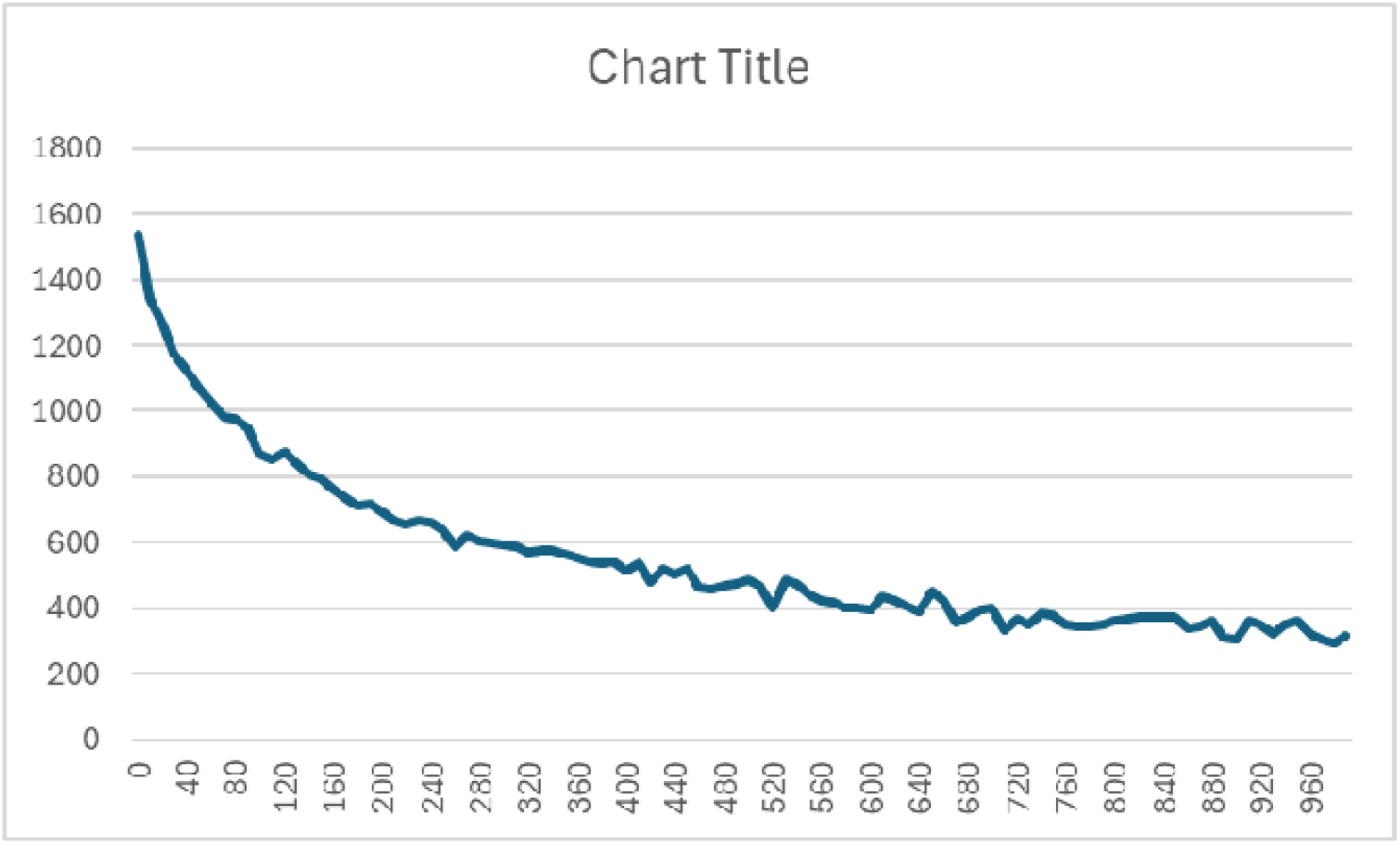

#### 6. Right 7.6 mm, second pulse sequence, 100-200 milliseconds

This measurement was taken with the electrode potential at –1.0 volts after the transition from 0.0 volts. The frequency transform shows bands between ∼37 kHz to 52 kHz, 56 kHz to 65 kHz with a spike at 65 kHz, 68 kHz to 74 kHz, 75 kHz to 81 kHz and 95 kHz to 100 kHz, with changes to all bands from the 100-200 millisecond readings for the first pulse sequence. There does not appear to be any significant decrease, but there are apparent frequency profile changes from the 100-200 millisecond readings from the first pulse sequence. There are no apparent 60 Hz or other low frequency noise components, indicating good shielding/grounding.

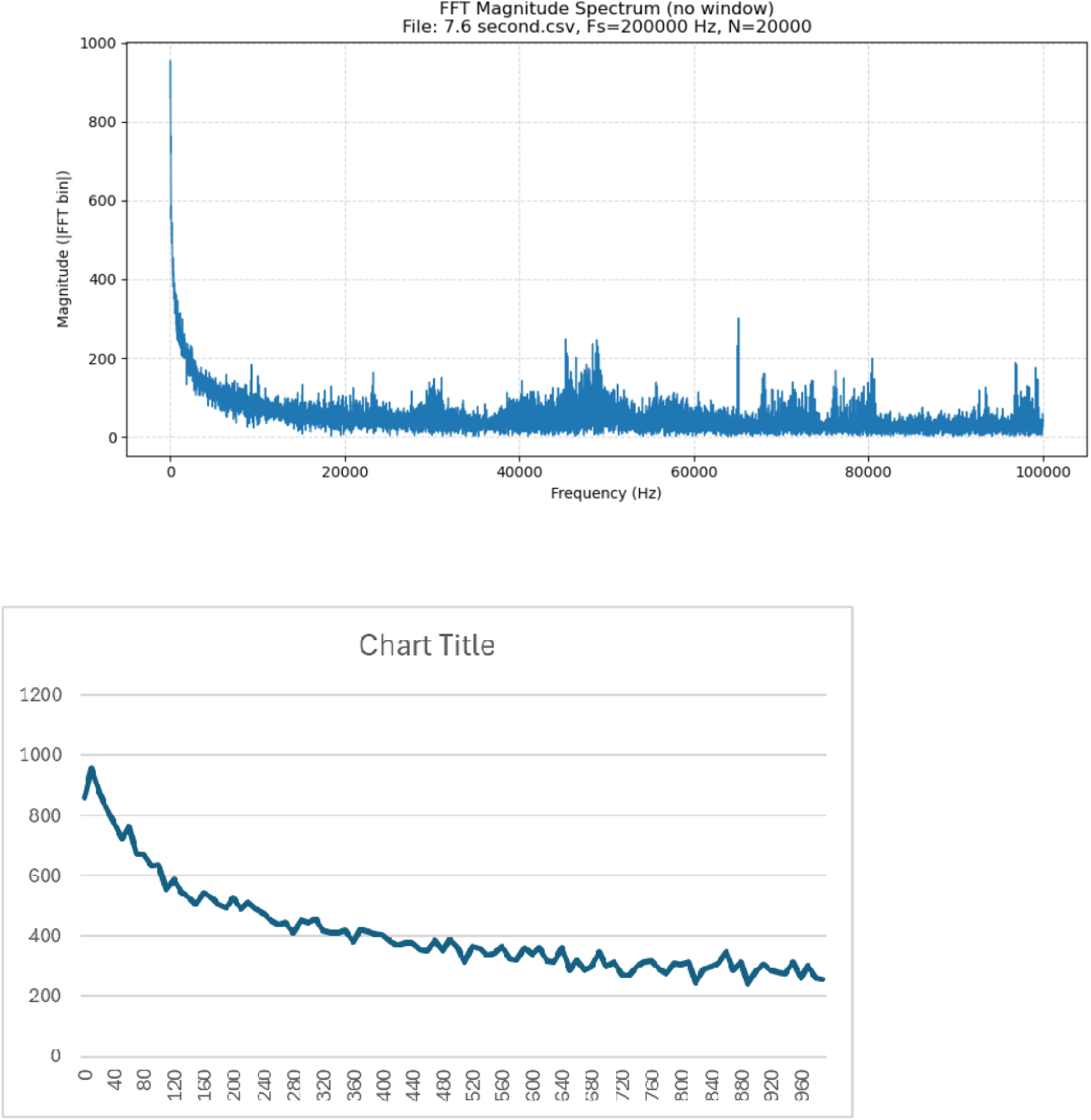

#### 7. Right 7.6 mm, third pulse sequence, 100-200 milliseconds

This measurement was taken after the EIS readings (which apply a higher voltage at different frequencies) with the electrode potential at –1.0 volts after the transition from 0.0 volts. The frequency transform shows bands between ∼37 kHz to 52 kHz, 56 kHz to 65 kHz with a spike at 65 kHz, 68 kHz to 74 kHz, 75 kHz to 81 kHz and 95 kHz to 100 kHz, with changes to all bands from the 100-200 millisecond readings for the first pulse sequence. There are significant increases 37 kHz to 40 kHz, as well as other apparent frequency profile changes from the 100-200 millisecond readings from the second pulse sequence. There are no apparent 60 Hz or other low frequency noise components, indicating good shielding/grounding.

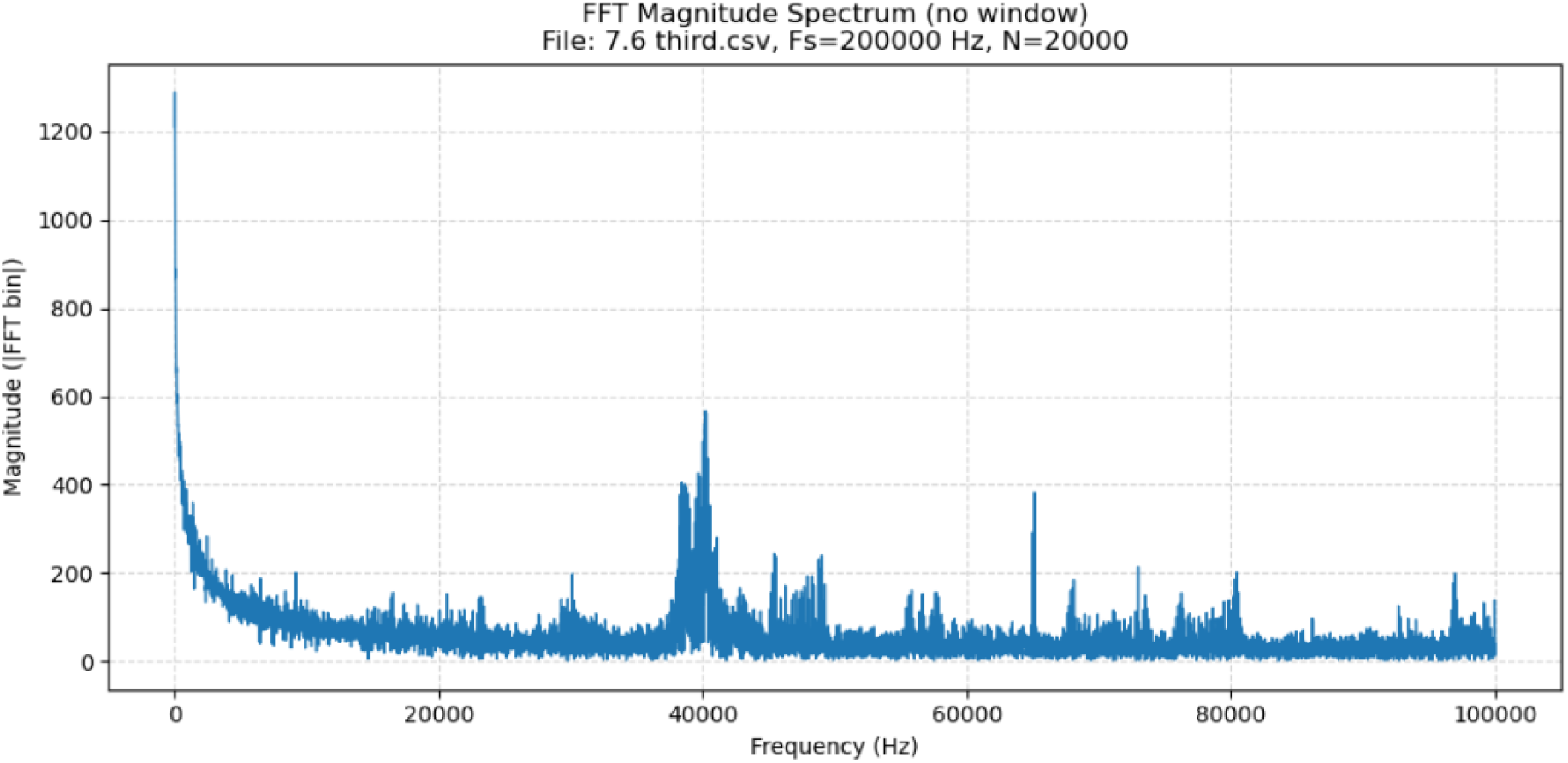
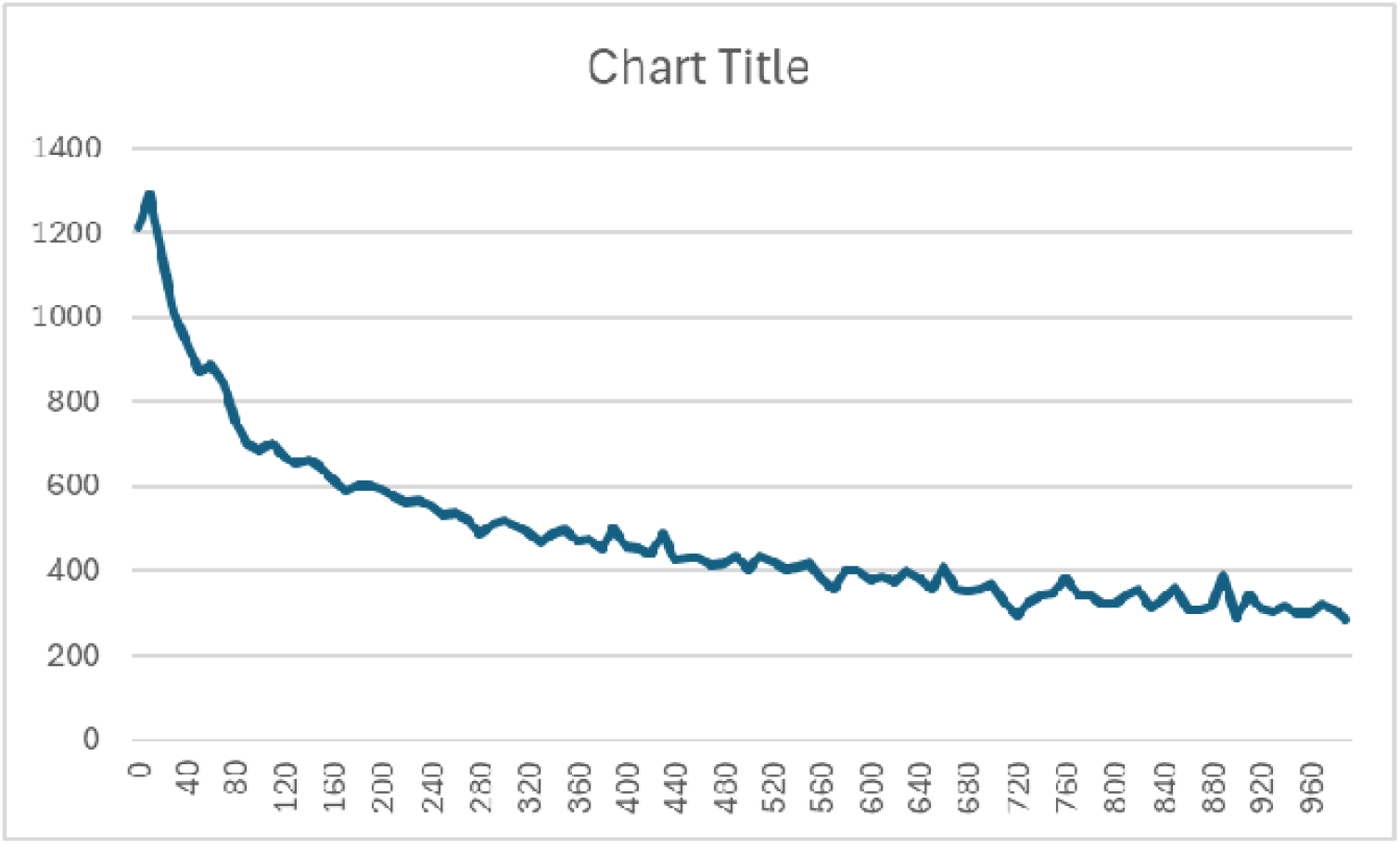

#### 8. Right 7.6 mm, fourth pulse sequence, 100-200 milliseconds

This measurement was taken with the electrode potential at –1.0 volts after the transition from 0.0 volts. The frequency transform shows bands between ∼37 kHz to 52 kHz, 56 kHz to 65 kHz with a spike at 65 kHz, 68 kHz to 74 kHz, 75 kHz to 81 kHz and 95 kHz to 100 kHz, with changes to all bands from the 100-200 millisecond readings for the first pulse sequence. There are significant increases 37 kHz to 40 kHz, as well as other apparent frequency profile changes from the 100-200 millisecond readings from the third pulse sequence. There are no apparent 60 Hz or other low frequency noise components, indicating good shielding/grounding.

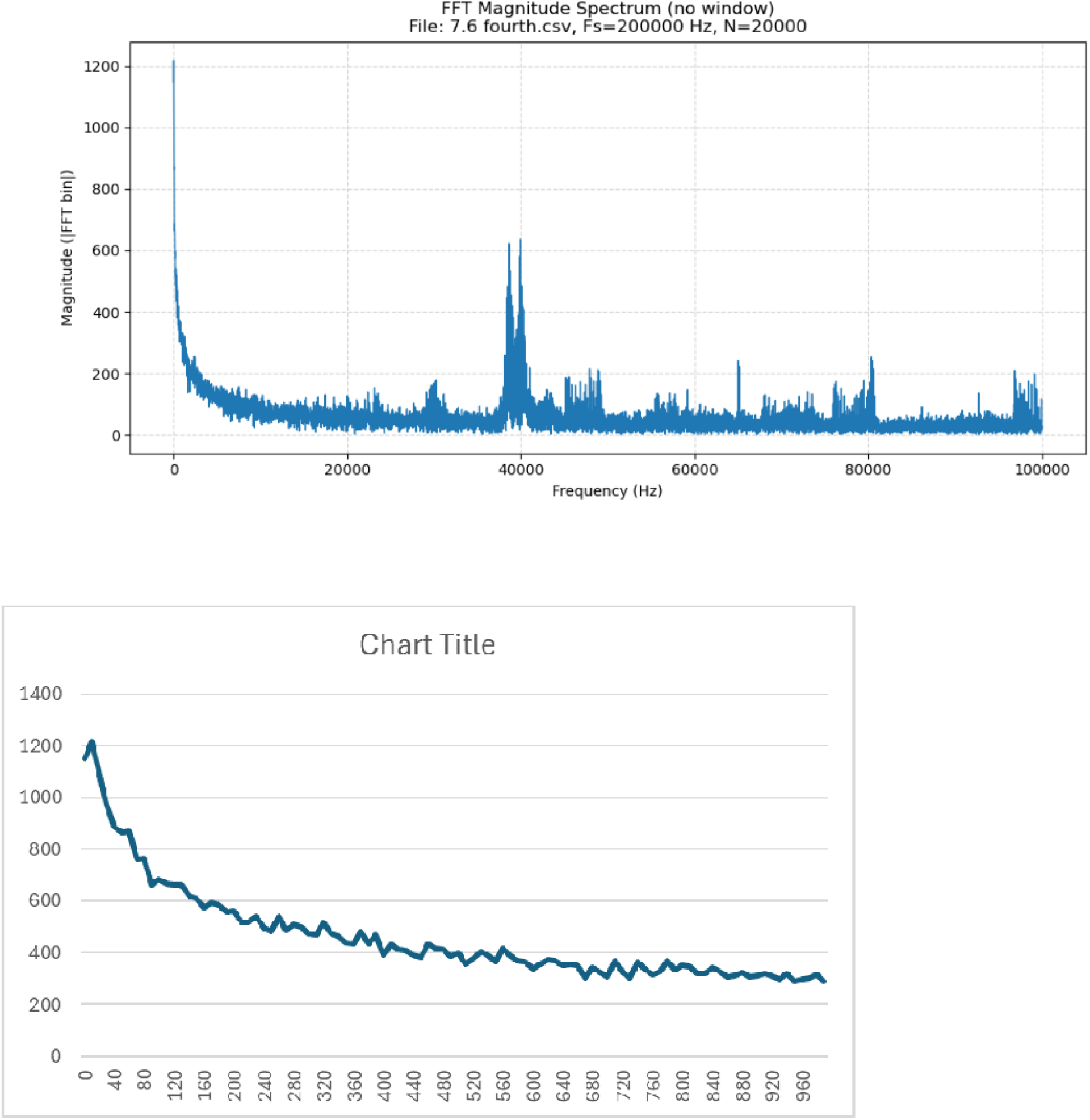

#### 9. Right 8.1 mm, first pulse sequence, 100-200 milliseconds

This measurement was taken with the electrode potential at –1.0 volts after the transition from 0.0 volts. The frequency transform shows bands between ∼37 kHz to 50 kHz, 56 kHz to 58 kHz, a spike at 65 kHz, 68 kHz to 74 kHz, 75 kHz to 81 kHz and 95 kHz to 100 kHz, with changes to all bands from the 100-200 millisecond readings for the first pulse sequence. There are apparent frequency profile changes from the 100-200 millisecond readings from the 7.6 mm depth readings. There are no apparent 60 Hz or other low frequency noise components, indicating good shielding/grounding.

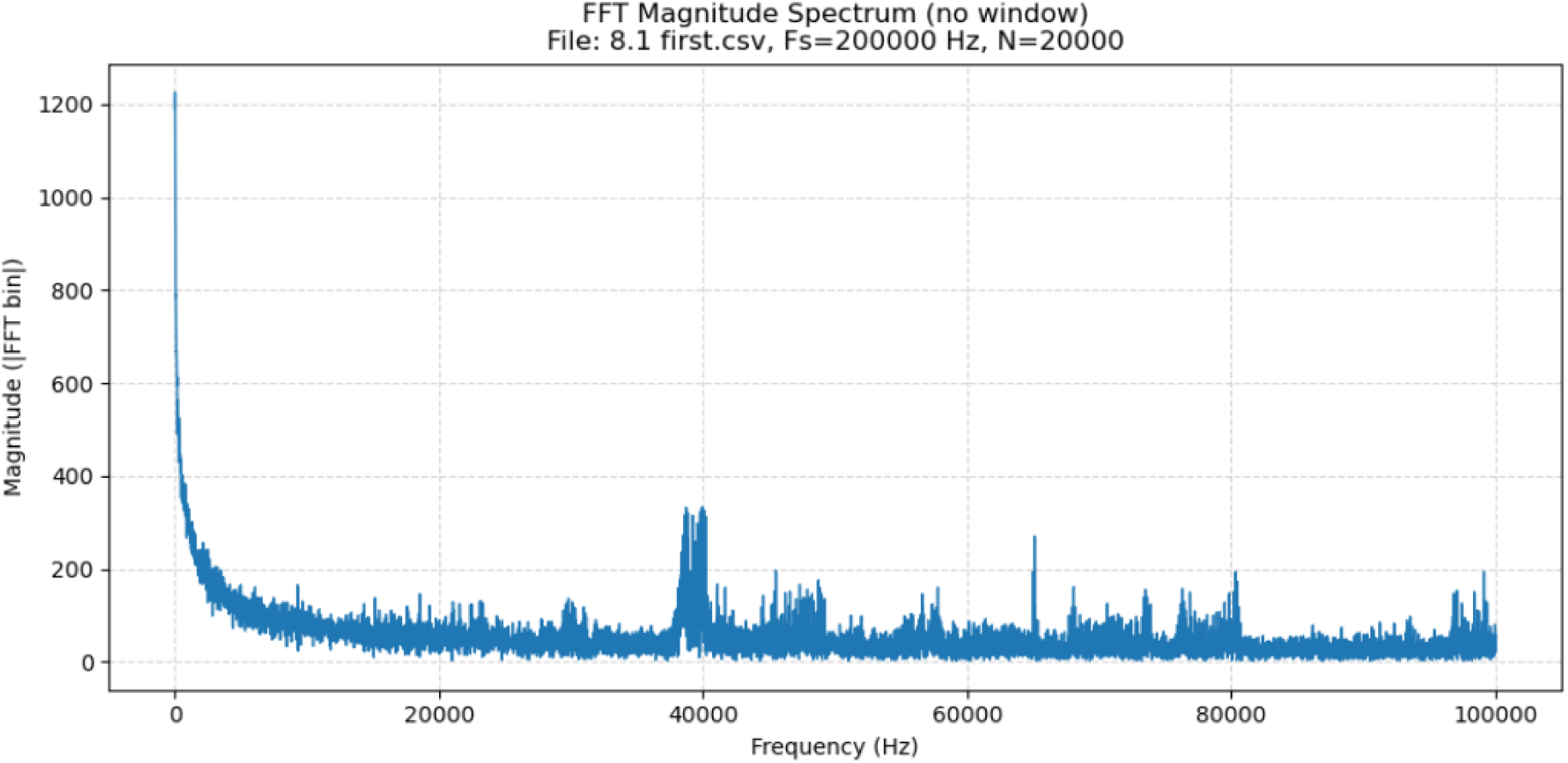
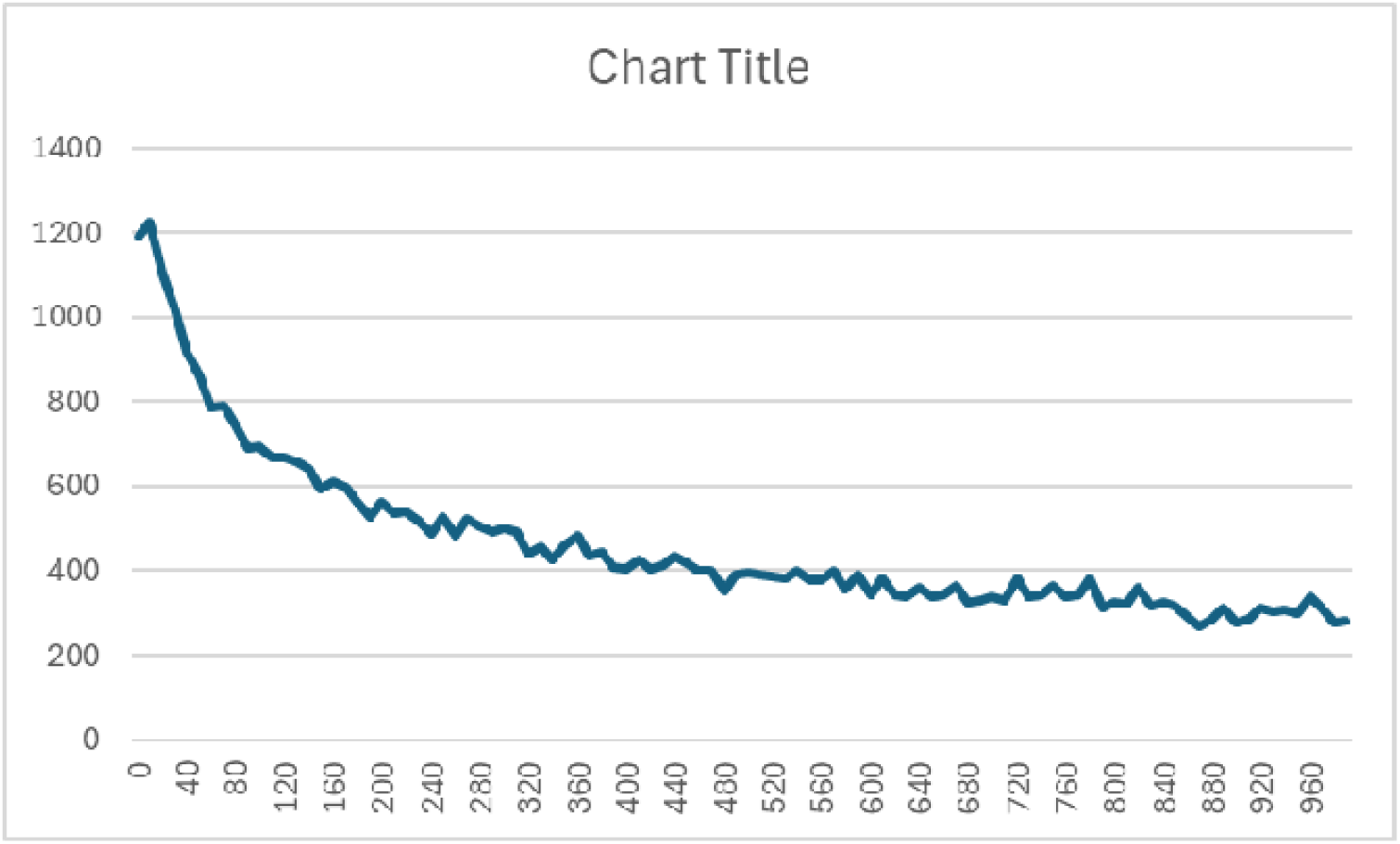

#### 10. Right 8.1 mm, second pulse sequence, 100-200 milliseconds

This measurement was taken with the electrode potential at –1.0 volts after the transition from 0.0 volts. The frequency transform shows bands between ∼37 kHz to 50 kHz, 56 kHz to 58 kHz, a spike at 65 kHz, 68 kHz to 74 kHz, 75 kHz to 81 kHz and 95 kHz to 100 kHz, with changes to all bands from the 100-200 millisecond readings for the first pulse sequence. There are apparent frequency profile changes from the 100-200 millisecond readings from the first pulse sequence readings. There are no apparent 60 Hz or other low frequency noise components, indicating good shielding/grounding.

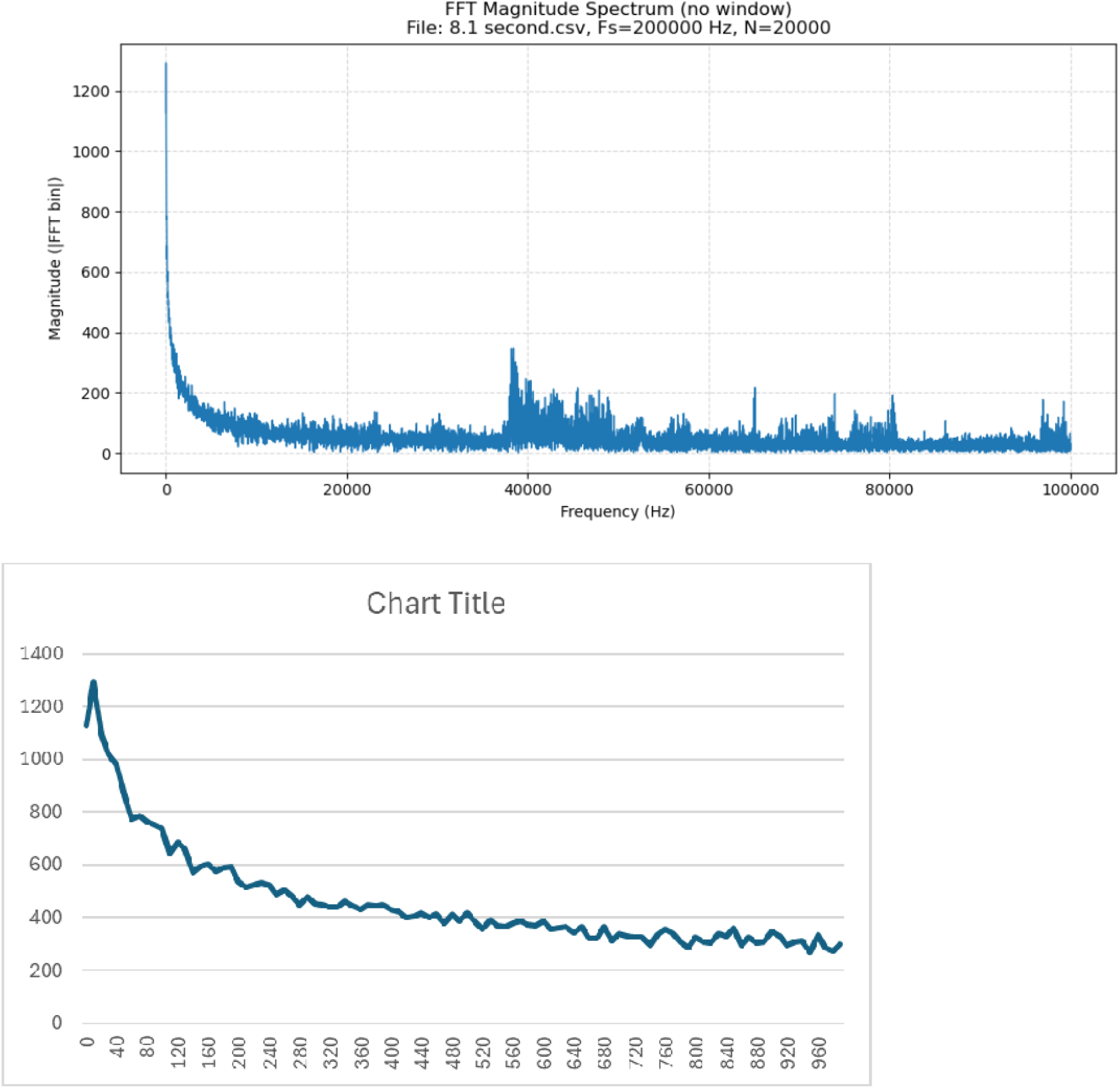

#### 11. Right 8.1 mm, third pulse sequence, 100-200 milliseconds

This measurement was taken after the EIS measurements, with the electrode potential at – 1.0 volts after the transition from 0.0 volts. The frequency transform shows bands between ∼37 kHz to 50 kHz, 56 kHz to 58 kHz, a spike at 65 kHz, 68 kHz to 74 kHz, 75 kHz to 81 kHz and 95 kHz to 100 kHz, with changes to all bands from the 100-200 millisecond readings for the first pulse sequence. There are apparent frequency profile changes from the 100-200 millisecond readings from the second pulse sequence readings. There are no apparent 60 Hz or other low frequency noise components, indicating good shielding/grounding.

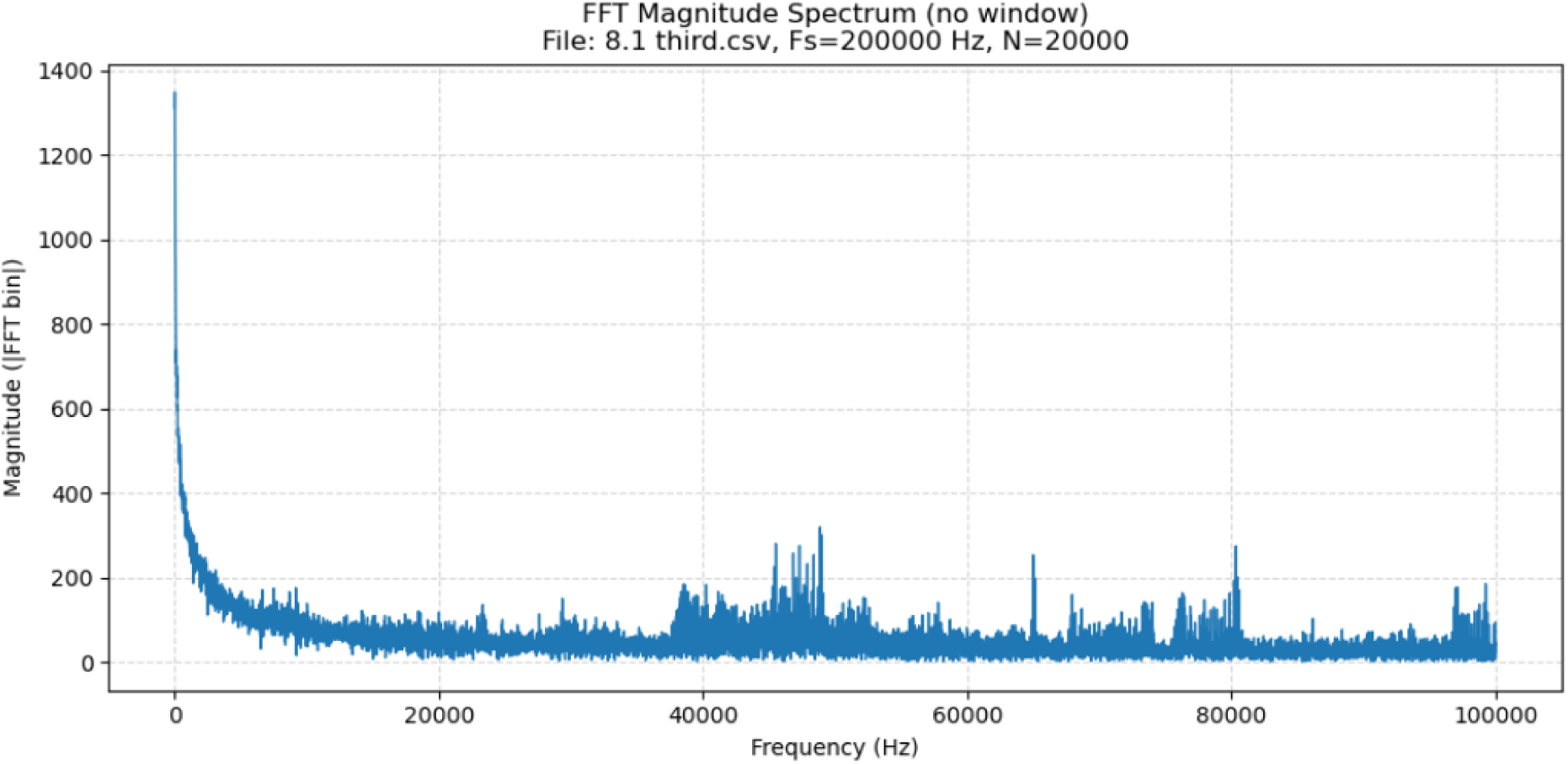
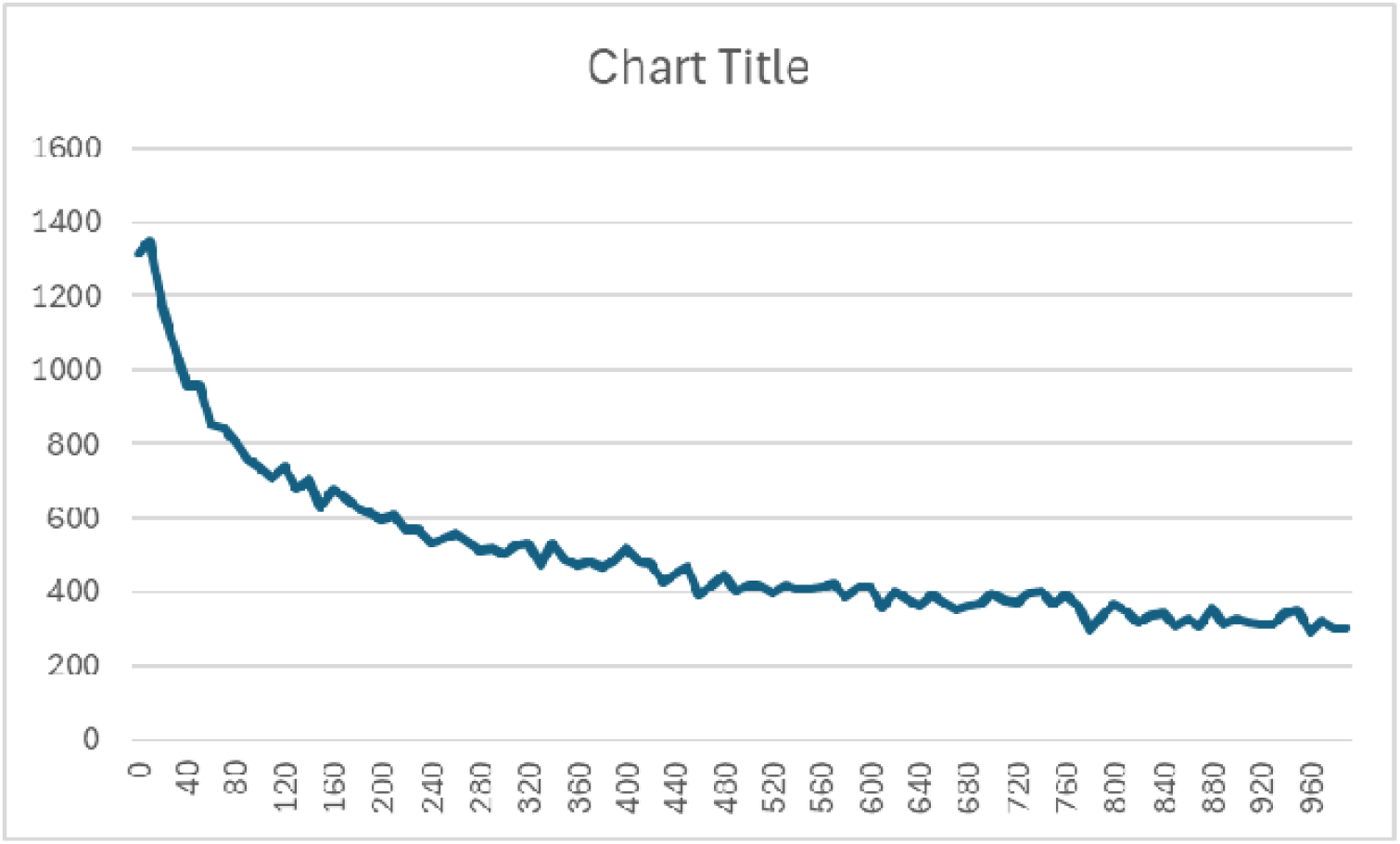

#### 12. Right 8.1 mm, fourth pulse sequence, 100-200 milliseconds

This measurement was taken with the electrode potential at –1.0 volts after the transition from 0.0 volts. The frequency transform shows bands between ∼37 kHz to 50 kHz, 56 kHz to 58 kHz, a spike at 65 kHz, 68 kHz to 74 kHz, 75 kHz to 81 kHz and 95 kHz to 100 kHz, with changes to all bands from the 100-200 millisecond readings for the first pulse sequence. There are apparent frequency profile changes from the 100-200 millisecond readings from the third pulse sequence readings. There are no apparent 60 Hz or other low frequency noise components, indicating good shielding/grounding.

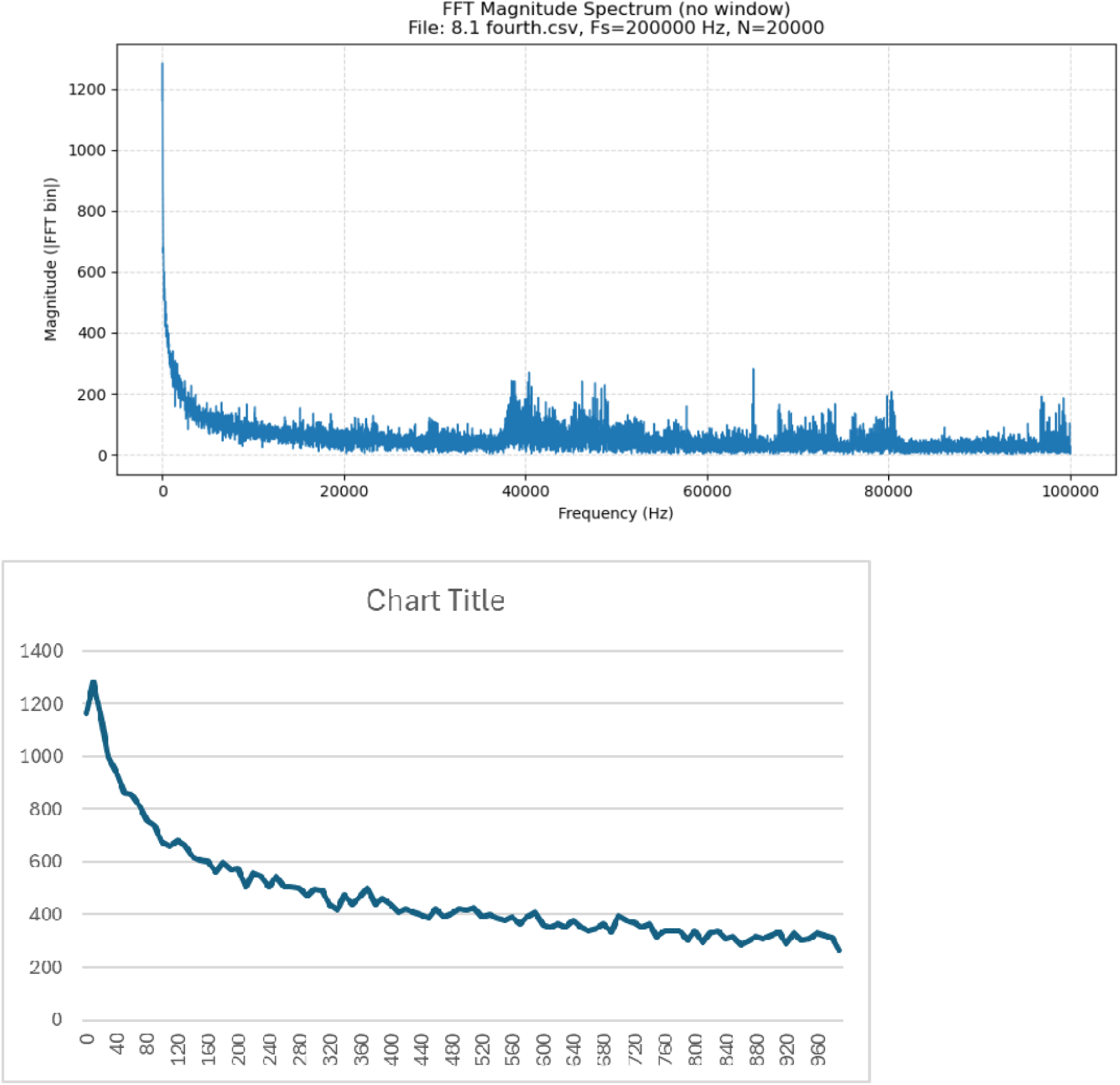

#### 13. Left 7.6 mm, fist pulse sequence 100 to 200 milliseconds

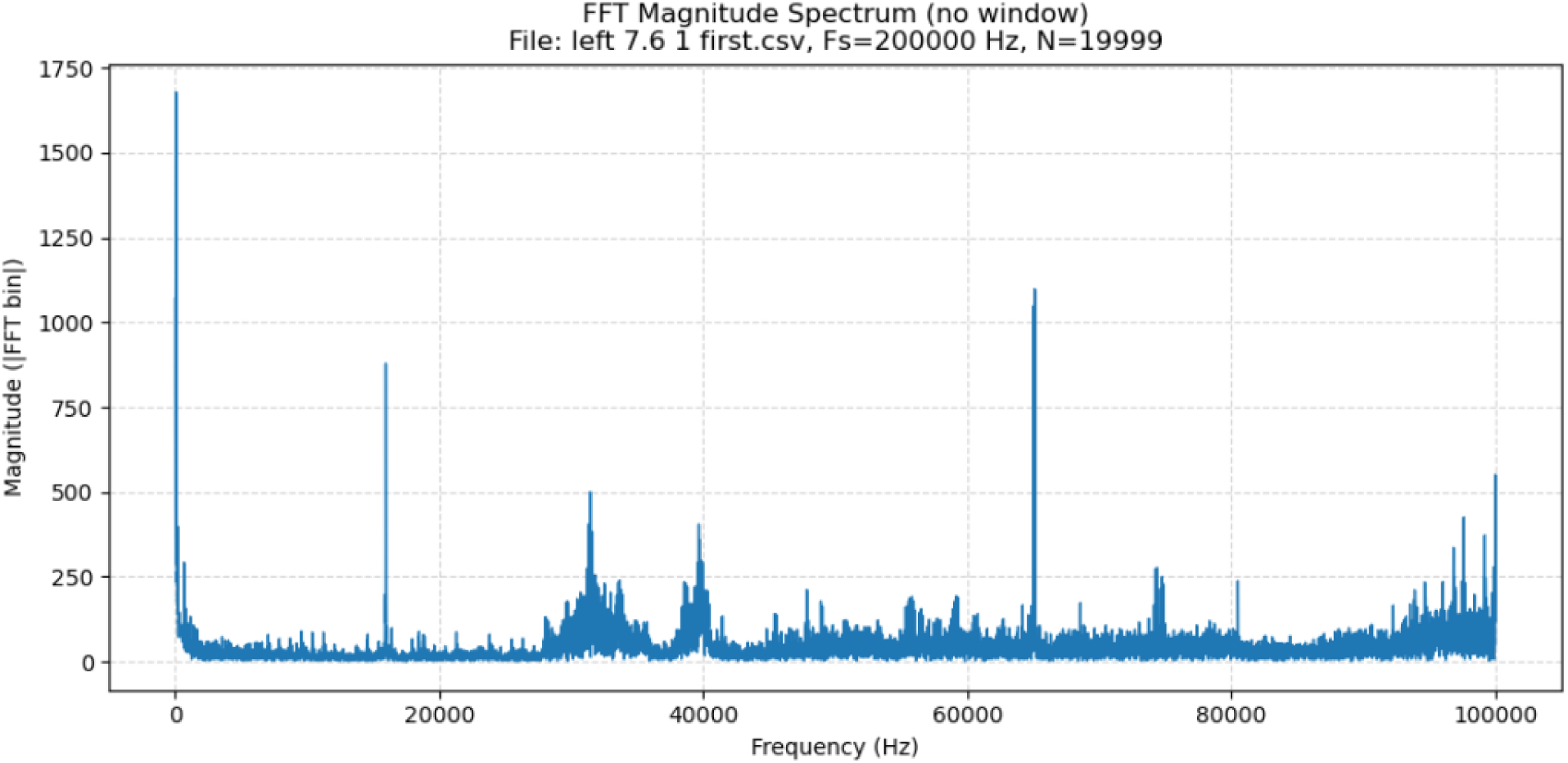

### Python FFT analysis code

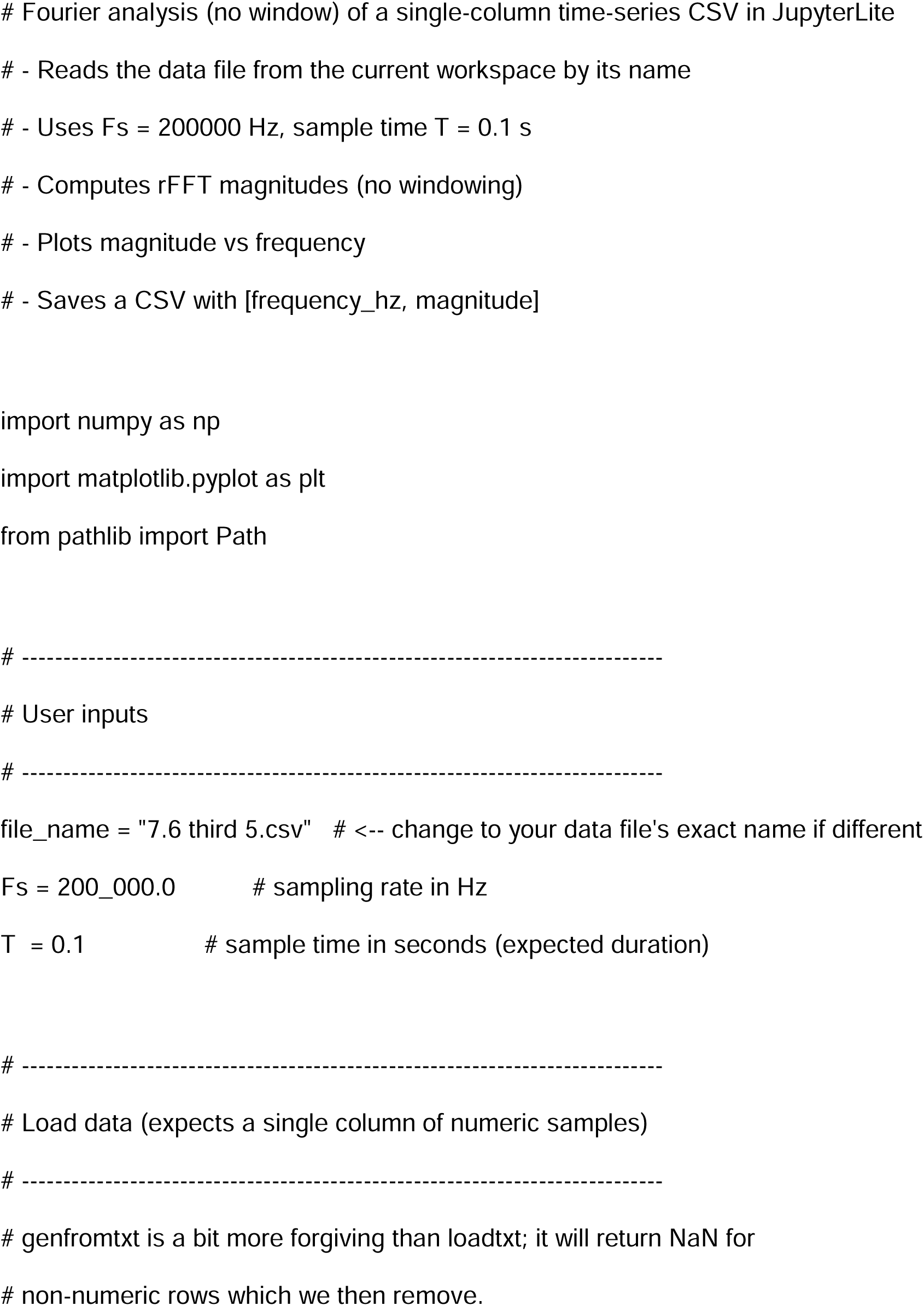
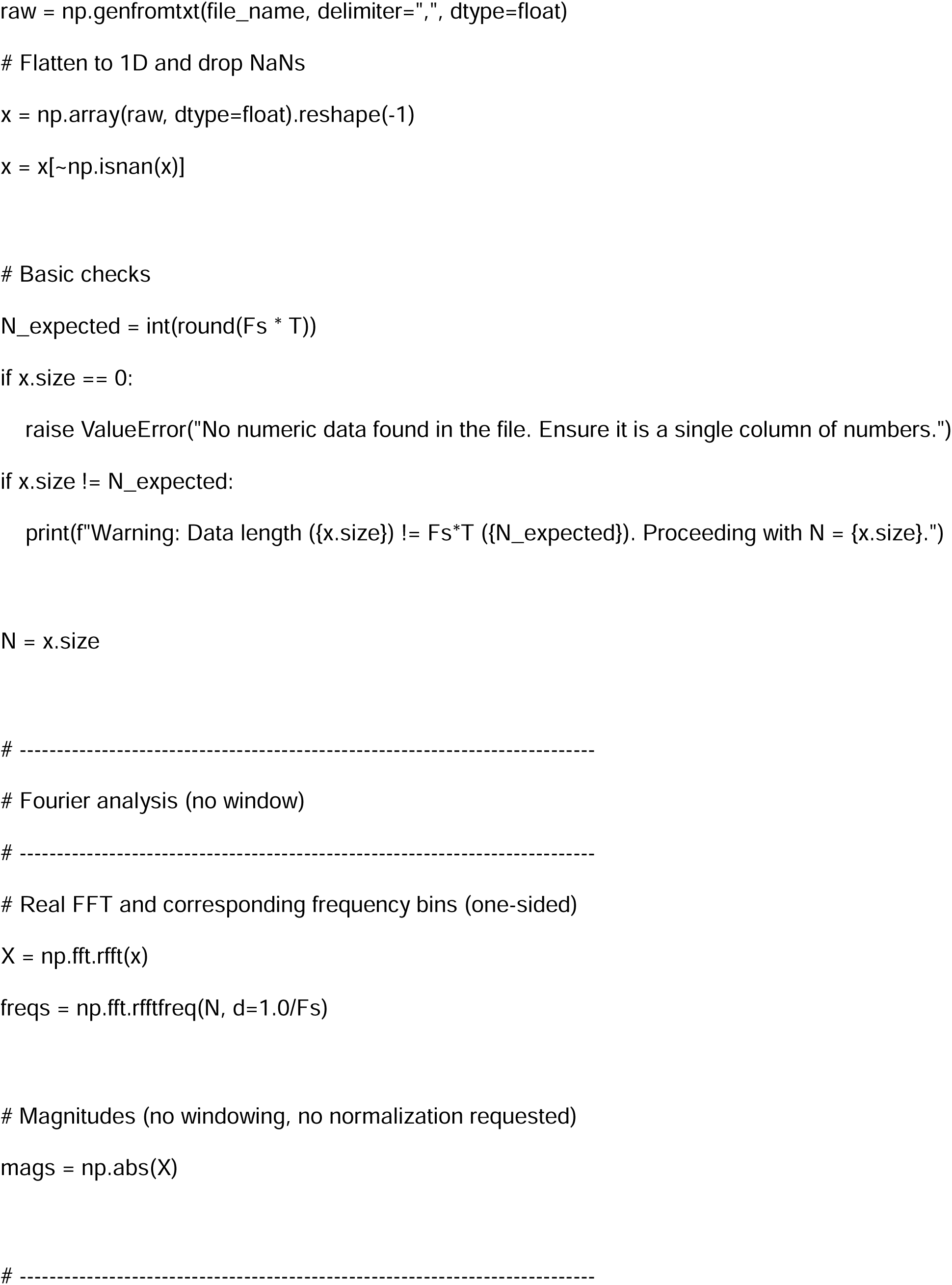
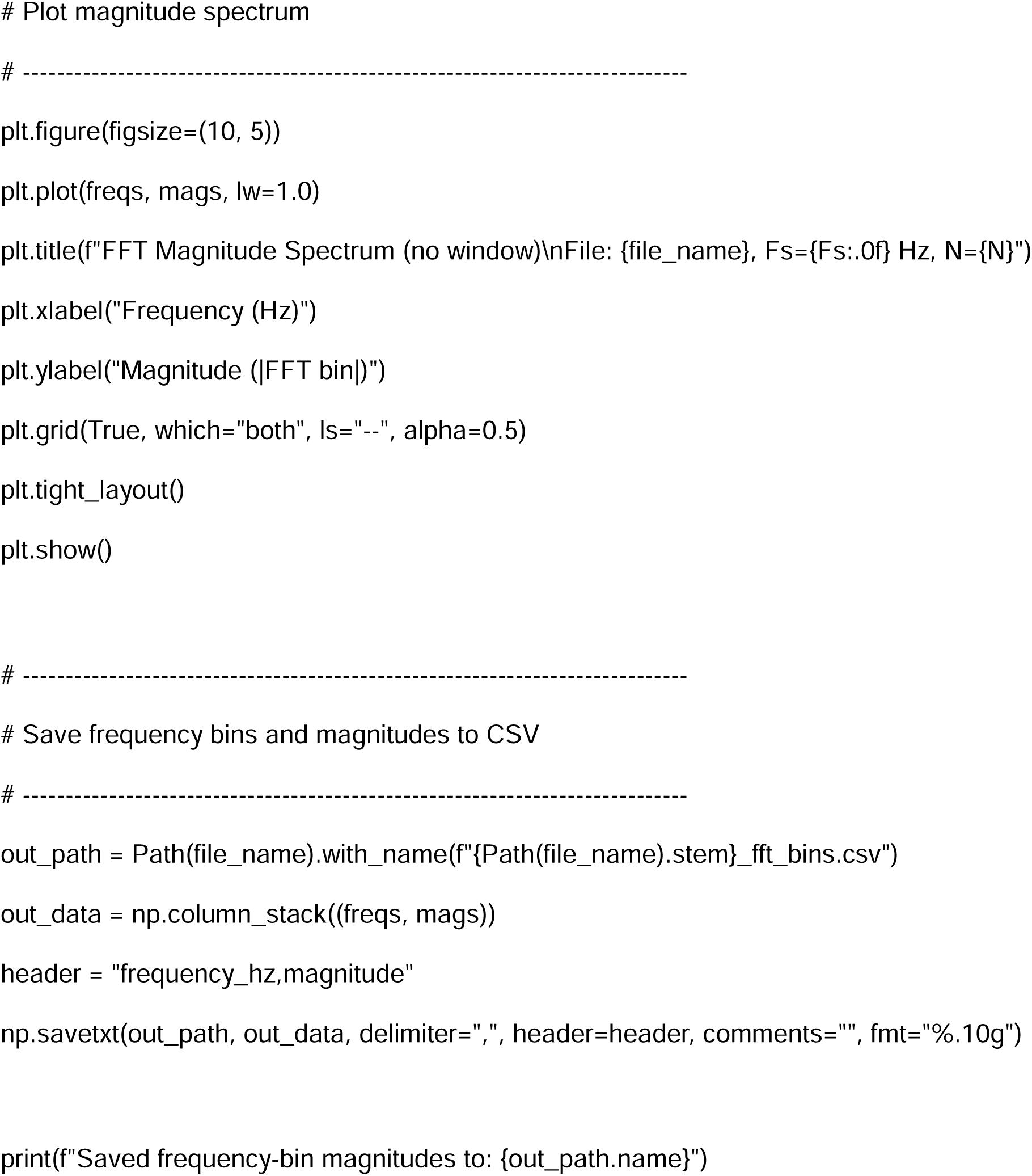

## APPENDIX 2 Similarity/difference analysis

### 1. Similarity and difference metrics

To determine a numerical metric for the similarity of two FFT plots, the cosine similarity value was used. This value treats the magnitude of each frequency bin as a vector and measures the cosine of the angle between them. A value of 1.0 indicates identical spectral shapes (regardless of overall amplitude scaling), while 0.0 would indicate completely orthogonal spectra and –1.0 would indicate a mirror spectra.

To determine a numerical metric for the difference between two FFT plots, the root mean square error (RMSE) was used. This value represents the average magnitude of the bin-by-bin differences between the two spectra, and is expressed in the same units as the raw magnitude values. A low value indicates a smaller difference and a larger value indicates a greater difference.

The results discussed below show that the sequence of scans from the same location at successive time steps have closer correlation (greater similarity, less difference) than scans at other adjacent locations, and that scans for the same type of tissue (dopaminergic neurons, non-dopaminergic neurons) have a closer correlation than scans for different tissue. Additional comparisons are also discussed herein.

1. SNc, same location, comparison before and after EIS tests (first scan versus third scan at same time sequence). High similarity, lower difference.

2. SNc, same location, comparison to different time sequence measurements. High similarity, mostly lower difference.

| Metric | Value |
| --- | --- |
| Cosine Similarity | 0.776353 |
| RMSE | 66.344604 |

3. Cortex left hemisphere versus right hemisphere at 5 mm depth (very high similarity, very low difference), and right hemisphere at 5 mm depth versus SNc right hemisphere at 7.6 mm (low similarity, high difference).

2. SNc, same location, comparison to different time sequence measurements. High similarity, mostly lower difference.
| Time sequence comparison | Cosine Similarity | RMSE |
| --- | --- | --- |
| (0-100) versus (100-200) | 0.832 | 60.53 |
| (0-100) versus (200-300) | 0.800856 | 70.949456 |
| (0-100) versus (300-400) | 0.703225 | 99.950157 |
| (0-100) versus (400-500) | 0.780381 | 75.121526 |
| (300-400) versus (400-500) | 0.873896 | 68.385782 |

| Comparison | CS | RMSE |
| --- | --- | --- |
| Cortex, right 5 mm vs left 5 mm (second time step) | 0.992979 | 14.201485 |
| Cortex, right 5 mm second time step vs. SNc right 7.6 mm first time step | 0.3507 | 111.88 |
| Cortex, right 5 mm second time step vs. SNc right 7.6 mm first second<br>step | 0.5594 | 98.08 |

### 2. Right hemisphere SNc, depth 7.6 mm, first pulse sequence, first period (0 to 100 ms) <u>versus</u> right hemisphere SNc, depth 7.6 mm, third pulse sequence, first period (0 to 100 ms)

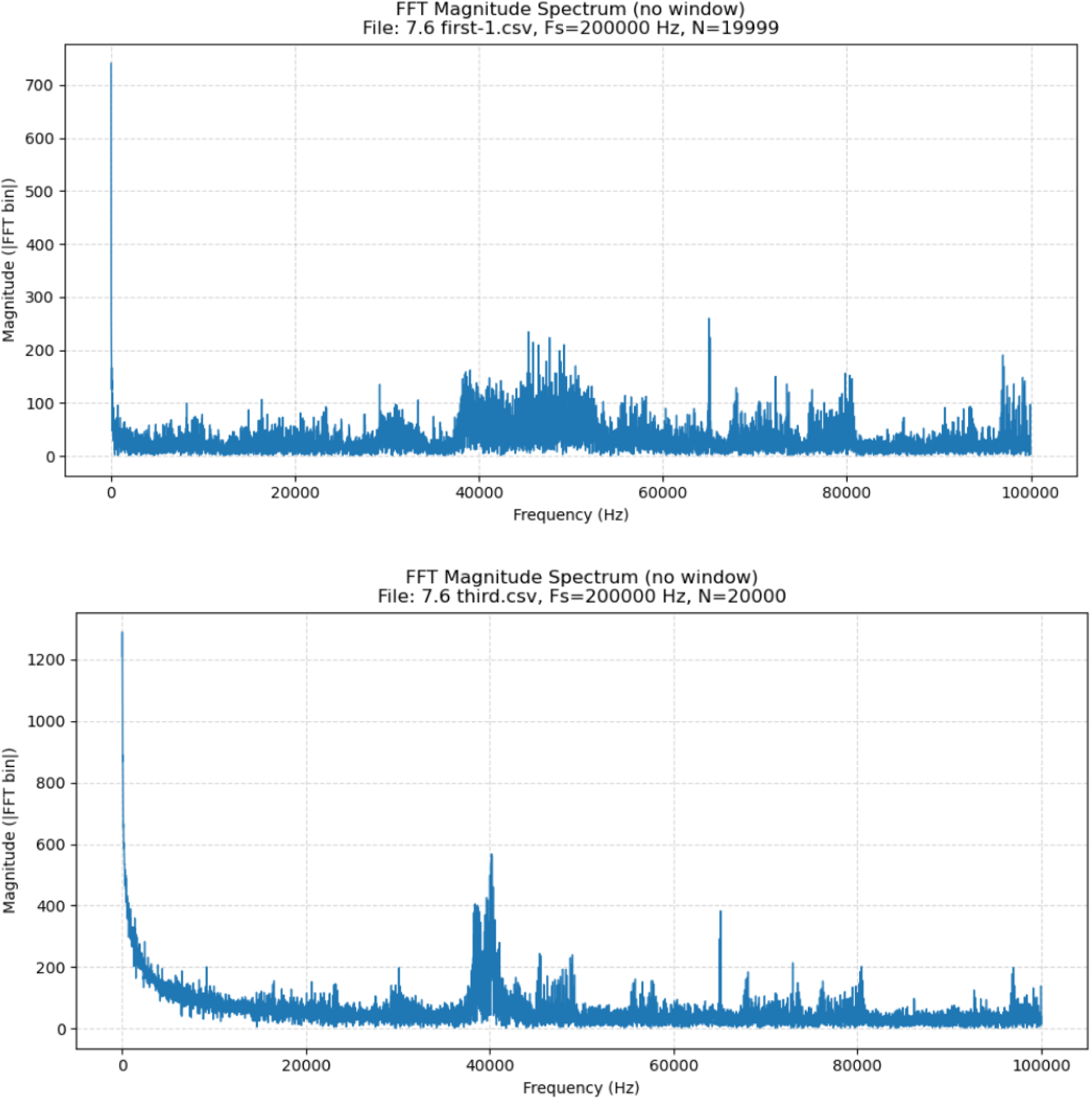

These results are scans that are separated by an intervening EIS scan, and while they differ in appearance, they have a relatively high cosine similarity (similarity) and a relatively low root mean square error (difference).

| Metric | Value |
| --- | --- |
| Cosine Similarity | 0.776353 |
| RMSE | 66.344604 |

### 3. Right hemisphere SNc, depth 7.6 mm, first pulse sequence, first period (0 to 100 ms) <u>versus</u> right hemisphere SNc, depth 7.6 mm, first pulse sequence, fifth period (400 to 500 ms)

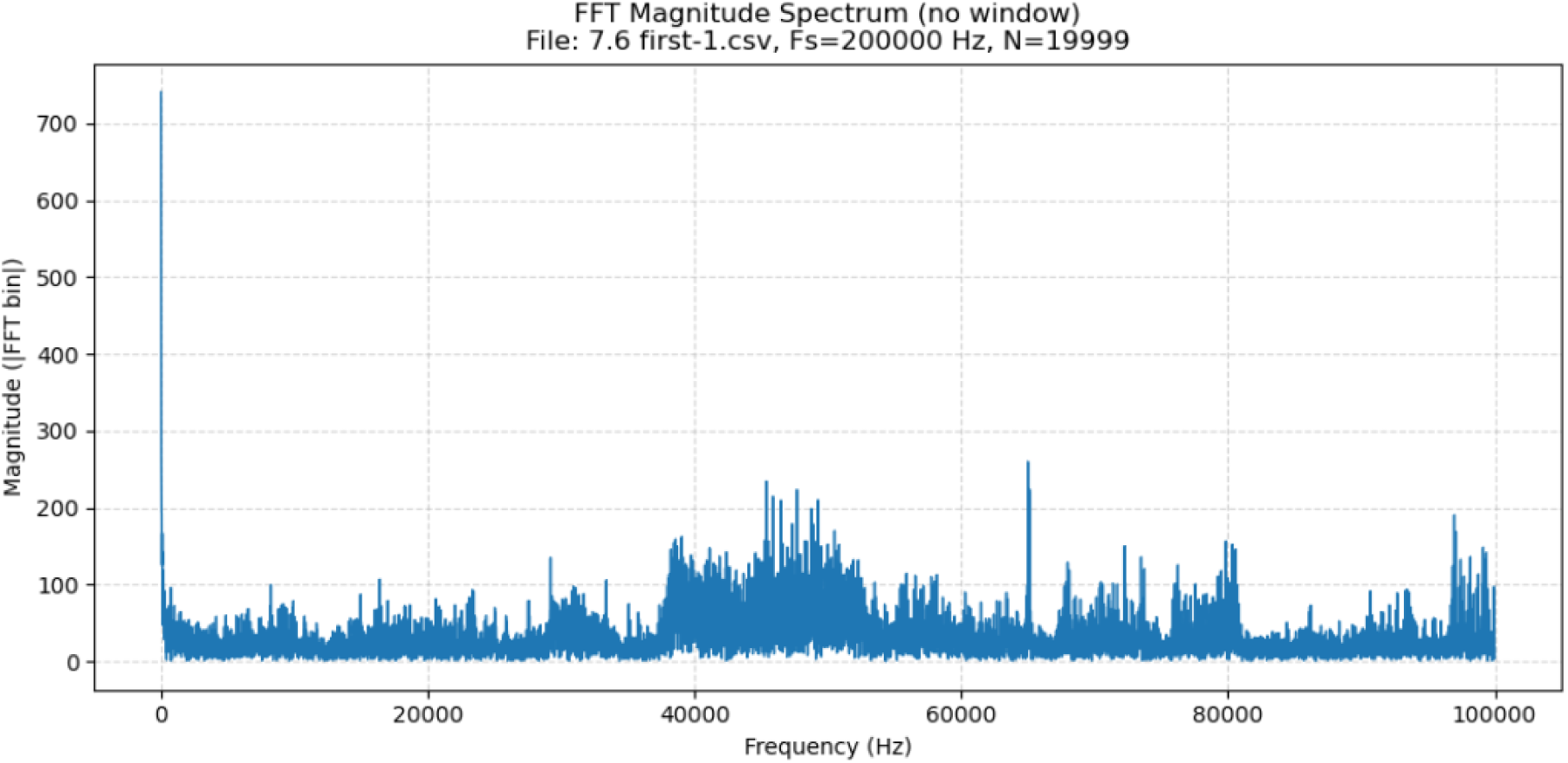
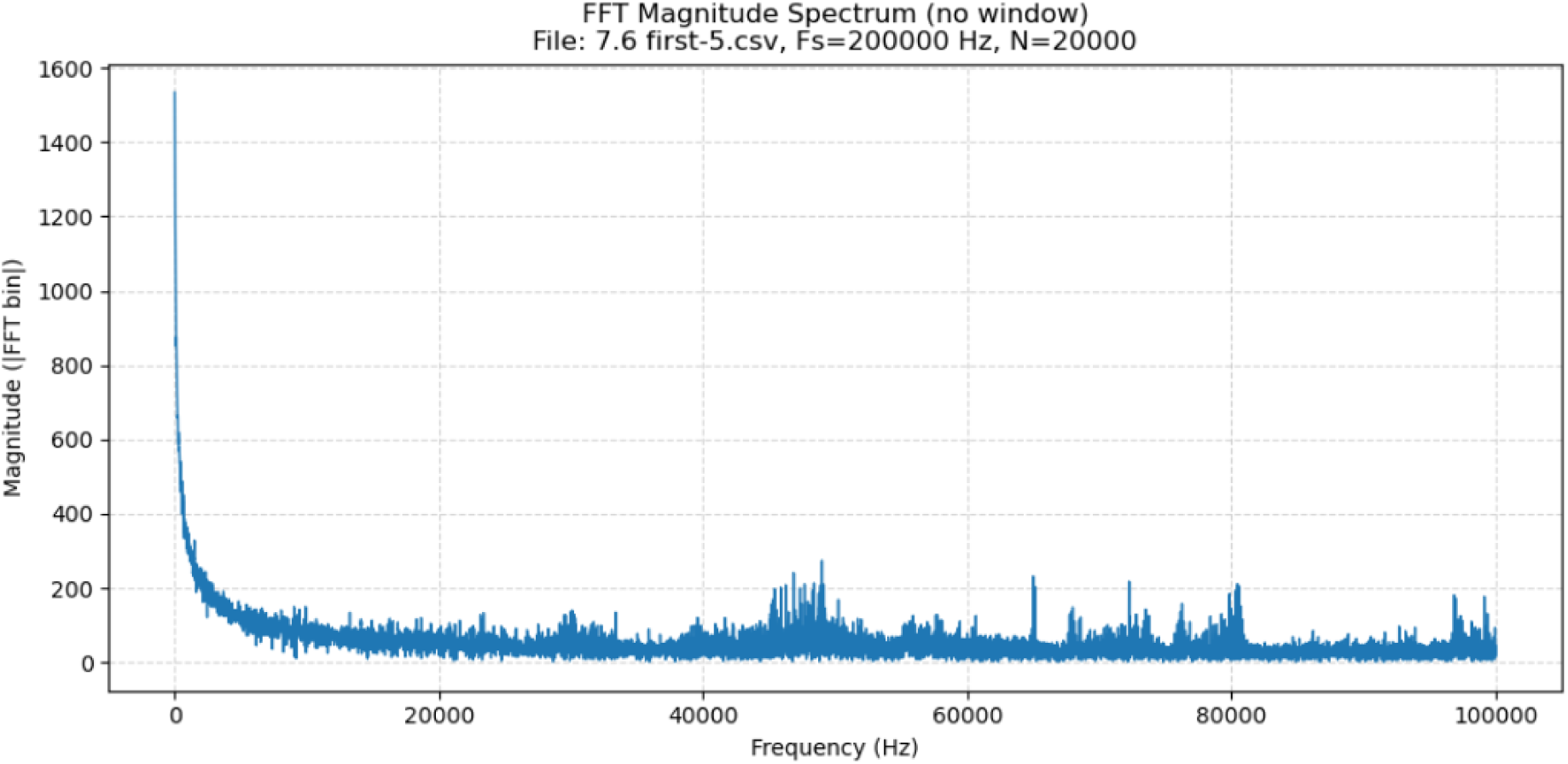

These data show more similarity and more difference between right hemisphere, depth 7.6 mm, first pulse sequence, first period (0 to 100 ms) versus right hemisphere, depth 7.6 mm, first pulse sequence, fifth period (400 to 500 ms) compared to right hemisphere, depth 7.6 mm, first pulse sequence, first period (0 to 100 ms) versus right hemisphere, depth 7.6 mm, third pulse sequence, first period (0 to 100 ms).

| Metric | Value |
| --- | --- |
| Cosine Similarity | 0.780381 |
| RMSE | 75.121526 |

### 4. Right hemisphere SNc, depth 7.6 mm, first pulse sequence, first period (0 to 100 ms) <u>versus</u> right hemisphere SNc, depth 7.6 mm, first pulse sequence, second period (100 to 200 ms)

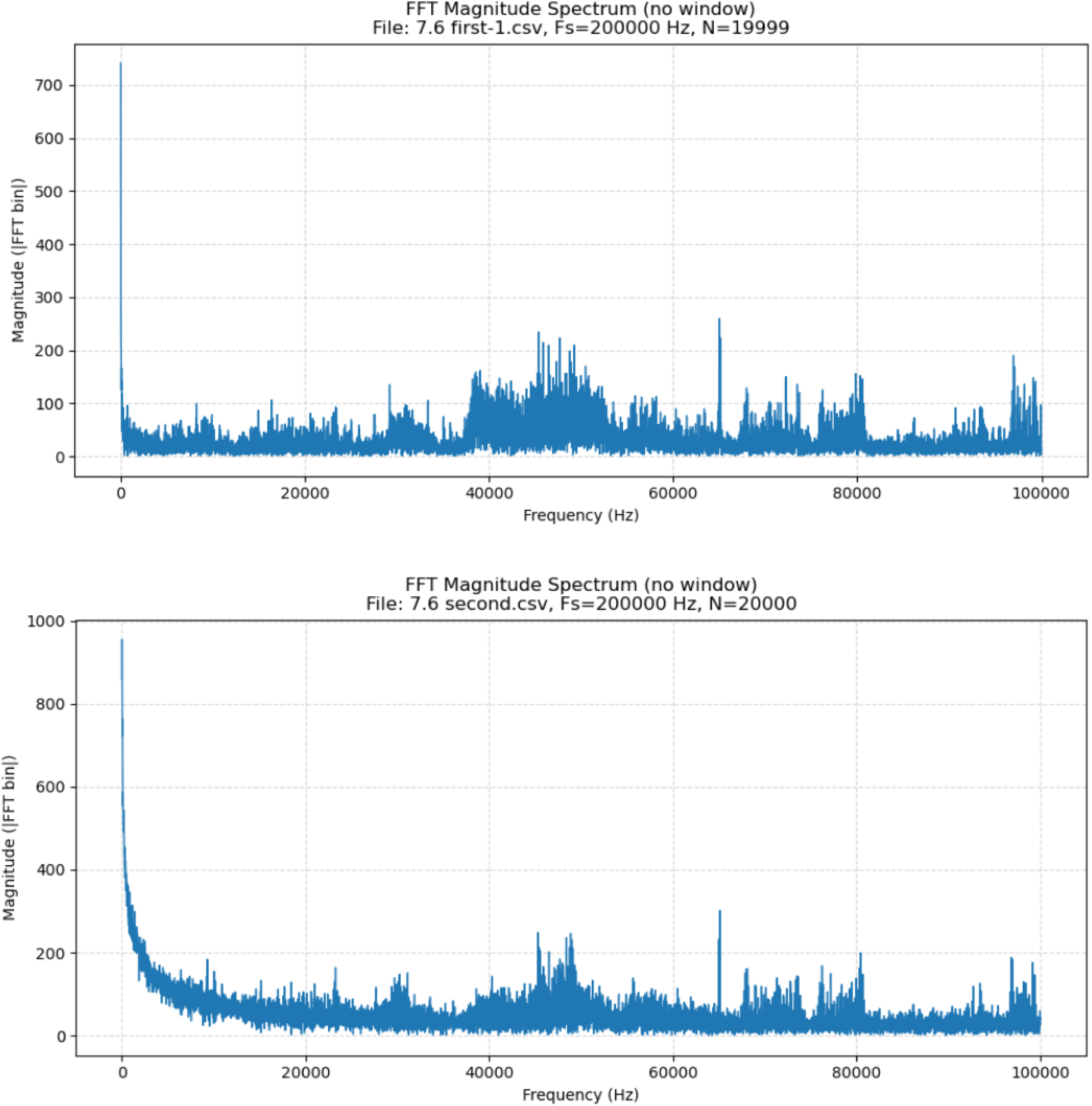

These metrics show more similarity and less difference between right hemisphere, depth 7.6 mm, first pulse sequence, first period (0 to 100 ms) versus right hemisphere, depth 7.6 mm, first pulse sequence, second period (100 to 200 ms) compared to right hemisphere, depth 7.6 mm, first pulse sequence, first period (0 to 100 ms) versus either of right hemisphere, depth 7.6 mm, third pulse sequence, first period (0 to 100 ms) or Right hemisphere, depth 7.6 mm, first pulse sequence, first period (0 to 100 ms) versus right hemisphere, depth 7.6 mm, first pulse sequence, fifth period (400 to 500 ms).

| Metric | Value |
| --- | --- |
| Cosine Similarity | 0.832 |
| RMSE | 60.53 |

### 5. Right hemisphere SNc, depth 7.6 mm, first pulse sequence, first period (0 to 100 ms) <u>versus</u> right hemisphere SNc, depth 7.6 mm, first pulse sequence, third period (200 to 300 ms)

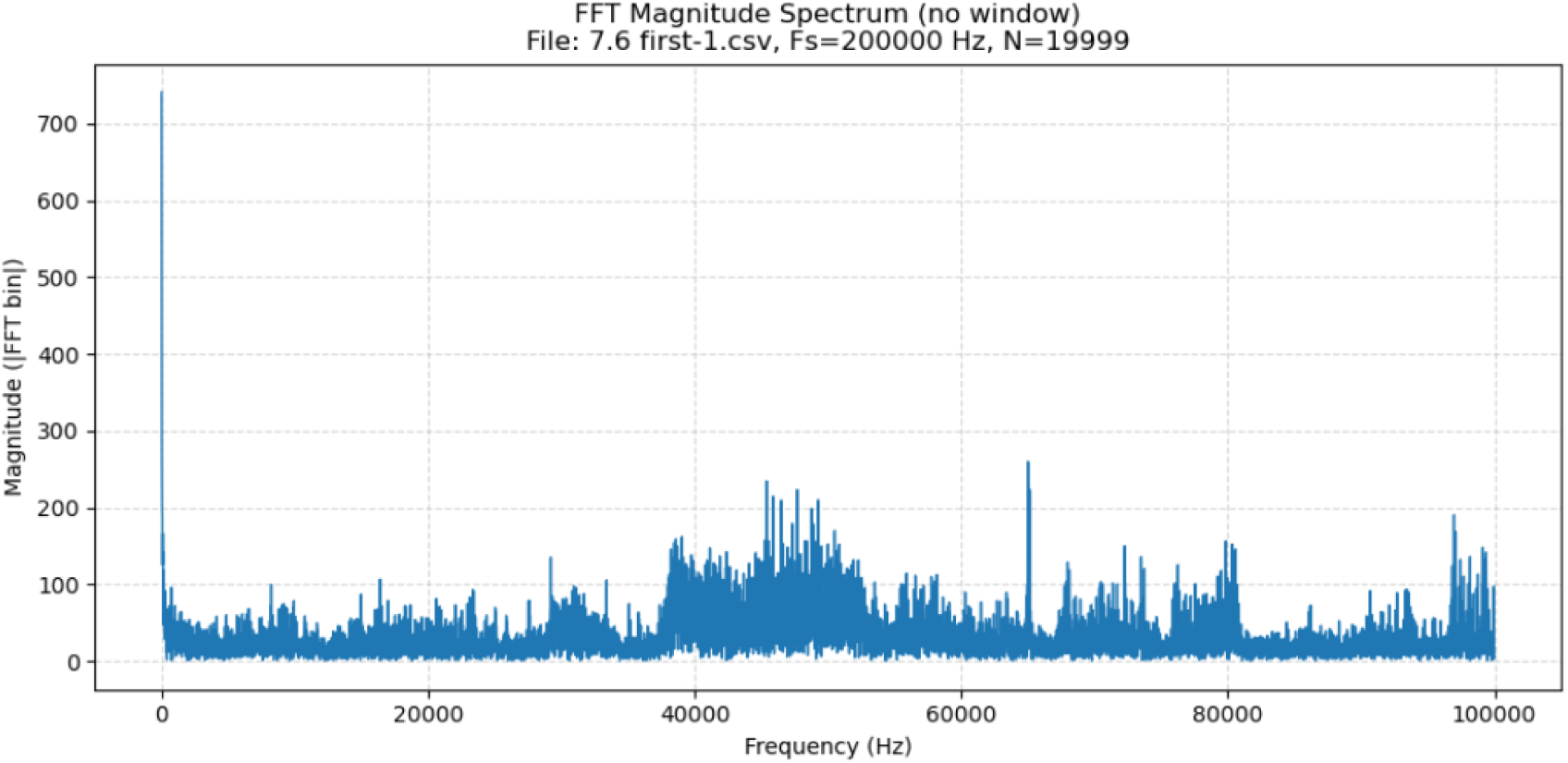
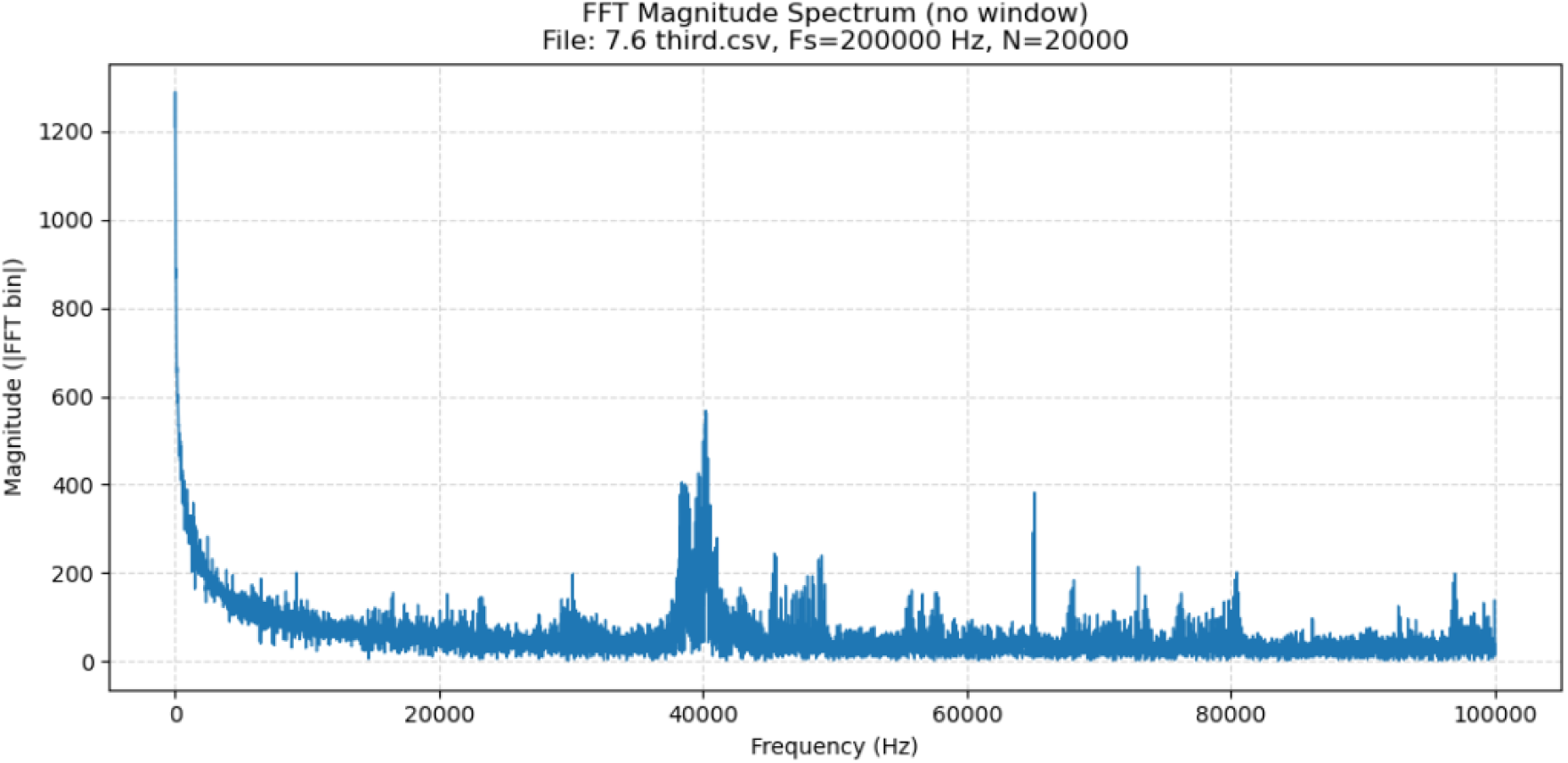

These greater similarity and greater difference between right hemisphere, depth 7.6 mm, first pulse sequence, first period (0 to 100 ms) versus right hemisphere, depth 7.6 mm, first pulse sequence, third period (200 to 300 ms) compared to right hemisphere, depth 7.6 mm, first pulse sequence, first period (0 to 100 ms) versus either of right hemisphere, depth 7.6 mm, third pulse sequence, first period (0 to 100 ms) or Right hemisphere, depth 7.6 mm, first pulse sequence, first period (0 to 100 ms) versus right hemisphere, depth 7.6 mm, first pulse sequence, fifth period (400 to 500 ms).

| Metric | Value |
| --- | --- |
| Cosine Similarity | 0.800856 |
| RMSE | 70.949456 |

### 6. Right hemisphere SNc, depth 7.6 mm, first pulse sequence, first period (0 to 100 ms) <u>versus</u> right hemisphere SNc, depth 7.6 mm, first pulse sequence, fourth period (300 to 400 ms)

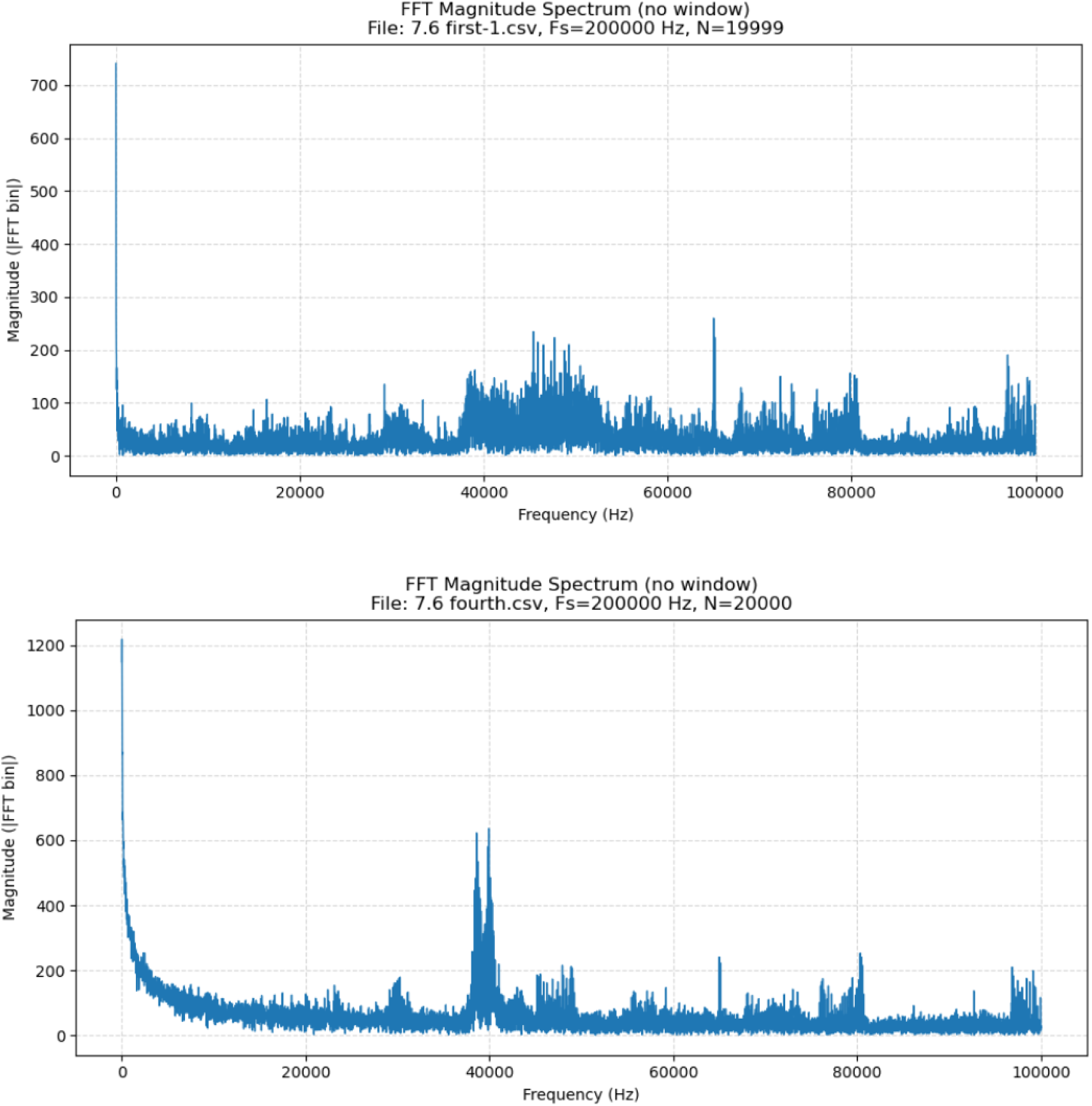

| Metric | Value |
| --- | --- |
| Cosine Similarity | 0.703225 |
| RMSE | 99.950157 |

### 7. Right hemisphere SNc, depth 7.6 mm, first pulse sequence, fourth period (300 to 400 ms) <u>versus</u> right hemisphere SNc, depth 7.6 mm, first pulse sequence, fifth period (400 to 500 ms)

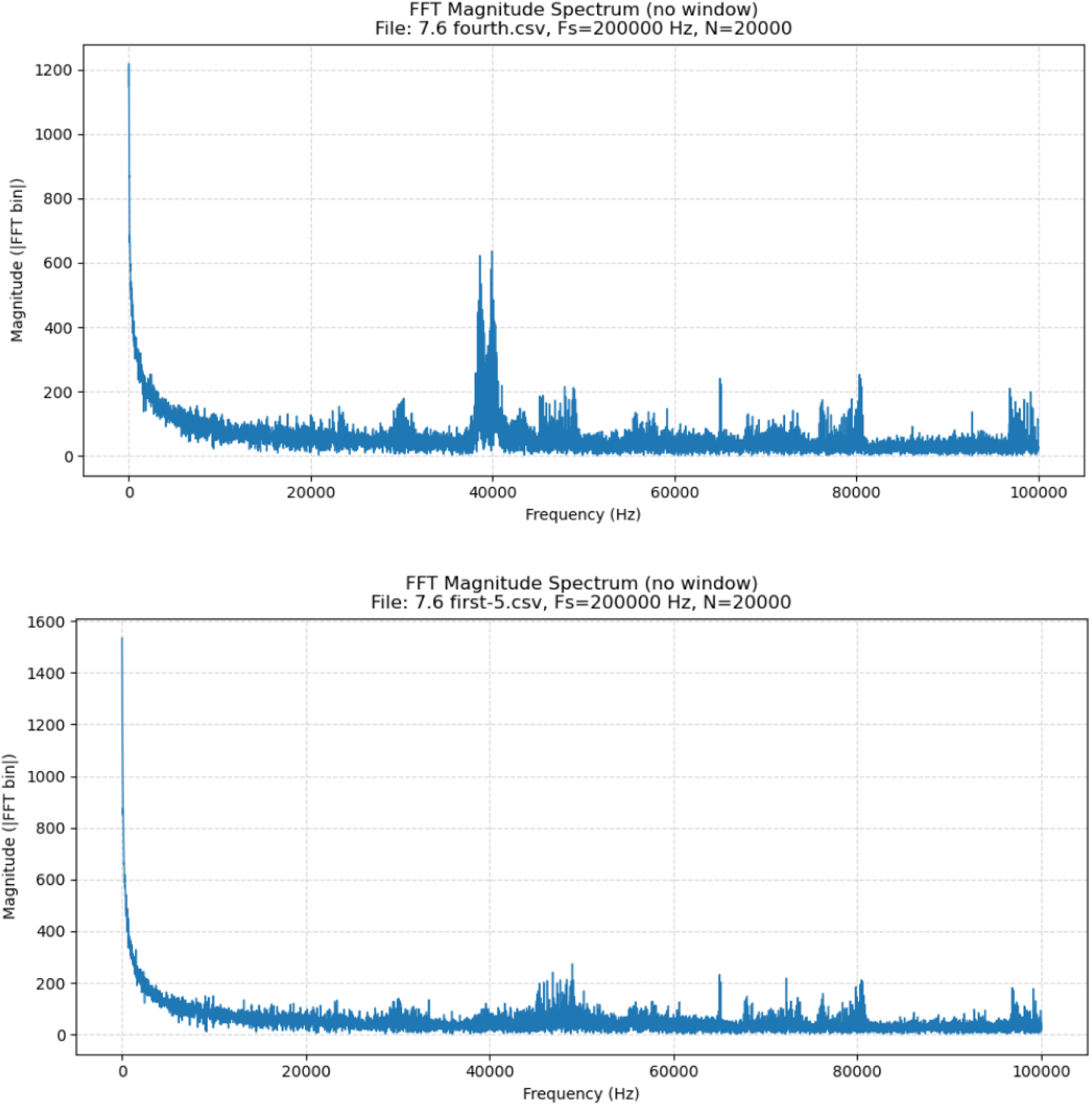

| Metric | Value |
| --- | --- |
| Cosine Similarity | 0.873896 |
| RMSE | 68.385782 |

### 8. Right hemisphere cortex, depth 5 mm, first pulse sequence, second period (100 to 200 ms) <u>versus</u> right hemisphere SNc, depth 7.6 mm, first pulse sequence, first period (0 to 100 ms) and second period (100 to 200 ms)

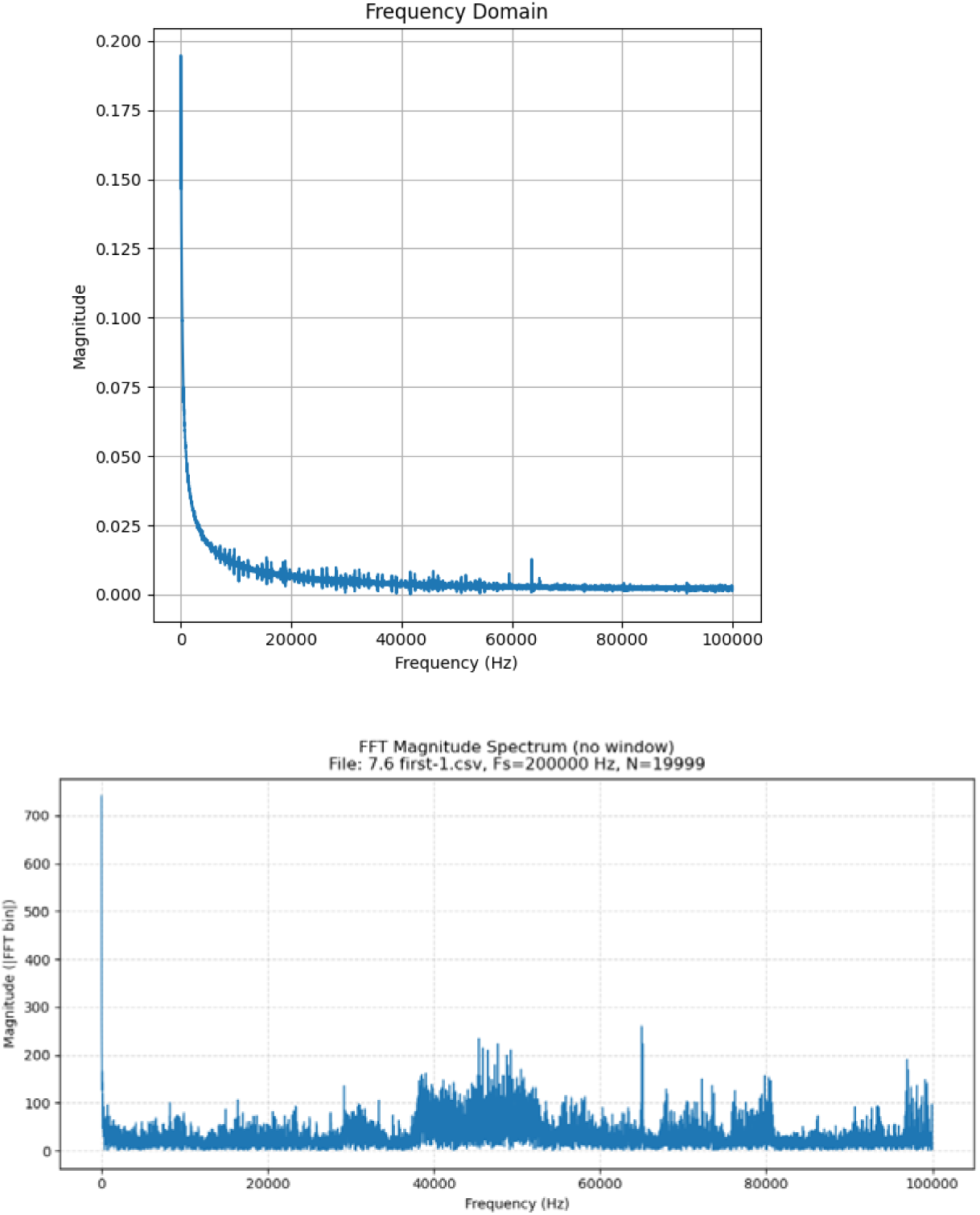
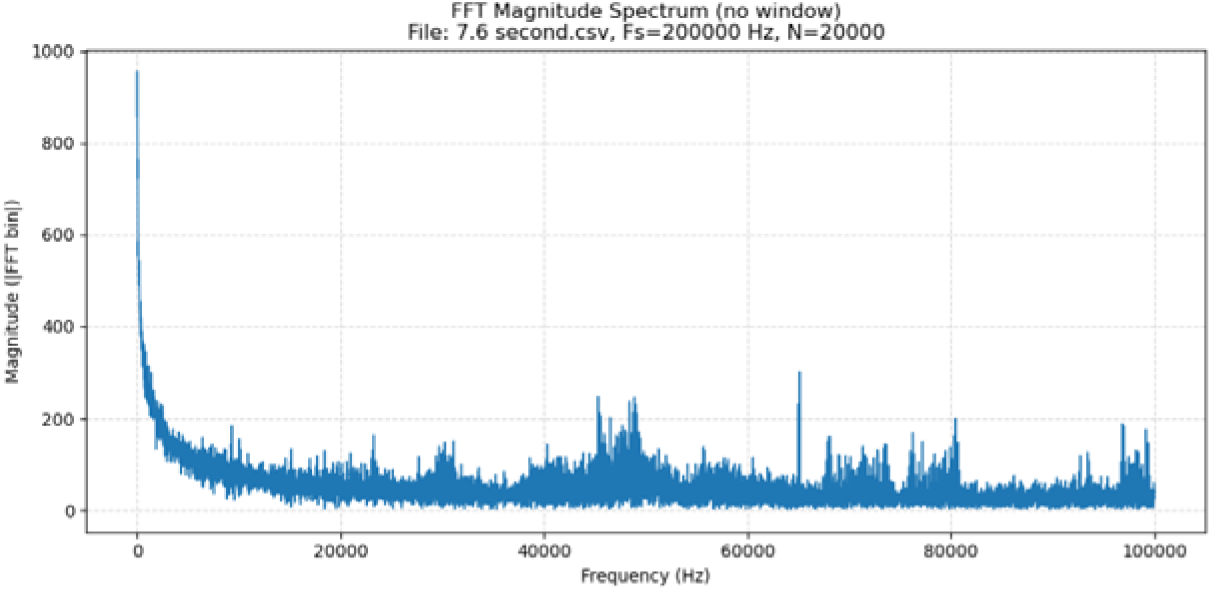

| Comparison | Cosine Similarity | RMSE |
| --- | --- | --- |
| r-5 vs 7.6 first 1 | **0.3507** | **111.88** |
| r-5 vs 7.6 first 2 | **0.5594** | **98.08** |

### 9. Left hemisphere, cortex depth 5 mm, first pulse sequence, second period (100 to 200 ms) <u>versus</u> right hemisphere, cortex depth 5 mm, first pulse sequence, second period (100 to 200 ms)

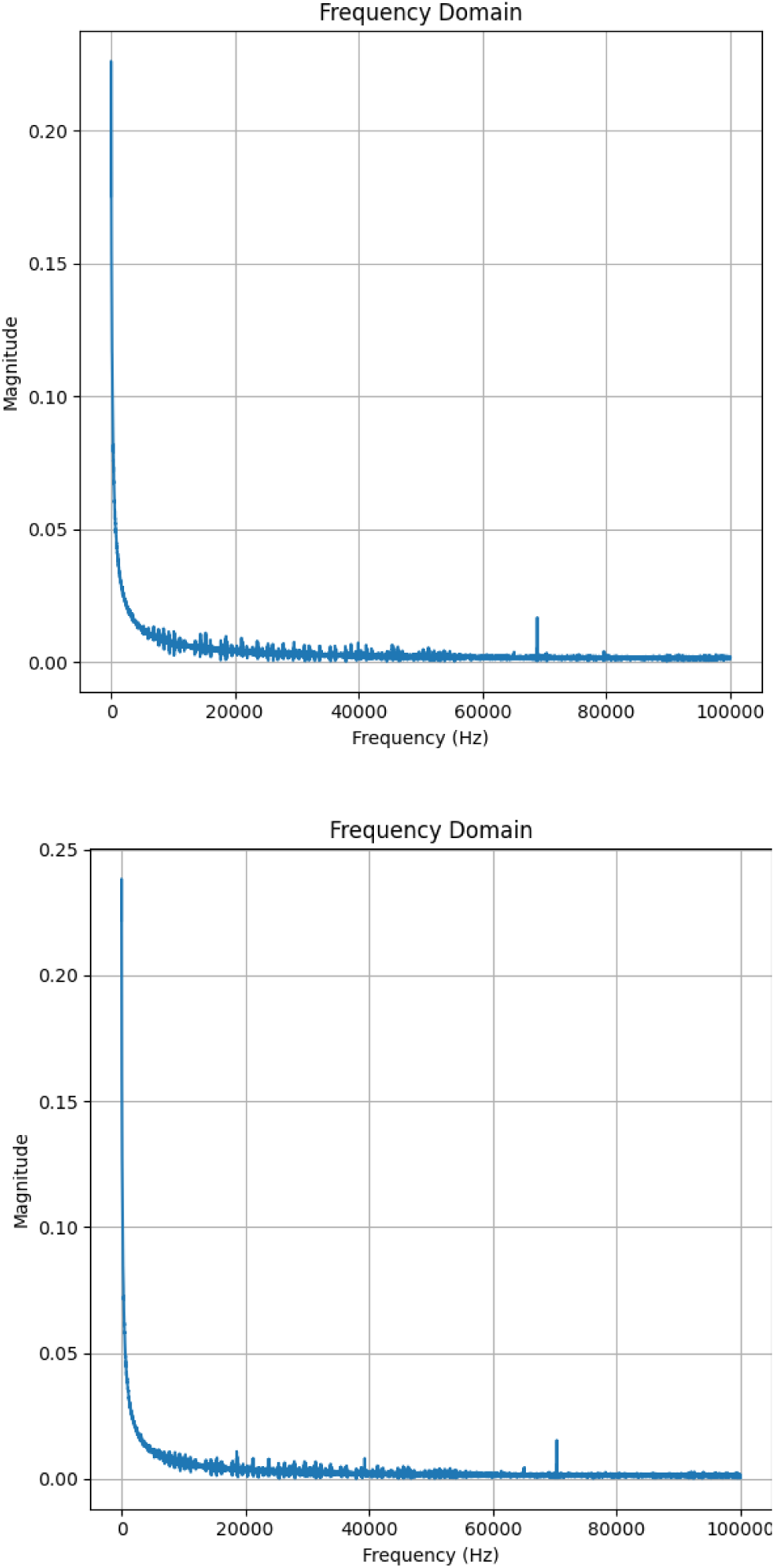

| Metric | Value |
| --- | --- |
| Cosine Similarity | 0.992979 |
| RMSE | 14.201485 |

### 10. Right hemisphere, zona incerta depth 7.6 mm, first pulse sequence, second period (100 to 200 ms) <u>versus</u> right hemisphere, SNc depth 7.6 mm, first pulse sequence, second period (100 to 200 ms)

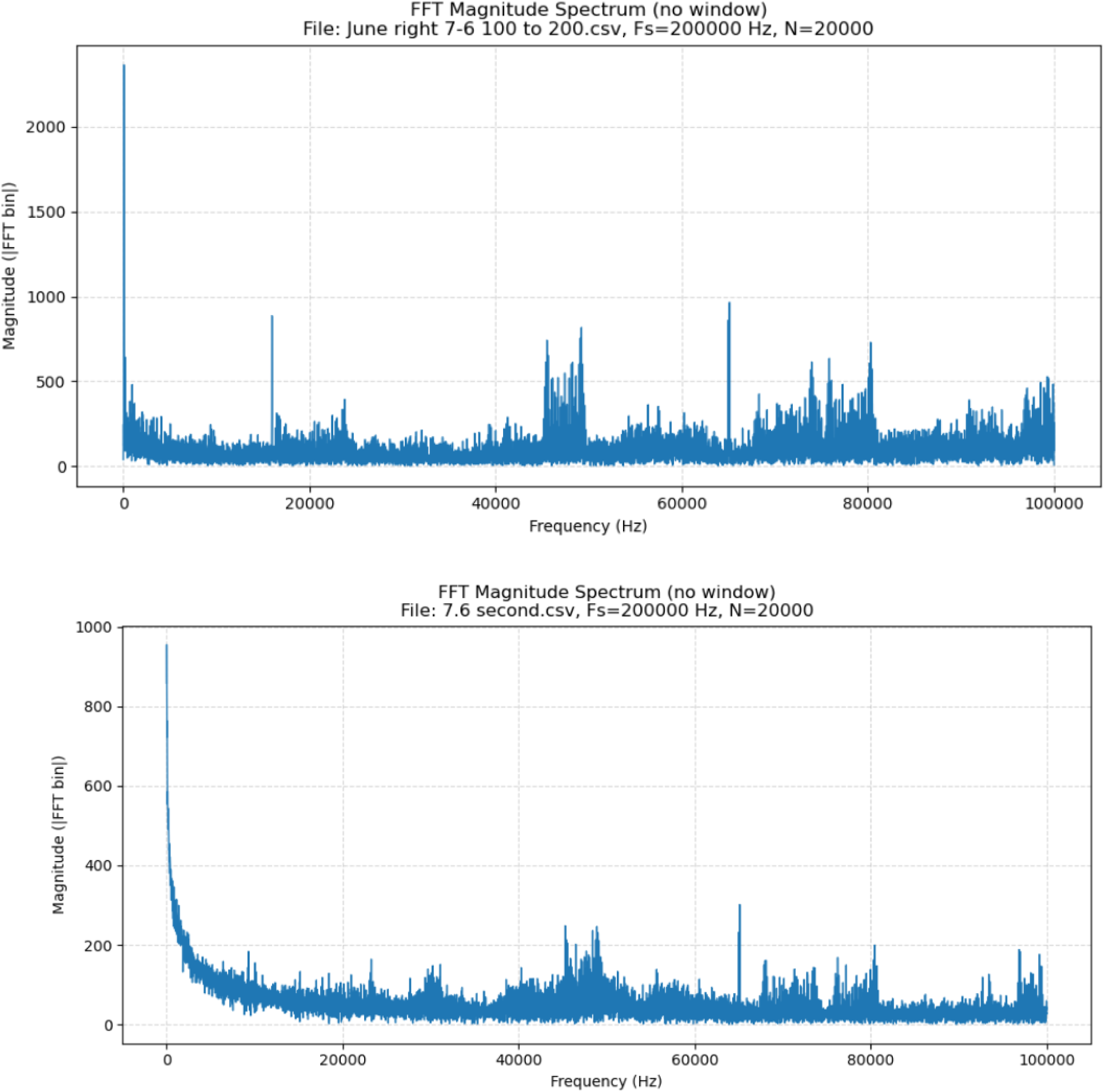

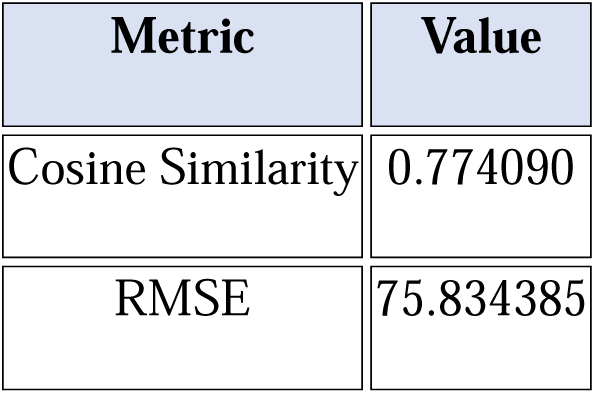

### 11. Right hemisphere, SNc depth 7.6 mm, first pulse sequence, second period (100 to 200 ms) <u>versus</u> left hemisphere, SNc depth 7.6 mm, first pulse sequence, second period (100 to 200 ms)

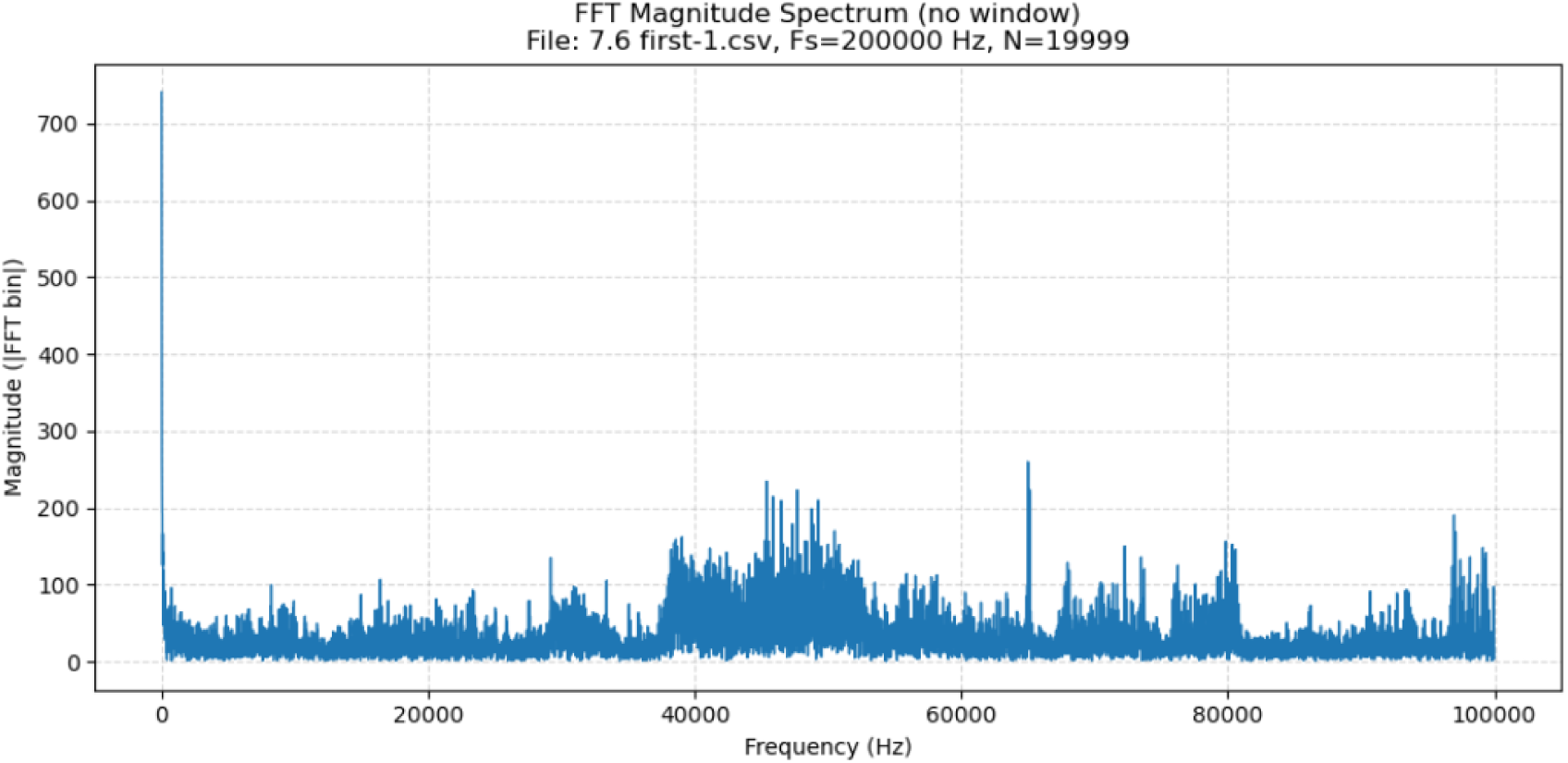
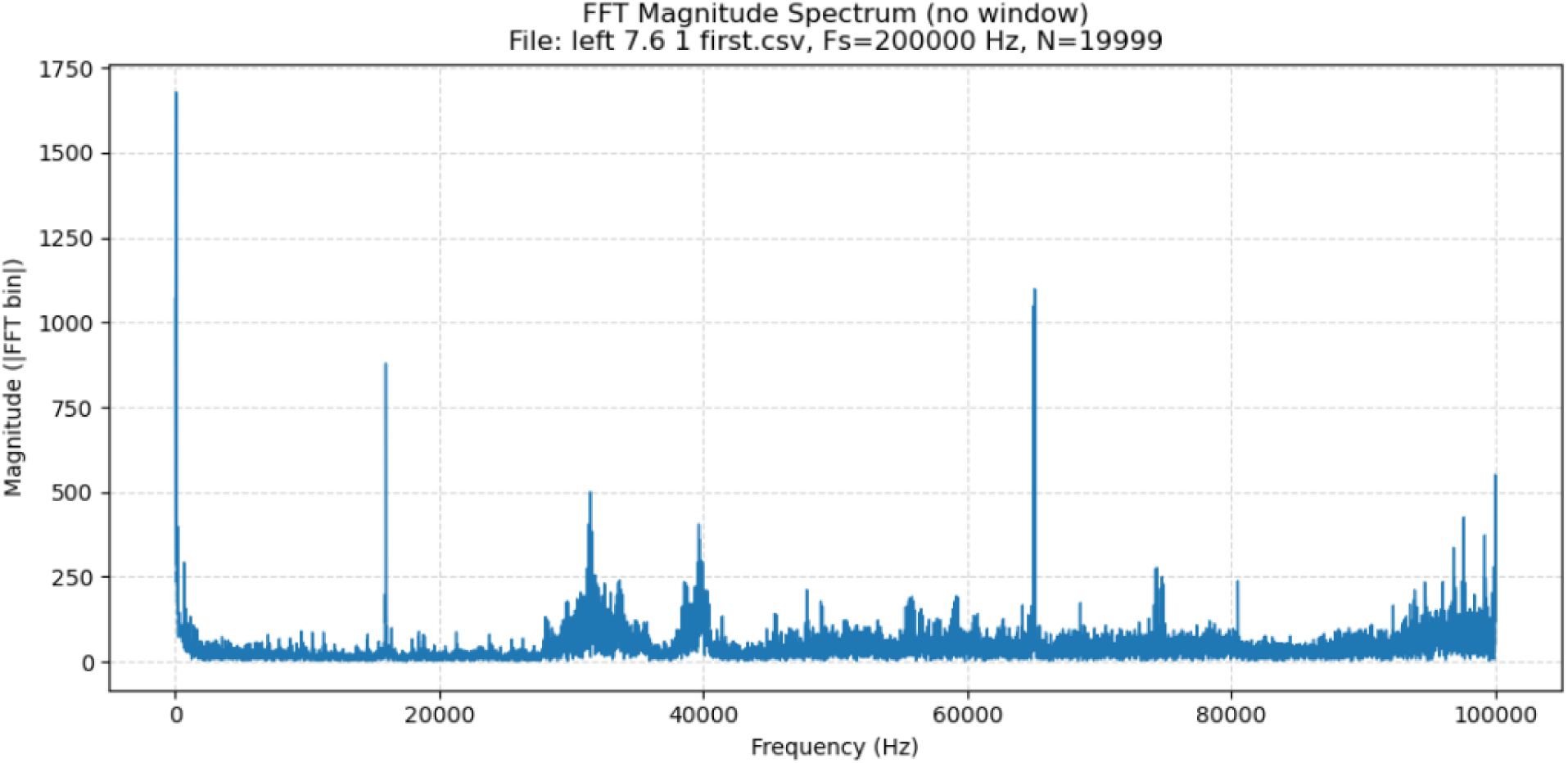

| Metric | Value |
| --- | --- |
| Cosine Similarity | 0.622843 |
| RMSE | 75.912104 |

### 12. Right hemisphere, SNc depth 7.6 mm, first pulse sequence, first period (0 to 100 ms) <u>versus</u> right lateral hypothalamus, depth 8.2 mm, first pulse sequence, first period (100 to 200 ms)

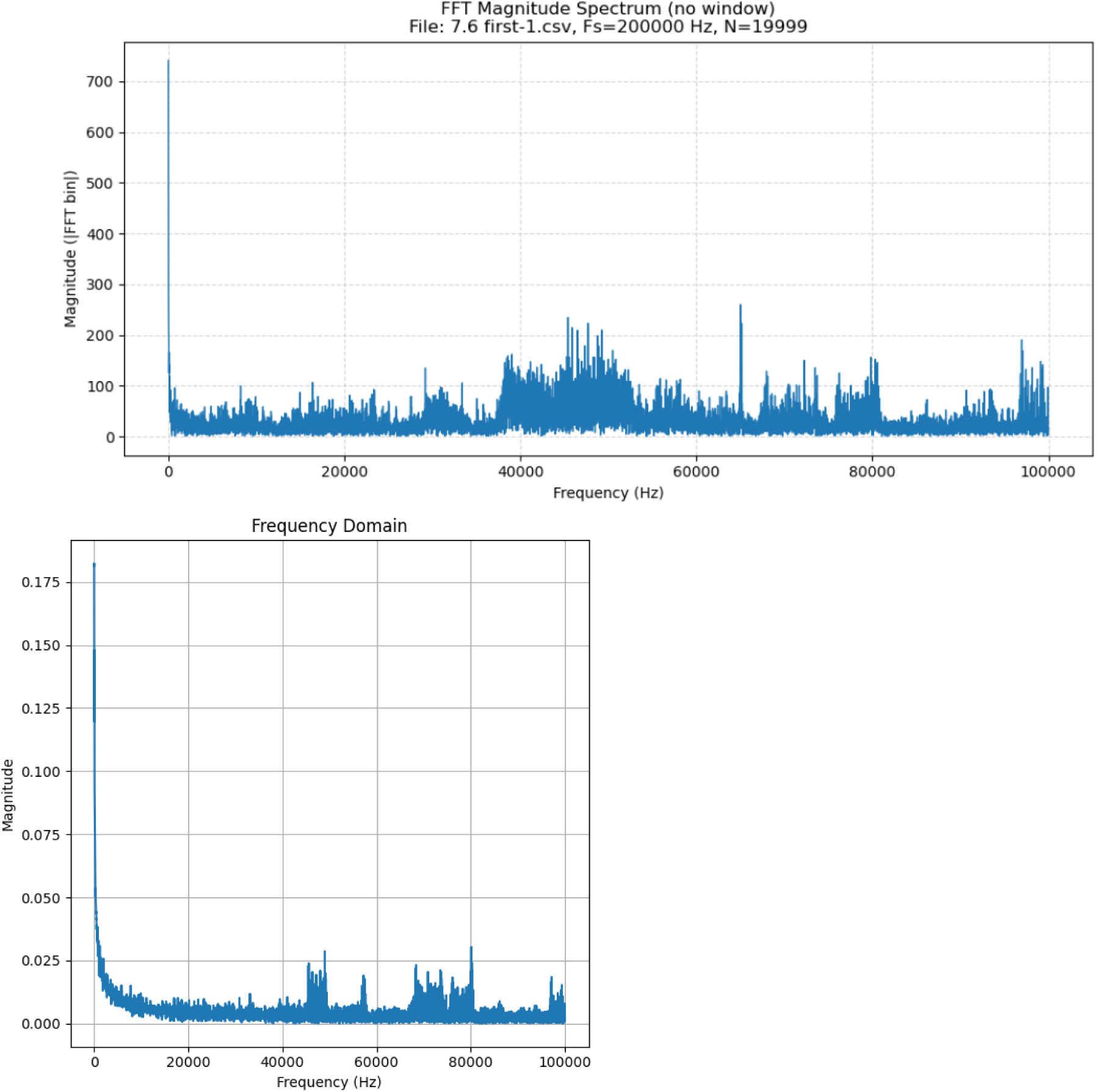

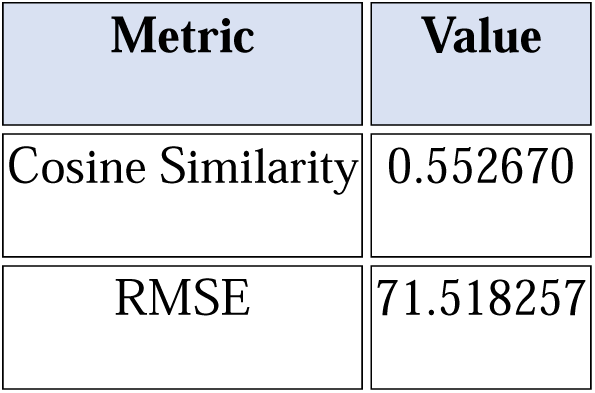

### 13. Right hemisphere, lateral hypothalamus depth 8.2 mm, first pulse sequence, second period (100 to 200 ms) <u>versus</u> cortex right hemisphere, depth 5 mm, first pulse sequence, second period (100 to 200 ms)

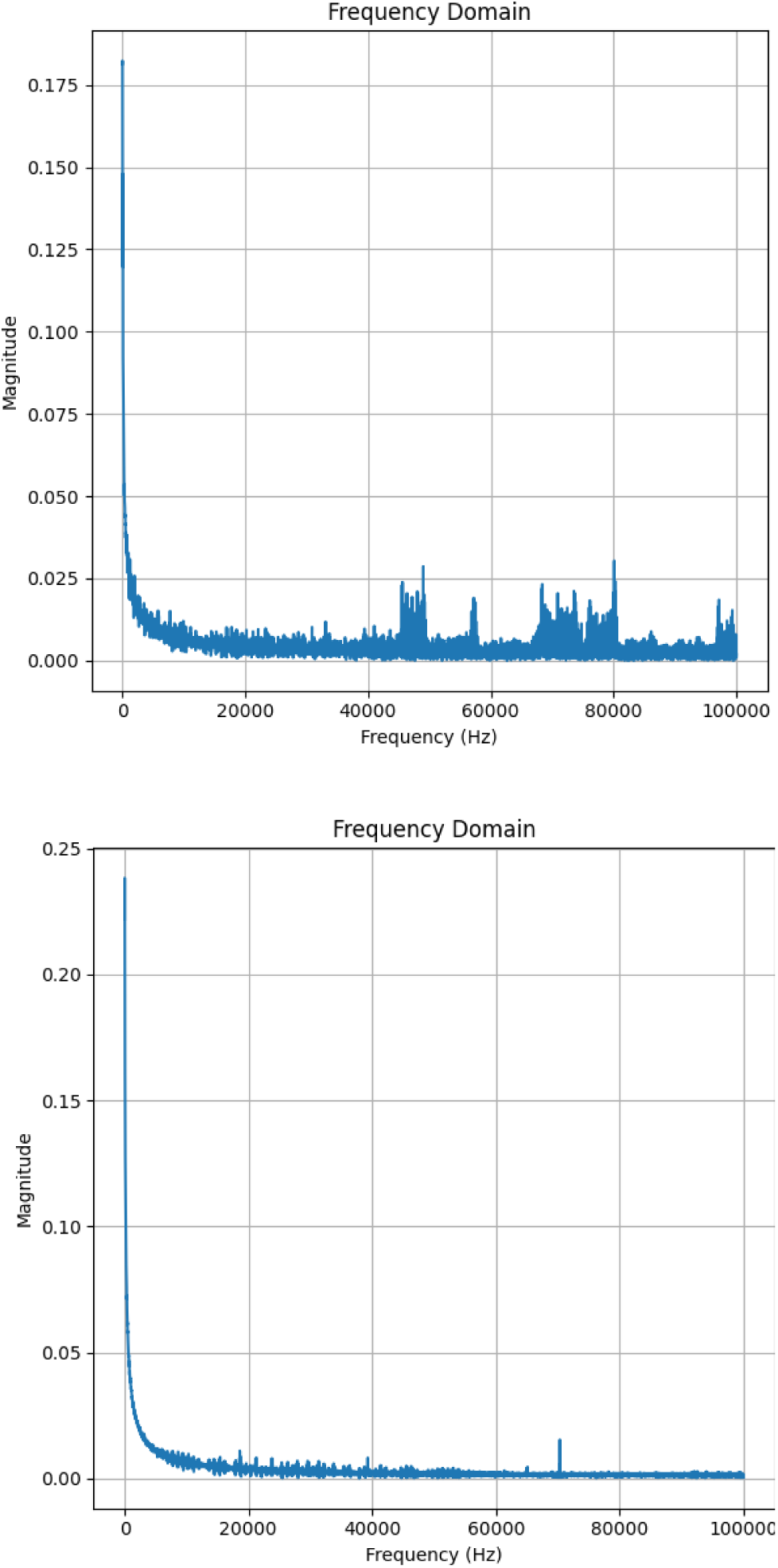

| Metric | Value |
| --- | --- |
| Cosine Similarity | 0.942145 |
| RMSE | 35.045408 |

### 14. Noise signal <u>versus</u> right hemisphere cortex, depth 5 mm, first pulse sequence

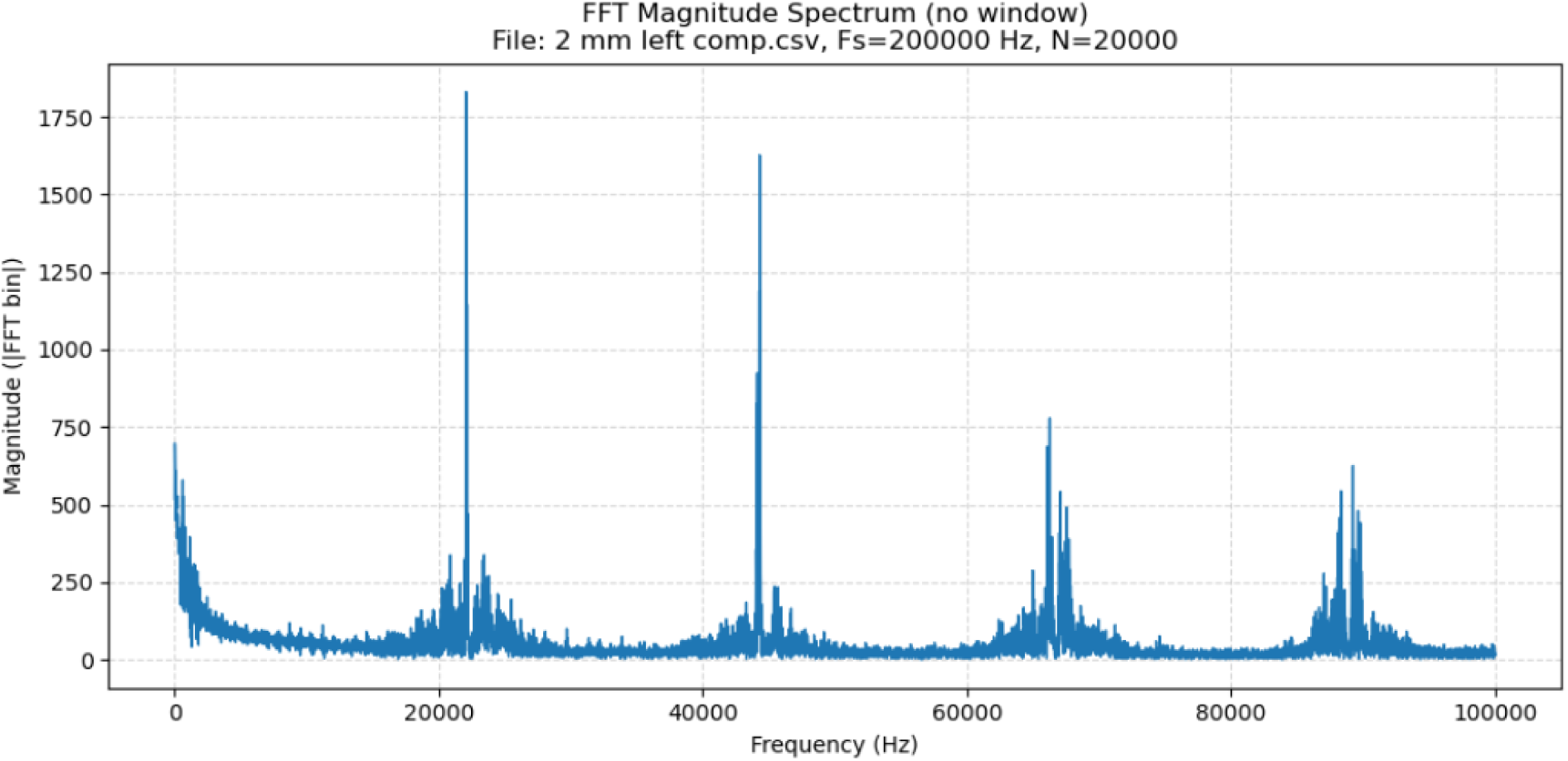
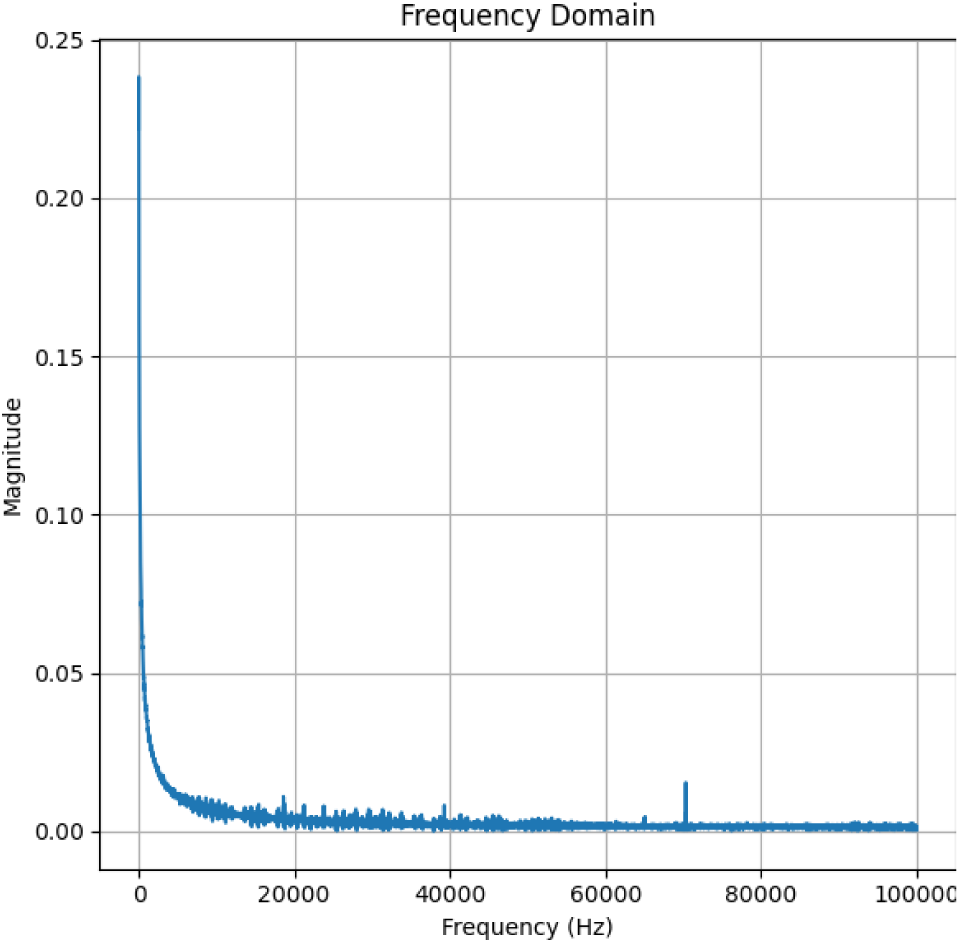

| Metric | Value |
| --- | --- |
| Cosine Similarity | 0.624995 |
| RMSE | 81.802386 |

### 15. Right hemisphere lateral hypothalamus, depth 8.2 mm, first pulse sequence, first period (0 to 100 ms) <u>versus</u> right hemisphere, depth 7.6 mm, first pulse sequence, first period (0 to 100 ms)

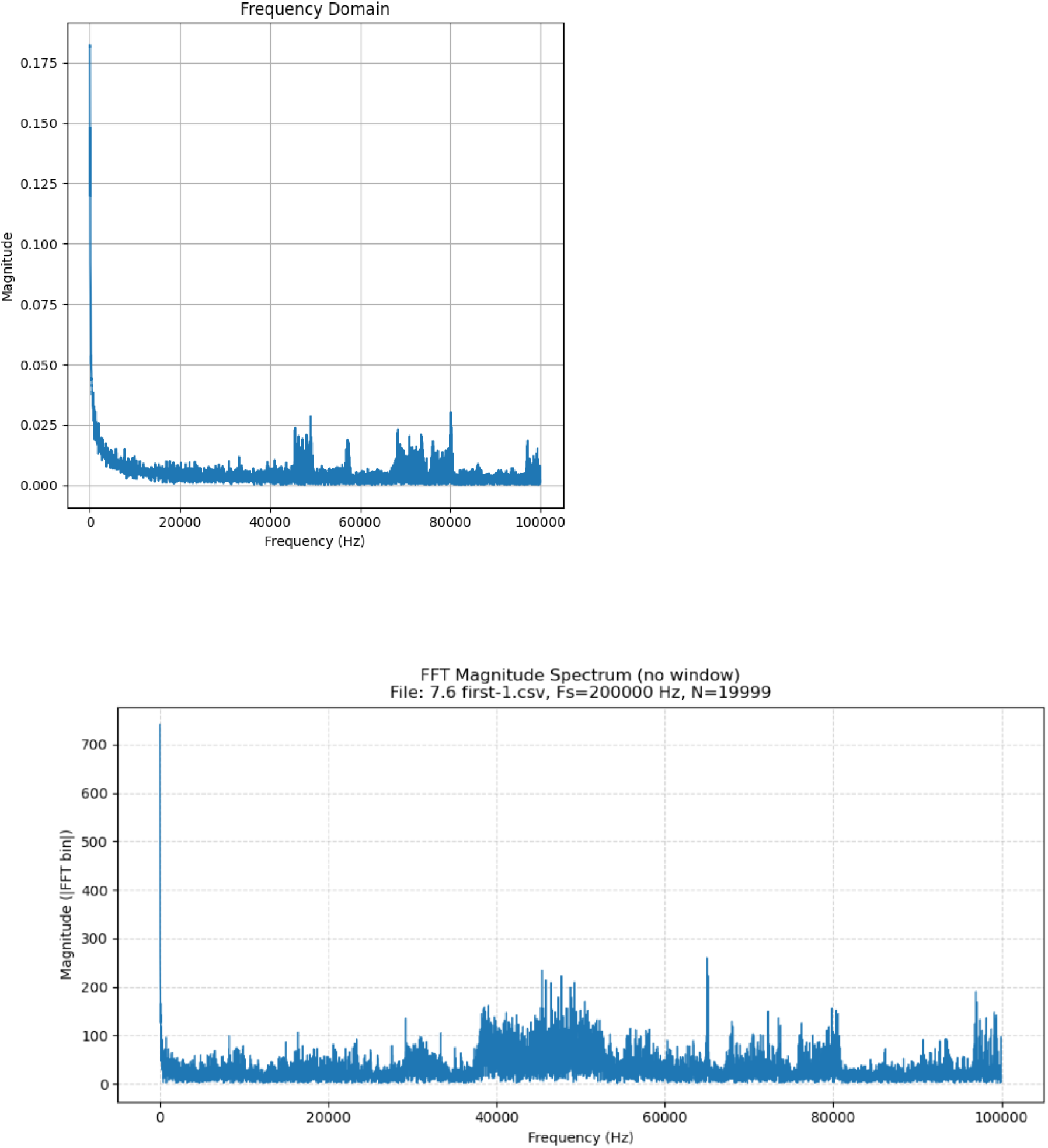

| Metric | Value |
| --- | --- |
| Cosine Similarity | 0.552670 |
| RMSE | 71.518257 |

### 16. Noise signal <u>versus</u> right hemisphere SNc, depth 7.6 mm, first pulse sequence, first period (0 to 100 ms)

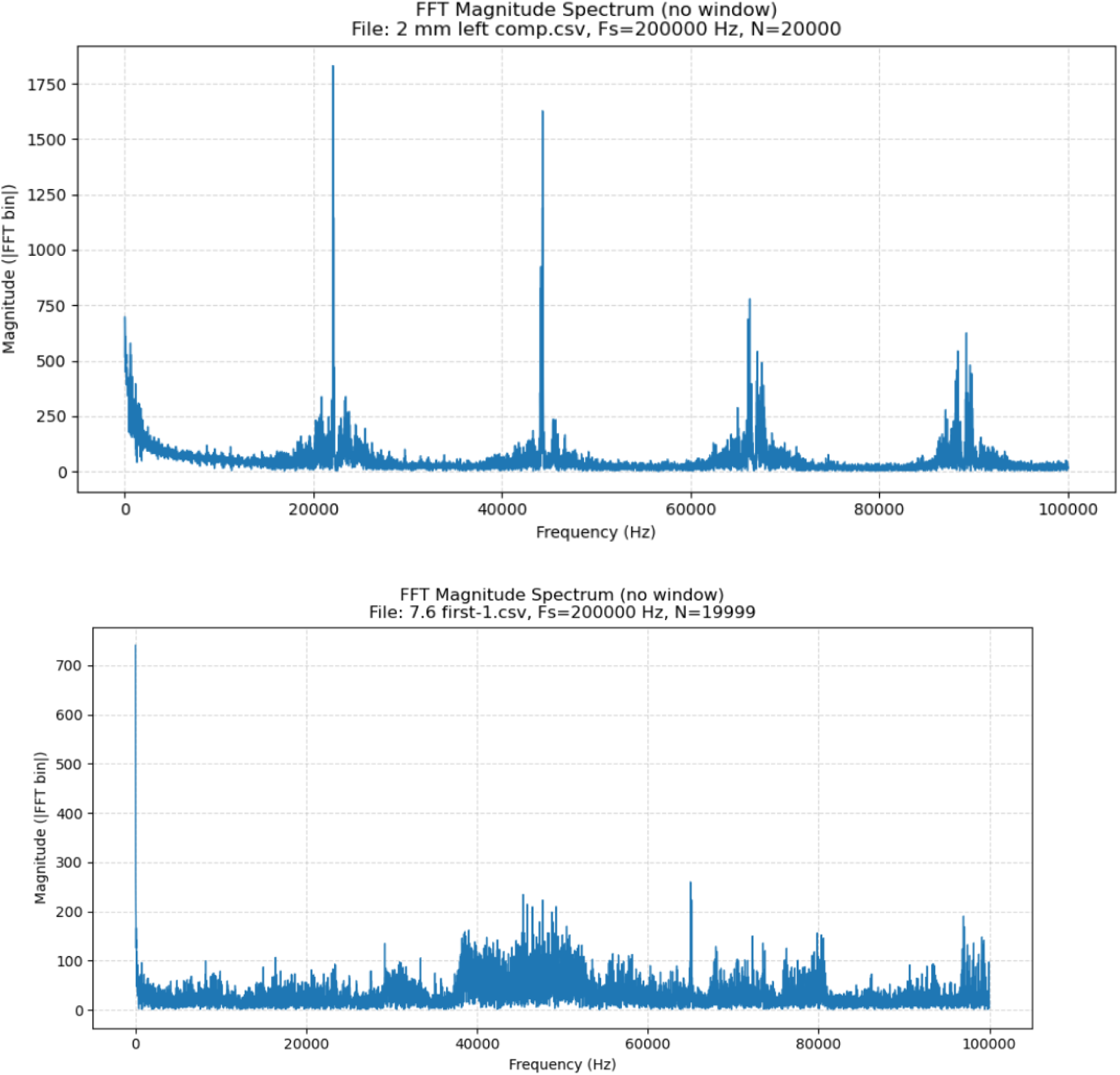

| Metric | Value |
| --- | --- |
| Cosine Similarity | 0.426005 |
| RMSE | 84.363581 |

